# AMPK targets PDZD8 to trigger carbon source shift to glutamine

**DOI:** 10.1101/2023.07.20.548338

**Authors:** Mengqi Li, Yu Wang, Xiaoyan Wei, Wei-Feng Cai, Mingxia Zhu, Luming Yao, Yongliang Wang, Yan-Hui Liu, Jianfeng Wu, Jinye Xiong, Xiao Tian, Qi Qu, Renxiang Xie, Xiaomin Li, Siwei Chen, Xi Huang, Cixiong Zhang, Changchuan Xie, Yaying Wu, Zheni Xu, Baoding Zhang, Bin Jiang, Yong Yu, Zhi-Chao Wang, Qinxi Li, Gang Li, Shu-Yong Lin, Li Yu, Hai-Long Piao, Xianming Deng, Chen-Song Zhang, Sheng-Cai Lin

## Abstract

The shift of carbon utilisation from glucose to other nutrients is a fundamental metabolic adaptation to cope with the decreased glucose oxidation during fasting or starvation^1^. AMP-activated protein kinase (AMPK) plays crucial roles in manifesting physiological benefits accompanying glucose starvation or calorie restriction^2^. However, the underlying mechanisms are unclear. Here, we show that low glucose-induced activation of AMPK plays a decisive role in the shift of carbon utilisation from glucose to glutamine. We demonstrate that endoplasmic reticulum (ER)-localised PDZD8, which we identify to be a new substrate of AMPK, is required for the glucose starvation-promoted glutaminolysis. AMPK phosphorylates PDZD8 at threonine 527 (T527), and promotes it to interact with and activate the mitochondrial glutaminase 1 (GLS1), a rate-limiting enzyme of glutaminolysis^3–5^, and as a result the ER-mitochondria contact is strengthened. In vivo, PDZD8 enhances glutaminolysis, and triggers mitohormesis that is required for extension of lifespan and healthspan in *Caenorhabditis elegans* subjected to glucose starvation or caloric restriction. Muscle-specific re-introduction of wildtype PDZD8, but not the AMPK-unphosphorylable PDZD8-T527A mutant, to *PDZD8*^−/−^ mice is able to rescue the increase of glutaminolysis, and the rejuvenating effects of caloric restriction in aged mice, including grip strength and running capacity. Together, these findings reveal an AMPK-PDZD8-GLS1 axis that promotes glutaminolysis and executes the anti-ageing effects of calorie restriction by promoting inter-organelle crosstalk between ER and mitochondria.

---

Fasting causes rapid depletion of stored carbohydrates, monomeric or polymeric, leading to declining blood glucose levels. Nutritional adaptation is hence a fundamental measure to maintain energy balance. In metazoans, there are orchestrated interplays among organs and tissues to produce and redistribute alternative fuels, mainly fatty acids and amino acids^6–12^. Fatty acids, particularly the long-chain fatty acids released from triglycerides, are first converted to fatty acyl-CoA, which is then transported into mitochondria via the carnitine palmitoyltransferase transporters (CPT1 and CPT2)^9,13–15^. Inside mitochondria, the acyl-CoA undergoes β-oxidation to generate acetyl-CoA that enters the tricarboxylic acid (TCA) cycle to produce energy^16–18^. Among amino acids, glutamine is the most abundant circulating amino acid, comprising more than 50% of free amino acid pool in the body during starvation, and serves as a key alternative carbon source^19–22^. It is known that glutamine, along with alanine, is converted from other amino acids, particularly the branched chain amino acids from muscle protein breakdown under starvation^22–24^. While alanine mainly contributes to hepatic gluconeogenesis in the liver, glutamine is utilised in various tissues to directly meet energy demand^20,23–25^, as well as for gluconeogenesis in the liver, intestines, and kidney^26–31^. In addition, glutamine can also act as a major source for GSH and NADPH synthesis during the starvation to maintain the cellular redox state^32–36^.

AMPK plays a central role in maintaining energy homeostasis, mainly through phosphorylating multiple targets to stimulate catabolism and inhibit anabolism, thereby promoting ATP production and reducing ATP consumption^2^. In addition to its classic role as an energy sensor regulated by increased AMP and ADP levels^37,38^, AMPK is highly sensitive to activation by falling levels of glucose under fasting conditions, independent of decrease of cellular energy status^39,40^. In this, it is the declining levels of glycolytic intermediate fructose-1,6-bisphosphate (FBP) that trigger activation of lysosomally localised AMPK by the upstream kinase LKB1 via the glucose-sensing pathway comprising aldolase (direct sensor for the presence or absence of FBP^40^), transient receptor potential V (TRPVs), vacuolar H^+^-ATPase (v-ATPase), Ragulator and AXIN^40–43^. Upon activation by the glucose-sensing axis, AMPK phosphorylates acetyl-CoA carboxylase 1 (ACC1)^44^, which inhibits the production of malonyl-CoA to remove the inhibition of CPT1, thereby promoting the transport of acyl-CoA into mitochondria and fatty acid oxidation (FAO)^45^. AMPK also promotes catabolism of amino acids by inhibiting translation, either through inhibiting the target of rapamycin complex 1 (TORC1)^46,47^, or through promoting the inhibition of the eukaryotic elongation factor 2 (eEF2) by eEF2 kinase (eEF2K)^48^. In addition, AMPK helps release free amino acids from cellular proteins either by promoting autophagy (ref. ^49–51^), or through increasing proteasomal degradation of labile proteins^52^. However, the mechanisms underlying the prioritisation and promotion of the alternative carbon sources remain unclear.

In this study, we set out to delineate the molecular events on the path to extension of lifespan following glucose starvation. First, we made an observation that glucose starvation induces an increase of mitochondria-associated membrane (MAM). Through proteomic analysis of the proteins pulled down from MAM by using an antibody against pan-AMPK phosphoproteins, we identified that PDZD8, an ER-localised protein, is a new substrate of AMPK. We show that AMPK-mediated phosphorylation of PDZD8 is required for the increase of glutaminolysis to compensate for the scarcity of glucose before the promotion of FAO. We demonstrate that phosphorylated PDZD8 interacts with and activates GLS1 to enhance glutaminolysis. Most surprisingly, we demonstrate that the enhanced glutaminolysis induces mitohormesis, which is a necessary process for the extension of lifespan and health-span in both mice and nematodes. In short, we have elucidated the molecular mechanism underlying the carbon source shift from glucose to glutamine, and have demonstrated that glutaminolysis is a crucial step on the path to longevity, as a benefit of calorie restriction.

## PDZD8 is a new substrate for AMPK

We were intrigued by the lower yields of pure mitochondria from glucose-starved mouse embryonic fibroblasts (MEFs) after subcellular fractionation^53^, and explored the reasons behind this phenomenon. It turned out that the reduction of the recovered mitochondria was caused by increased association of mitochondria with ER (mitochondria-associated ER membrane, or MAM) (Fig. 1a). The increase of ER-mitochondria contact was also seen with confocal microscopy and electron microscopy (Fig. 1b-e, Extended Data Fig. 1a-e, and Supplementary Note 1 for details). We then wondered whether AMPK played a role in the increase of ER-mitochondria contact, and found that knockout of *AMPK*α (both *AMPK*α*1* and *AMPK*α*2*, the catalytic subunits of AMPK) blocked the enhancement of ER-mitochondria contact in glucose starved MEFs (Fig. 1a-e, Extended Data Fig. 1a-e). We then enriched AMPK substrates from the purified ER-mitochondria contact (MAM and the MAM-tethered mitochondria) of glucose-starved MEFs, by using an antibody specifically recognising pan-phospho-substrates of AMPK that contains the conserved motif to be phosphorylated by AMPK^54–58^. Through mass spectrometry of the pulldown samples, we identified 12 proteins that were preferentially phosphorylated in glucose-starved cells (listed in Supplementary Table 1), among which PDHA1 is a known AMPK substrate^59^. We next generated expression plasmids for these 12 proteins, and found that 3 of them, i.e. PDZD8 and RMDN3, and PDHA1 (as a positive control), were phosphorylated by AMPK in low glucose (Extended Data Fig. 2a). Through knocking out these 3 individual genes in MEFs, we found that PDZD8, known as a component of mammalian ER-mitochondria encounter structure (ERMES) complex required for maintaining ER-mitochondria and ER-lysosome contacts^60,61^, was required for the promotion of ER-mitochondria contact or MAM formation during glucose starvation (Fig. 1f, g, Extended Data Fig. 2c-f; see knockout validation data in Extended Data Fig. 2b). In comparison, although knockout/knockdown of *RMDN3* led to a decrease of basal ER-mitochondria contact, which is consistent with previous reports^62,63^, glucose starvation can still promote the ER-mitochondria contact, and knockout of *PDHA1* did not affect ER-mitochondria contact regardless of glucose starvation (Extended Data Fig. 2h; see knockout validation data in Extended Data Fig. 2g). We next determined the phosphorylation site(s) of PDZD8 by AMPK. PDZD8 contains 160 predicted sites (according to ref. ^54–58^; see Supplementary Note 2 for the prediction), among which 78 were hit by mass spectrometry (Supplementary Table 1). We individually mutated those 78 sites and the other predicted sites as well, and found that T527 (for human; T521 for mouse), conserved in mammals, is the site of PDZD8 for phosphorylation by AMPK. First of all, p-T527 was hit by the mass spectrometry analysis (see representative spectrogram in Extended Data Fig. 3a); secondly, mutation of T527 to alanine (PDZD8-T527A) rendered it unphosphorylable by AMPK in vitro (Fig. 1h); and thirdly, PDZD8-T527A was also unphosphorylable after re-introduction into *PDZD8*^−/−^ MEFs under glucose starvation (Fig. 1i). We then developed a phospho-specific antibody against p-T527-PDZD8 (see validation data using *PDZD8*^−/−^ MEFs expressing wildtype PDZD8 or PDZD8-T527A in Extended Data Fig. 3b), and found that glucose starvation led to a significant elevation of p-T527 signal in the immunoprecipitants of endogenous PDZD8 (Fig. 1j-l). Moreover, knockout of *AMPK*α, as well as *AXIN* or *LAMTOR1* which are known components of the glucose-sensing-AMPK axis^41,43^, abolished p-T527 signal under glucose starvation (Fig. 1j-l). These results indicate that PDZD8 is a novel substrate of AMPK that is activated by the lysosomal glucose sensing pathway. Re-introduction of PDZD8-WT, but not PDZD8-T527A, into *PDZD8*^−/−^ MEFs rescued the promotion of ER-mitochondria contact by glucose starvation (Fig. 1m-p and Extended Data Fig. 3c-g). Therefore, PDZD8 plays an important role in promoting ER-mitochondria contact under glucose starvation, in an AMPK-dependent manner.

**Fig. 1.**
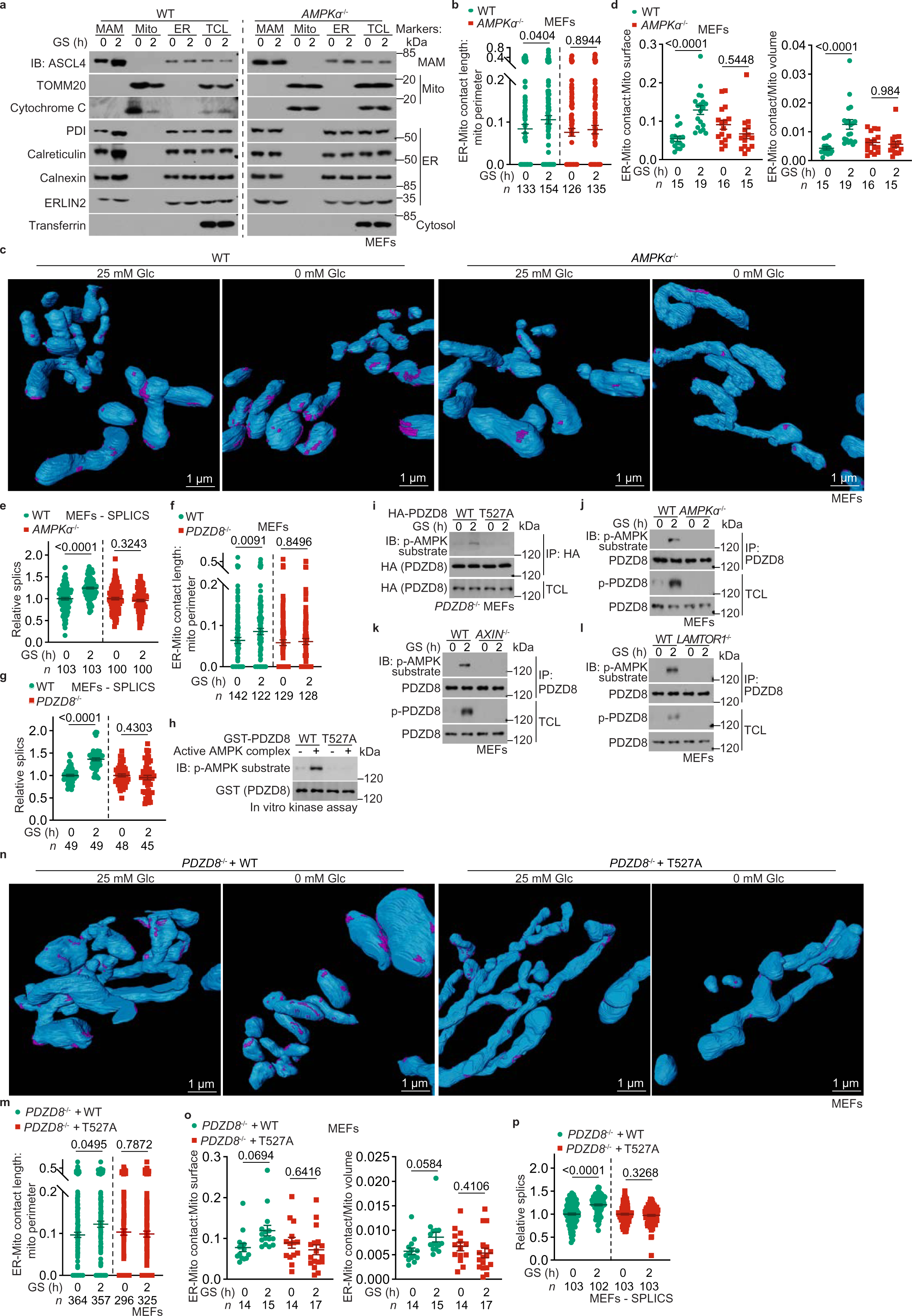
PDZD8 is a substrate of AMPK. **a-e**, AMPK promotes the association between mitochondria and ER. Wildtype MEFs and *AMPK*α^−/−^ MEFs were glucose-starved (GS) for 2 h, and were subjected to purification of MAM, mitochondria (mito), and ER (**a**), to TEM (**b**) or FIB-SEM (**c**, **d**), and to quantification of the signal of SPLICS (**e**, in cells stably expressing SPLICS reporter). The formation of ER-mitochondria contact was determined either by the protein levels of markers for each subcellular structure (**a**, via immunoblotting), by the length of ER-mitochondria contact normalised to mitochondrial perimeter on the TEM images (**b**, see also other readouts in Extended Data Fig. 1a-d), by area/volume of contact normalised to the area/volume of mitochondria on the FIB-SEM images (**c**, where mitochondria are decorated in blue, and the ER-mitochondria contact magenta, and the same hereafter for all FIB-SEM images; see also other readouts in Extended Data Fig. 1e), or by the numbers of SPLICS puncta (**e**, normalised to the unstarved group of each genotype, and the same hereafter for all SPLICS assays, unless otherwise specified). **f**, **g**, PDZD8 is required for the promotion of ER-mitochondria contact. Experiments were performed as in **b** and **e**, except that wildtype and *PDZD8*^−/−^ MEFs were used. **h**, AMPK phosphorylates T527 residue of PDZD8 in vitro. Some 1 μg of GST-tagged recombinant PDZD8 or its T527A mutant was incubated with 0.1 μg of holo-AMPK pre-phosphorylated by CaMKK2, followed by determining the phosphorylation of PDZD8 by immunoblotting. **i**-**l**, AMPK phosphorylates T527 residue of PDZD8 in cells. MEFs with HA-tagged PDZD8 or PDZD8-T527A stably expressed (**i**), or with knockout of *AMPK*α (**j**), *AXIN* (**k**), or *LAMTOR1* (**l**), were glucose-starved for 2 h, followed by immunoprecipitation of HA-PDZD8 (**i**) or of endogenous PDZD8 (**j**-**l**). The immunoprecipitates were then subjected to immunoblotting to determine the levels of p-T527. **m**-**p**, Mutation of T527 to alanine abolishes the ability of PDZD8 to promote ER-mitochondria contact. Experiments were performed as in **b** to **e**, except that *PDZD8*^−/−^ MEFs with wildtype PDZD8 or PDZD8-T527A re-introduced, were used. Data are shown as mean ± s.e.m., with *n* values (labelled on each panel) representing the numbers of mitochondria (**b**, **d**, **f**, **m**, **o**) or cells (**e**, **g**, **p**); *P* values were determined by two-tailed Mann-Whitney test (**b**, **f**, **m**, and WT cells of **g**), or by two-way ANOVA, followed by Tukey (**d**, **o**) and by unpaired two-tailed Student’s *t*-test (**e**, **p** and KO cells of **g**). Experiments in this figure were performed three times, except **i** and **j** four times.

## PDZD8 mediates the utilisation of glutamine during early starvation

We next determined whether PDZD8 participates in the dynamic utilisation of alternative carbon sources, i.e., glutamine and fatty acid, after glucose starvation. We pre-treated MEFs separately with both [U-^13^C]palmitate and [U-^13^C]glutamine, and subjected these cells to glucose starvation. The rates of glutamine utilisation, as determined by the levels of ^13^C-labelling of TCA cycle intermediary metabolites (determined by the levels of m+5 α-ketoglutarate (α-KG); and m+4 succinate, fumarate, malate and citrate) in MEFs pre-treated with [U-^13^C]glutamine, were elevated within 2 h of glucose starvation (Fig. 2a and Extended Data Fig. 4a, b). In comparison, increase of ^13^C-labelled TCA cycle intermediary metabolites in [U-^13^C]palmitate pre-treated MEFs (determined by the levels of m+2 α-KG, succinate, fumarate, malate and citrate) occurred at around 12 h of starvation, much slower than that with [U-^13^C]glutamine (Fig. 2b and Extended Data Fig. 4c-e; see also Supplementary Note 3 for detailed analysis). Therefore, the promotion of glutaminolysis under glucose starvation occurs ahead of the increase of FAO. We also found that the increased utilisation of glutamine, as early as 1 h after glucose starvation, was able to compensate for the reduction of oxidation of glucose in the TCA cycle (Fig. 2c and Supplementary Note 4), indicative of a shift of carbon source utilisation from glucose to glutamine from the early stage of glucose starvation. Knockout of *AMPK*α, *AXIN* or *LAMTOR1* all blocked the promotion of both glutaminolysis within 2-h glucose starvation, and FAO at 8-h glucose starvation in MEFs (Fig. 2d-g, Extended Data Fig. 4f-k), leading to deficient energy levels with a drastic accumulation of AMP (Extended Data Fig. 4l, see also ref. ^40,64,65^). Importantly, the AMPK-PDZD8 axis is specifically involved in the promotion of glutaminolysis, as re-introduction of PDZD8-T527A into *PDZD8*^−/−^ MEFs only blocked the promotion of glutaminolysis during early starvation, but not the increase of FAO that occurs later on (Fig. 2h, i, Extended Data Fig. 5a, b). We also determined the AMPK-PDZD8 axis-dependent promotion of glutaminolysis at the organismal level. Similar to those observed in MEFs, muscular and hepatic glutaminolysis was also found to be promoted much earlier than FAO in starved mice, as measured by using infused U-^13^C-labelled glutamine and palmitate (Fig. 2j, k, Extended Data Fig. 5d-g; see AMPK activation in tissues in Extended Data Fig. 5c; see also levels of serum β-hydroxybutyrate, an indicator of hepatic FAO^66,67^, as an additional control in Fig. 2n). Muscle-specific re-introduction of PDZD8-T527A, but not PDZD8-WT, into *PDZD8*-MKO (muscle-specific knockout) mice blocked the fasting-induced glutaminolysis (Fig. 2l, m, Extended Data Fig. 5h-k; see validation data of muscular PDZD8 re-introduction in Extended Data Fig. 5l), showing a similar level of glutaminolysis to that after muscle-specific knockout of *AMPK*α (Fig. 2j, k, Extended Data Fig. 5d-g; see validation data of *AMPK*α-MKO mice in Extended Data Fig. 5c). In line with the results from the isotopic labelling experiments, we observed a rapid increase of oxygen consumption rates (OCR) in both 2 h glucose-starved MEFs and 8 h-starved muscle tissues, which did not occur in PDZD8-T527A-reintroduced *PDZD8*^−/−^ MEFs and *AMPK*α^−/−^ MEFs (Fig. 2o), or PDZD8-T527A-reintroduced *PDZD8*-MKO, and *AMPK*α-MKO mouse muscles (Fig. 2p). In addition, knockdown of *GLS1* (both *GAC* and *KGA* isoforms) or treatment of GLS1 inhibitor BPTES^68^ blocked the increase of OCR (Fig. 2q, Extended Data Fig. 5m), while knockout of *CPT1* (both *CPT1*α and *CPT1*β) or treatment of CPT1 inhibitor etomoxir^69^ failed to do so (Fig. 2r, Extended Data Fig. 5m). As an additional control, the protein contents of the mitochondrial electron transport chain or the efficiency of electron transfer was unchanged after glucose starvation (Extended Data Fig. 5n), re-assuring that it is the utilisation of glutamine that elevates OCR. Together, these results demonstrate that AMPK phosphorylates PDZD8 at T527 to promote glutamine utilisation ahead of use of fatty acids, to compensate for depletion of glucose under starvation.

**Fig. 2.**
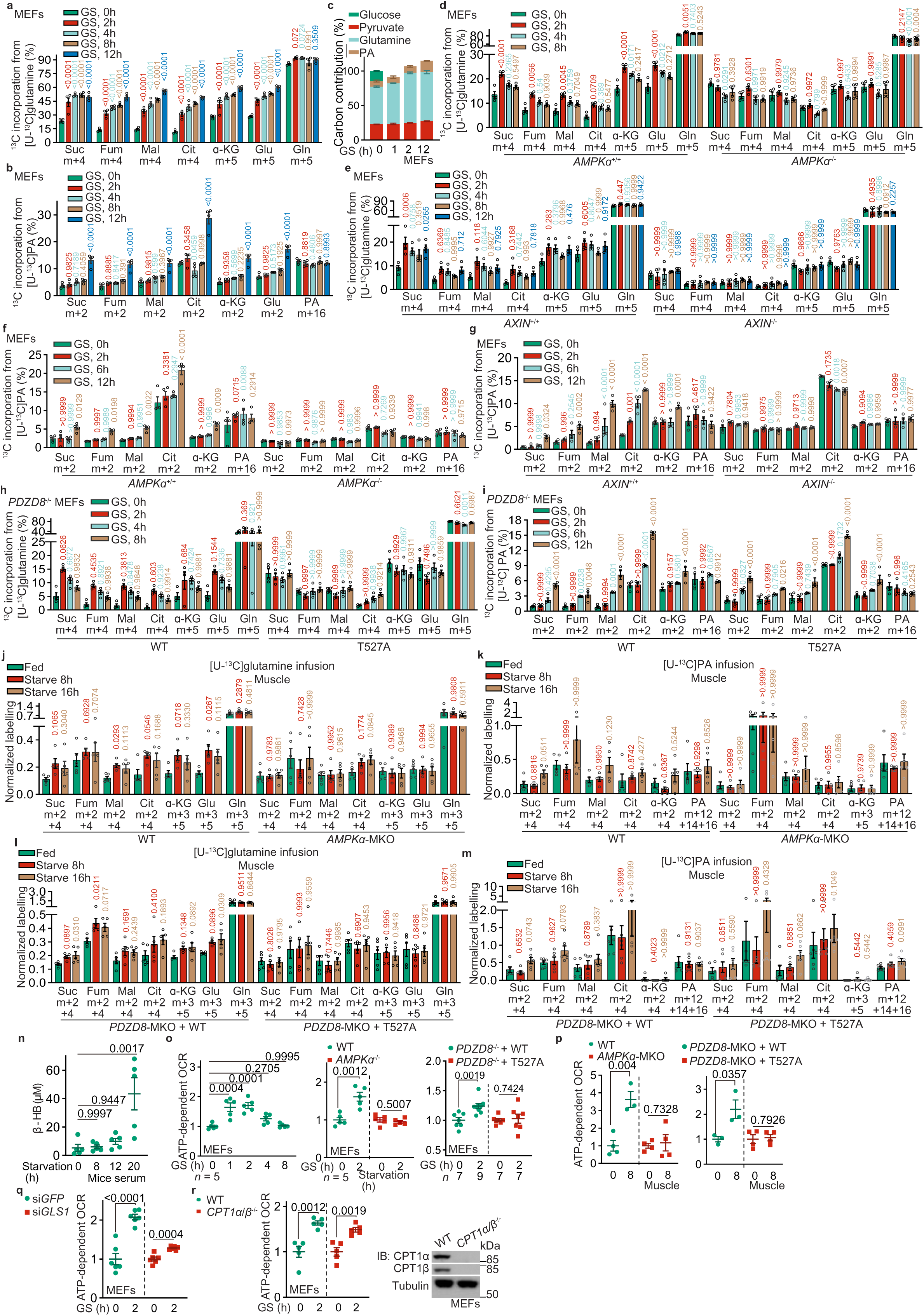
PDZD8 promotes the utilisation of glutamine during early starvation. **a, b**, Glutaminolysis is promoted ahead of the increase of FAO under glucose starvation. MEFs were glucose starved for desired durations. At 20 min and 12 h before sample collection, cells were labelled with [U-^13^C]-glutamine (**a**) and [U-^13^C]-PA (**b**), respectively, followed by determination of the levels of labelled TCA cycle intermediates, including succinate (Suc), fumarate (Fum), malate (Mal), citrate (Cit), α-ketoglutarate (α-KG), along with glutamate (Glu), by GC-MS. Levels of m+5 α-ketoglutarate and glutamate; and m+4 succinate, fumarate, malate and citrate that reflect the rates of glutaminolysis (**a**), along with levels of m+2 α-ketoglutarate, glutamate, succinate, fumarate, malate and citrate that reflect the rates of FAO (**b**), were shown. See also Extended Data Fig. 4b and c for the levels of other isotopomers of the labelled metabolites shown in **a** and **b**. Data are shown as mean ± s.e.m.; *n* = 4 samples for each condition; *P* values were determined by two-way ANOVA, followed by Tukey, all compared to the unstarved group. **c**, Glutamine utilisation compensates for the reduction of glucose oxidation in the TCA cycle in low glucose. MEFs were respectively labelled with [U-^13^C]-glutamine, [U-^13^C]-PA and [U-^13^C]-pyruvate, all for 24 h, followed by glucose starvation for 1 h, 2 h and 12 h. For the group of normally cultured (unstarved) cells, MEFs were labelled with [U-^13^C]-glucose for 24 h, in addition to labelling with [U-^13^C]-glutamine, [U-^13^C]-PA and [U-^13^C]-pyruvate as in the glucose-starved conditions. The contributions of each carbon source to the TCA cycle, as calculated by the total levels of labelled succinate (sum of m+1 to m+4), were then shown in a stacked bar chart. Data are shown as mean ± s.d.; *n* = 4 samples for each condition. **d**-**i**, AMPK-PDZD8 axis promotes the utilisation of glutamine early during starvation in MEFs. Experiments in **d**, **e**, and **h** (for determining glutaminolysis) were performed as in **a**, and those in **f**, **g**, and **i** (for determining FAO) as in **b**, except that *AMPK*α^−/−^MEFs (**d**, **f**), *AXIN*^−/−^ MEFs (**e**, **g**), and *PDZD8*^−/−^ MEFs with wildtype PDZD8 or PDZD8-T527A re-introduced (**h**, **i**) were used. Data are shown as mean ± s.e.m.; *n* = 4 samples for each condition; and *P* values were determined by two-way ANOVA, followed by Tukey, all compared to the unstarved group. **j**-**m**, AMPK-PDZD8 axis promotes the utilisation of glutamine early during starvation in mouse muscle. Mice were starved for desired durations, followed by jugular-vein infused with [U-^13^C]-glutamine or [U-^13^C]-PA tracer, for 2 h, respectively. Mice were then sacrificed, followed by determining the rates of glutaminolysis and FAO as in **a** and **b**. After normalisation to the serum levels of corresponding labelled tracers, data were shown as mean ± s.e.m.; *n* = 5 (**j**, **k**, **l**), or 6 (**m**) samples for each condition; and *P* values were determined by one-way ANOVA, followed by Tukey (**j**; **l**; α-KG and citrate of **k**; succinate, malate and PA from WT mice of **k**; succinate, malate and PA of **m**; and fumarate from WT mice in **m**), or Dunn (others), all compared to the unstarved group. **n**, Induction of serum β-hydroxybutyrate, an indicator of hepatic FAO, occurs after prolonged starvation. Mice were starved for desired durations, followed by determining the levels of serum β-hydroxybutyrate. Data are shown as mean ± s.e.m.; *n* = 5 mice for each condition; and *P* values were determined by one-way ANOVA, followed by Tukey. **o**, **p**, AMPK-PDZD8 axis promotes OCR early during starvation. Wildtype MEFs, PDZD8-T527A-reintroduced *PDZD8*^−/−^ MEFs and *AMPK*α^−/−^ MEFs (**o**), or wildtype, PDZD8-T527A-reintroduced *PDZD8*-MKO, and *AMPK*α-MKO mice (**p**) were glucose-starved for desired durations, followed by determining OCR through Seahorse Analyzer. Data were normalised to the unstarved group of each genotype (same hereafter for all OCR measurements), and are shown as mean ± s.e.m.; *n* = 5 (**o**), 3 (muscles from starved WT mice and the PDZD8-WT-reintroduced PDZD8-MKO mice, of **p**), or 4 (**p**, others) biological replicates for each condition; and *P* values were determined by one-way ANOVA, followed by Tukey (left panel of **o**) or by unpaired two-tailed Student’s *t*-test (others). **q**, **r**, Inhibition of glutaminolysis, but not FAO, prevents OCR increases. MEFs with *GLS1* knockdown (**q**), or *CPT1* knockout (**r**) were glucose-starved for 2 h (early starvation), followed by determining OCR as in **o**. Data are shown as mean ± s.e.m.; *n* = 6 (**q**) or 5 (**r**) biological replicates for each condition; and *P* values were determined by unpaired two-tailed Student’s *t*-test. See also knockout validation data of *CPT1* on the right panel of **r**. Experiments in this figure were performed three times, except **o** and **p** four times.

## PDZD8 promotes GLS1 activity

We next explored the mechanism through which PDZD8 promotes glutaminolysis. It was found that the activity of GLS1 was significantly promoted in cells starved for glucose, by using a semi-permeabilised assay system (Fig. 3a, see detailed protocol in Methods section). Knockout of *AMPK*α blocked the promotion of GLS1 activity (Fig. 3a). We also found that re-introduction of PDZD8-WT, but not PDZD8-T527A, rescued glucose starvation-induced GLS1 activity in *PDZD8*^−/−^ MEFs (Fig. 3b). These data indicate that the AMPK-PDZD8 axis controls glutaminolysis through regulating GLS1. As a control, we also examined if glucose starvation causes GLS1 filamentation (supratetrameric oligomerisation) that has been shown to enhance the catalytic activity of GLS1 under glutamine starvation^70^, and found that GLS1 oligomerisation was not changed, indicating that GLS1 filamentation did not apply to the regulation by glucose starvation (Extended Data Fig. 6a). We also performed cell-free assays, and found that the wildtype PDZD8, but not the AMPK-unphosphorylable T527A mutant, promoted GLS1 activity in an AMPK-dependent manner (Fig. 3c, d; see *K*_m_ and *k*_cat_ values of each reaction in Supplementary Table 2). Free inorganic phosphate in cell-free systems could further activate GLS1 on top of the activation by PDZD8 (Fig. 3e, f and Supplementary Table 2), in line with inorganic phosphate being a co-factor of GLS1^71,72^, indicating that the phosphorylation of PDZD8 and the inorganic phosphate stimulate GLS1 via two independent mechanisms. Data in Fig. 3c-f also revealed that AMPK-phosphorylated PDZD8 increased the affinity of GLS1 towards the substrate glutamine (after phosphorylation by AMPK: *K*_m_ of KGA decreased from 14.63 mM to 6.08 mM in the absence of inorganic phosphate, and from 7.30 mM to 3.46 mM in the presence of inorganic phosphate; and ditto for GAC). Note that the whole-cell concentrations of glutamine were around 2 mM (see also ref. ^73–75^), and remained similar after glucose starvation (Fig. 3g). These data indicate that GLS1 is unsaturated with its substrate at all times, consistent with results that PDZD8 can boost glutamine catabolism to increase glutaminolysis.

**Fig. 3.**
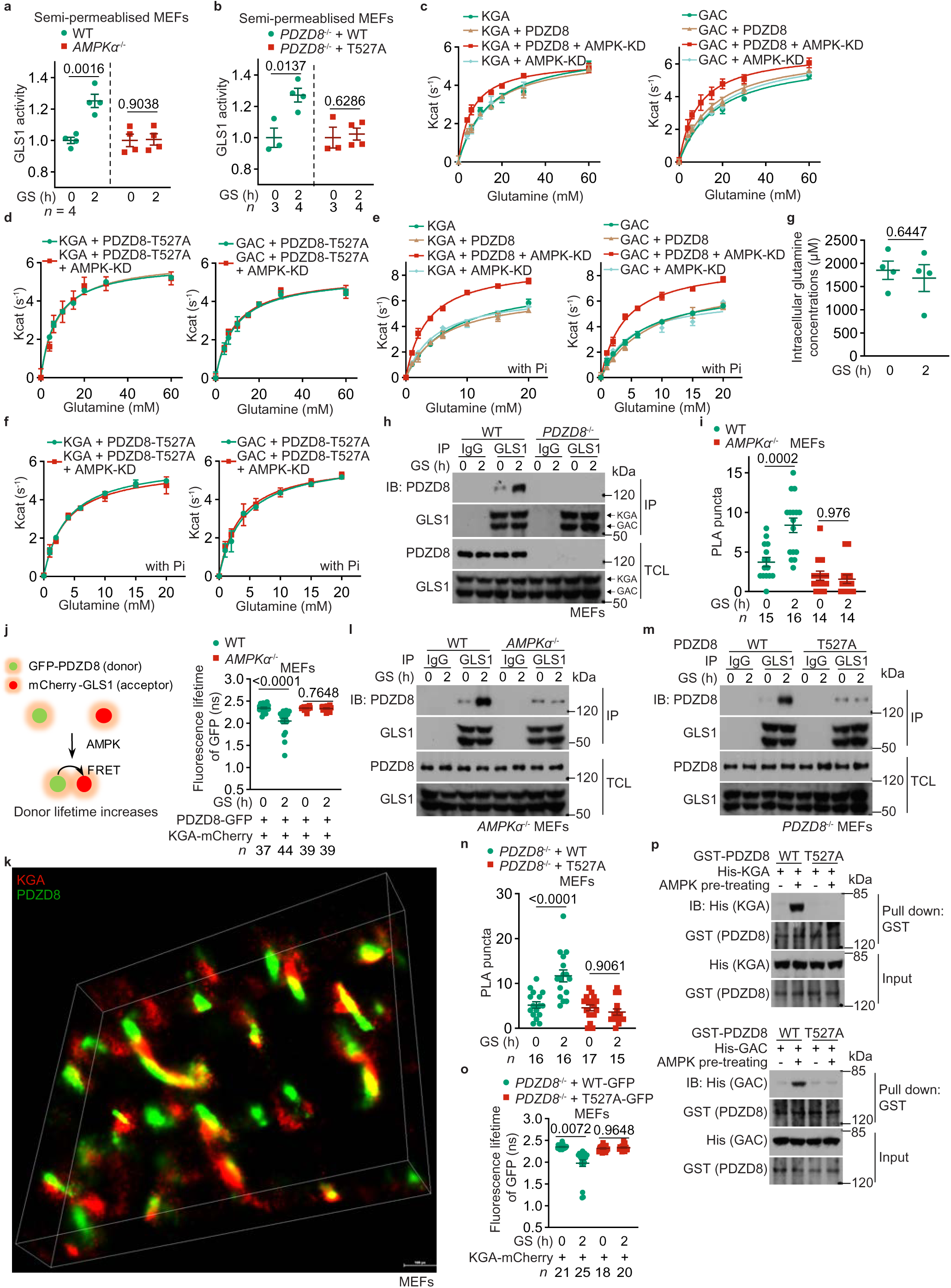
PDZD8 promotes GLS1 activity. **a, b**, AMPK-PDZD8 axis promotes GLS1 activity in permeabilised cells. Wildtype MEFs, *AMPK*α^−/−^ MEFs (**a**), and wildtype PDZD8 or PDZD8-T527A-reintroduced *PDZD8*^−/−^ MEFs (**b**) were glucose-starved for 2 h, followed by permeabilisation with 0.01% (v/v) NP-40. The activities of GLS1, as evaluated by the production of glutamate after glutamine addition, were then measured. Data are shown as mean ± s.d.; *n* = 4 (**a**), or labelled on the panel (**b**; representing biological replicates) for each condition; and *P* values were determined by Mann-Whitney test (T527A cells of **b**) and by unpaired two-tailed Student’s *t*-test (others). **c**-**f**, AMPK-PDZD8 axis promotes GLS1 activity in cell-free systems. Recombinant KGA (left panel) and GAC (right panel) isozymes of GLS1 were mixed with recombinant PDZD8 (**c**, **e**) or PDZD8-T527A (**d**, **f**) protein that was pre-incubated with the constitutive active kinase domain of AMPKα (AMPK-KD; see “Phosphorylation of PDZD8 by AMPK in vitro” in Methods section), followed by determination of the enzymatic activities of GLS1. In **e** and **f**, 20 mM K_2_HPO_4_ (Pi) was added to the reactions. Data are shown as mean ± s.d.; *n* = 3 biological replicates for each condition. See also *K*_m_ and *k*_cat_ values for each reaction in Supplementary Table 2. The experiments in **c** and Fig. 4a were performed at same time and shared control (the KGA- and GAC-alone groups), and ditto for **e** and Fig. 4b. **g**, Glucose starvation does not change the intracellular levels of glutamine. Cells were glucose-starved for 2 h, and the intracellular levels of glutamine were determined via HPLC-MS. Data are shown as mean ± s.e.m.; *n* = 4 samples for each condition; and *P* values were determined by unpaired two-tailed Student’s *t*-test. **h**, **l**, **m**, PDZD8 interacts with GLS1, depending on AMPK. Wildtype MEFs and *PDZD8*^−/−^ MEFs (**h**), *AMPK*α^−/−^ MEFs (**l**), and wildtype PDZD8 or PDZD8-T527A-reintroduced *PDZD8*^−/−^ MEFs (**m**), were glucose-starved for 2 h. Endogenous GLS1 proteins (both KGA and GAC) were immunoprecipitated, followed by immunoblotting to determine co-precipitated PDZD8. **i**, **j**, **n**, **o**, AMPK promotes PDZD8-GLS1 interaction in situ. *AMPK*α^−/−^ MEFs (**i**, **j**), or *PDZD8*^−/−^ MEFs (**n**, **o**) were infected with lentiviruses carrying HA-tagged PDZD8 or PDZD8-T527A (**i**, **n**; for PLA assay), or KGA-mCherry, along with PDZD8-GFP (**j**, **o**; for FRET-FLIM assay, see strategy of this assay on the left panel of **j**) or PDZD8-T527A-GFP (**o**). Cells were then glucose-starved for 2 h, followed by quantifying the numbers of PLA puncta in each cell (**i**, **n**; data are shown as mean ± s.e.m.; *n* values (labelled on each panel) represent cell numbers for each condition), or measuring the fluorescence lifetime of GFP (the FRET donor; **j**, **o**; data are shown as mean ± s.e.m.; *n* values represent cell numbers for each condition); and *P* values were determined by two-way ANOVA, followed by Tukey. **k**, STORM images showing that PDZD8 is juxtaposed with GLS1 inside cells. MEFs stably expressing FLAG-tagged KGA and Myc-tagged PDZD8 were subjected to STORM imaging, and the representative, reconstituted 3D-STORM image is shown. **p**, AMPK promotes PDZD8-GLS1 interaction in vitro. Recombinant His-tagged KGA (upper panel) and GAC (lower panel) isozymes of GLS1 were separately mixed with recombinant GST-tagged PDZD8 or PDZD8-T527A protein that was pre-incubated with AMPK pre-phosphorylated with CaMKK2 (see “Phosphorylation of PDZD8 by AMPK in vitro” in Methods section), followed by pulling down GST-tag and immunoblotting. Experiments in this figure were performed three times.

We also tested for possible interaction between PDZD8 and GLS1, and found that they indeed interacted with each other, endogenous or ectopic, and that the interaction became more prominent in cells starved for glucose, as determined by co-immunoprecipitation (Fig. 3h, Extended Data Fig. 6b, c). This glucose starvation-enhanced PDZD8-GLS1 interaction could also be detected in situ, by both the proximity ligation assays (PLA) in fixed MEFs, and the FRET-FLIM assay in living MEFs (Fig. 3i, j). We also observed that GLS1 is juxtaposed with PDZD8 as determined by both structured illumination microscopy (SIM; Extended Data Fig. 6d) and stochastic optical reconstruction microscopy (STORM; Fig. 3k). We also showed that PDZD8 is juxtaposed with the mitochondrial marker TOMM20, and GLS1 with the ER marker PDI (Extended Data Fig. 6e, f). Given that PDZD8 resides on ER^60^, and GLS1 mitochondria^76^, these data indicate that PDZD8-GLS1 interaction occurs at the ER-mitochondria contact. Knockout of *AMPK*α, or reintroduction of PDZD8-T527A abrogated the increase of the interaction between PDZD8 and GLS1 in low glucose (Fig. 3l-o). Consistently, *in vitro* reconstitution experiments showed that prior phosphorylation with recombinant AMPK increased the affinity of PDZD8, but not PDZD8-T527A towards bacterially purified GLS1 (Fig. 3p). Domain mapping experiments showed that the C-terminus of PDZD8 (PDZD8-CT) constitutes the interface for GLS1 (Extended Data Fig. 7a), as PDZD8-CT alone was sufficient to promote GLS1 activity to the same extent as the full-length PDZD8 pre-treated with AMPK (Fig. 4a, b). Consistently, re-introduction of PDZD8-CT into *PDZD8*^−/−^ MEFs promoted the utilisation of glutamine and OCR even in high glucose, to similar levels by full-length PDZD8 in low glucose (Fig. 4c, d, Extended Data Fig. 7b). These data all suggest that the CT domain acts in a dominant-positive manner for interacting with GLS1. Consistently, we found the N-terminus of PDZD8 (PDZD8-NT) interacts with PDZD8-CT, when expressed separately as truncate proteins in high glucose, and the interaction was abolished in low glucose when AMPK is activated, which indicates that AMPK phosphorylation releases the intramolecular autoinhibition of the N-terminal region towards the C-terminal region of PDZD8 (Fig. 4j, Extended Data Fig. 8c). Indeed, AMPK phosphorylation led to an increased affinity of full-length PDZD8 towards GLS1 to an extent similar to that of PDZD8-CT alone towards GLS1 (Fig. 4i; see Supplementary Note 5 for details). In addition, we generated a GLS1 mutant (GLS1-33A) carrying mutations to alanine of the 33 amino acid residues on the interface for interacting with PDZD8, which were identified by in silico docking assays (Extended Data Fig. 8a). Although GLS1-33A showed similar enzymatic activities to that of the wildtype GLS1, it was no longer regulated by PDZD8 (Fig. 4e, f). GLS1-33A also blocked the promotion of glutaminolysis or OCR in low glucose (Fig. 4g, h, Extended Data Fig. 8b). Results above demonstrate that PDZD8 promotes GLS1 activity through direct interaction in low glucose. We also found that PDZD8-GLS1 interaction is responsible for tightening the ER-mitochondria contact (Fig. 4k, l, Extended Data Fig. 8d-g; see Supplementary Note 5 for details).

**Fig. 4.**
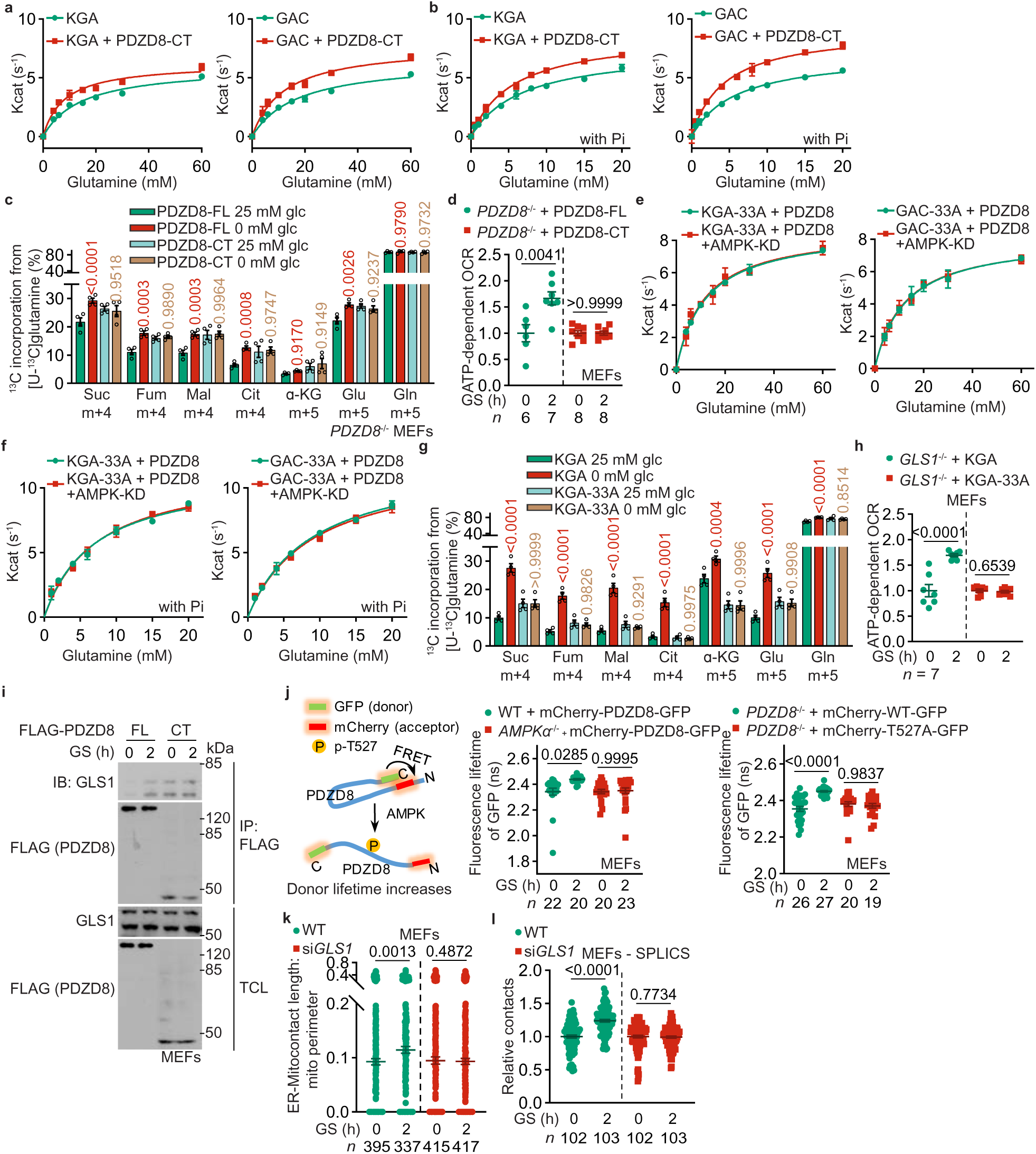
Interaction of PDZD8 promotes GLS1 activity. **a, b**, PDZD8-CT that constitutively interacts with GLS1, promotes GLS1 activity in vitro independently of AMPK. Recombinant KGA (left panel) or GAC (right panel) isozymes of GLS1 was mixed with recombinant PDZD8-CT, followed by determining the enzymatic activities of GLS1 in the presence (**b**) or absence (**a**) of 20 mM K_2_HPO_4_ (Pi). Data are shown as mean ± s.d.; *n* = 3 for each condition. See also *K*_m_ and *k*_cat_ values for each reaction in Supplementary Table 2. The experiments in **a** and Fig. 3c were performed at same time and shared control (the KGA- and GAC-alone groups), and ditto for **b** and Fig. 3e. **c**, **d**, PDZD8-CT promotes glutaminolysis and OCR in high glucose. *PDZD8*^−/−^ MEFs were infected with lentiviruses carrying full-length (FL) PDZD8 PDZD8-CT, followed by incubating in medium containing doxycycline for 12 h. Cells were then labelled with [U-^13^C]-glutamine to determine glutaminolysis (**c**, performed as in Fig. 2a), or subjected to Seahorse Analyzer to determine OCR (**d**). Data are shown as mean ± s.e.m.; *n* = 4 (**c**), or labelled on the panel (**d**; representing biological replicates) for each condition; and *P* values were determined by two-way ANOVA, followed by Tukey (*P* values in **c** represent the comparisons between the starved and the unstarved groups of each genotype). **e**-**h**, GLS1-33A that loses the interface for PDZD8, fails to promote GLS1 activity (**e**, **f**), glutaminolysis (**g**) or OCR (**h**) in low glucose. Experiments in **e** and **f** were performed as in **a** and **b**, except that the recombinant KGA-33A (left panel) and GAC-33A (right panel) were mixed with AMPK-phosphorylated PDZD8. See also lowered *K*_m_ and increased *k*_cat_ values in each reaction in Supplementary Table 2. Experiments in **g** and **h** were performed as in **c** and **d**, except that *GLS1*^−/−^ MEFs with wildtype KGA or KGA-33A stably expressed were used. Data are mean ± s.d.; *n* = 3 (**e**, **f**) or 4 (**g**), or labelled on the panel (**h**; representing biological replicates) for each condition; and *P* values were determined by two-way ANOVA, followed by Tukey (**g**) or by unpaired two-tailed Student’s *t*-test (**h**). **i**, AMPK releases the autoinhibition of PDZD8-NT towards PDZD8-CT. MEFs stably expressing FLAG-tagged PDZD8-FL or PDZD8-CT were glucose-starved for 2 h, followed by immunoprecipitation with anti-FLAG and immunoblotting for co-precipitated GLS1. **j**, AMPK causes PDZD8-NT to move away from PDZD8-CT. *AMPK*α^−/−^ MEFs (middle panel), or *PDZD8*^−/−^ MEFs (right panel) were infected with lentiviruses carrying mCherry-PDZD8-GFP (middle and right panels) or mCherry-PDZD8-T527A-GFP (right panel), followed by determination of the fluorescence lifetime of GFP (FRET donor; see principles of this assay on the left panel). Data are shown as mean ± s.e.m.; *n* values were labelled on the panel representing cell numbers; and *P* values were determined by two-way ANOVA, followed by Tukey. **k**, **l**, PDZD8-GLS1 interaction is responsible for tightening the ER-mitochondria contact in low glucose. MEFs with knockdown of *GLS1* were glucose-starved for 2 h, followed by determination of the formation of ER-mitochondria via TEM (**k**) or SPLICS staining (**l**). Data were analysed as in Fig. 1b and 1e, and are shown as mean ± s.e.m.; *n* values represent mitochondria (**k**) or cell (**l**) numbers for each condition; and *P* values were determined by Mann-Whitney test (**k**), or by unpaired two-tailed Student’s *t*-test (**l**). Experiments in this figure were performed three times.

## Phenotypes in animal models

We next explored the physiological functions of the AMPK-PDZD8-GLS1 axis in animal models. We constructed a *pdzd*-*8* (PDZD8 homologue in *C*. *elegans*) knockout *C*. *elegans* strain with re-introduced human wildtype PDZD8 or T527A mutant (validated in Extended Data Fig. 9a). We next starved the nematodes by treating with the nonmetabolisable glucose analogue 2-deoxy-glucose (2-DG), which has been shown to activate AMPK in nematodes due to decreased levels of FBP^42,77^. We found that expression of PDZD8-T527A, but not wildtype PDZD8, abrogated the effect of 2-DG in the promotion of ER-mitochondria contact (Fig. 5a, Extended Data Fig. 9b). We also observed a T527 phosphorylation-dependent increase of glutaminolysis in nematodes in isotopic labelling experiments (Fig. 5b, Extended Data Fig. 9c). In addition, OCR in the 2-DG-treated *C*. *elegans* was also found to be increased in a PDZD8-T527 phosphorylation-dependent manner (Fig. 5c). We next determined whether AMPK-dependent PDZD8 phosphorylation was also critical for the lifespan extension of *C*. *elegans* under the 2-DG treatment, and found that expression of PDZD8-T527A blocked the extension of lifespan (Fig. 5d; see also statistical analyses on Supplementary Table 3, and the same hereafter for all lifespan data). Similarly, the lifespan-extending effects of constitutively active aak-2 (AMPKα homologue in *C*. *elegans*)^78^ were abrogated in nematodes expressing PDZD8-T527A (Fig. 5e), suggesting that PDZD8 and AMPK lie in the same pathway. We also found that the enhanced glutaminolysis is required for AMPK-PDZD8-mediated lifespan extension, as depletion of all the three *glna* genes (*glna*-*1* to *glna*-*3*; glutaminase homologues in *C*. *elegans*) by knockdown of *glna*-*2* in *glna*-*1* and *glna*-*3* double knockout strain, or re-introduction of GLS1-33A defective in interacting with PDZD8 into this glna-null strain, blocked lifespan extension in low glucose (Fig. 5f, g). The linkage between p-T527-dependent increase of OCR and lifespan extension is reminiscent of the AMPK-mediated mitohormesis, which is defined as an increase in fitness and longevity consequential to the adaptive responses to mild mitochondrial oxidative stress (mitochondrial ROS) induced under the conditions such as low glucose^77,79^. Consistent with the characteristics of mitohormesis, we observed a PDZD8-T527 phosphorylation- and GLS1-dependent increase of mitochondrial ROS in *C*. *elegans* under 2-DG treatment, as assessed by the fluorescent signal of mitoSOX dye which specifically responds to the mitochondrial ROS (Fig. 5h, Extended Data Fig. 9d). The ROS increase levelled off soon afterwards, accompanied with an increased expression of ROS-depleting enzymes (such as SOD), as determined via RNA-sequencing (Fig. 5i). We also cultured nematodes on agar containing diluted bacteria to mimic caloric restriction^80,81^, and similarly found that reintroduction of PDZD8-T527A to *pdzd*-*8* knockout nematodes, depletion of *glna*-*1* to *glna*-*3* in nematodes, or expression of GLS1-33A in glna-depleted nematodes abolished the effects of mitohormesis and lifespan-extension after calorie restriction (Fig. 5j-m, Extended Data Fig. 9e). Consistently, the expression of PDZD8-T527A or GLS1-33A blocked the enhancement of pharyngeal pumping rates and the resistance to oxidative stress in nematodes subjected to caloric restriction (Fig. 5n-r, Extended Data Fig. 9f-i). Data shown above indicate that the glucose starvation-promoted extension of lifespan and healthspan in nematodes depends on the AMPK-PDZD8 axis. We also examined the rejuvenating roles of the AMPK-PDZD8 axis in mice. As shown in Fig. 5s-u, after three months of caloric restriction (by reducing the daily food supply to 70% which sufficiently induced p-T527 of PDZD8, see Extended Data Fig. 10a), aged (8-month-old) *PDZD8*-MKO mice with muscle-specific re-introduction of wildtype PDZD8 showed a significant increase of running distance, duration, grip strength and muscular NAD^+^ levels. Caloric restriction also led to a transient increase of ROS, or mitohormesis in the muscle in these mice (Extended Data Fig. 10b). Such rescued phenotypes were not observed when PDZD8-T527A was re-introduced (Fig. 5s-u).

**Fig. 5.**
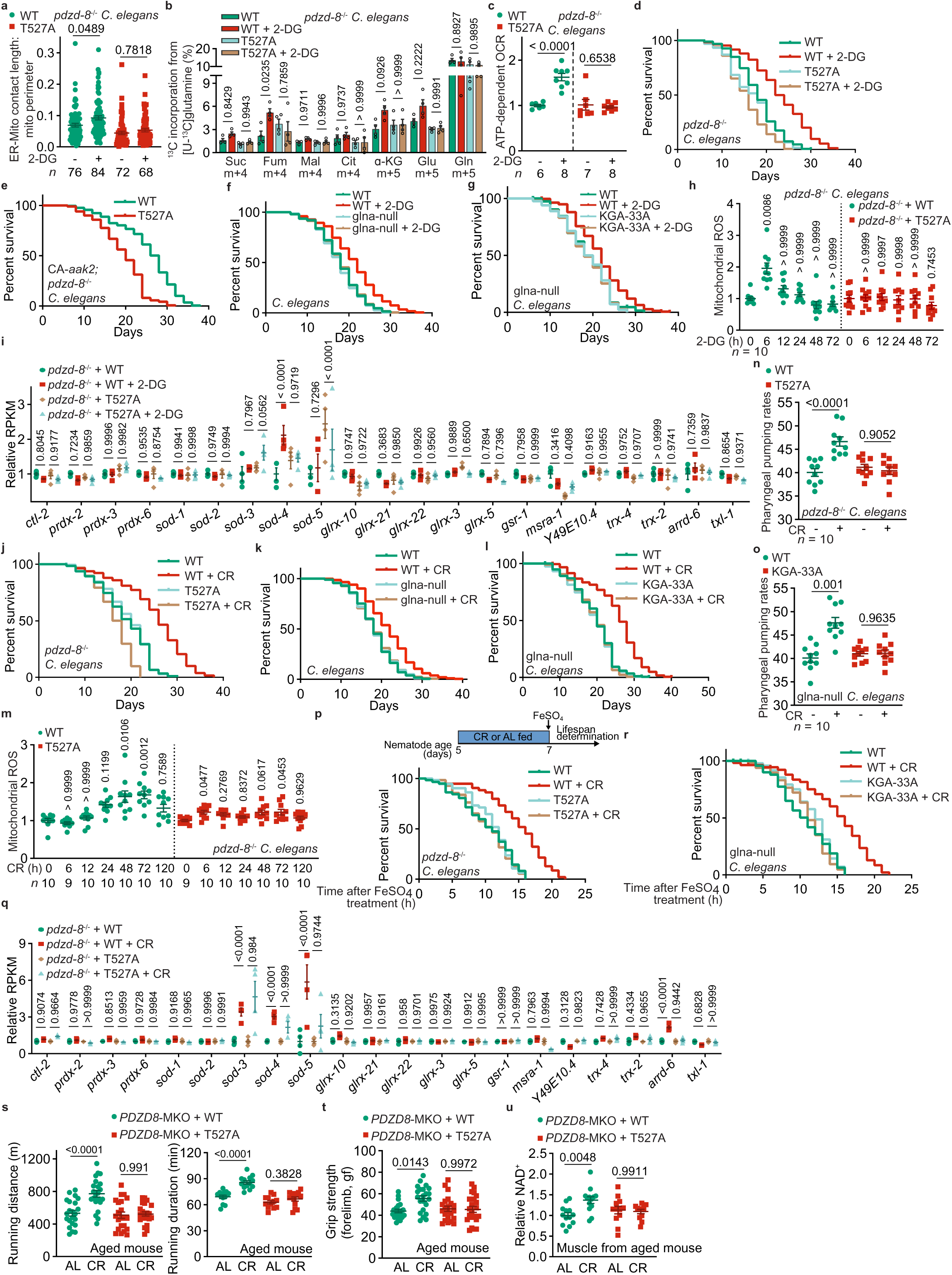
PDZD8 mediates rejuvenating effects of glucose starvation and caloric restriction. **a-c**, AMPK-PDZD8 axis promotes ER-mitochondria contact, glutaminolysis and OCR in nematodes under glucose starvation. The *pdzd*-*8^−/−^ C*. *elegans* strains with re-introduced human wildtype PDZD8 or T527A mutant were treated with 2-DG that mimics glucose starvation, for 2 days, followed by determination of the ER-mitochondria contact through TEM (**a**; data are shown as mean ± s.e.m.; *n* represents mitochondria numbers for each condition), glutaminolysis through determining the levels of labelled TCA cycle intermediates through GC-MS (**b**; see labelling procedures in “Determination of glutaminolysis and FAO rates” of Methods section; data are shown as mean ± s.e.m.; n = 4 samples for each condition), and OCR through Seahorse Analyzer (**c**; data are shown as mean ± s.e.m.; *n* values indicate biological replicates for each condition). *P* values were determined by two-way ANOVA, followed by Tukey (**a**, **b**), or by unpaired two-tailed Student’s *t*-test (**c**). **d**-**g**, AMPK-PDZD8 axis extends lifespan of nematodes under glucose starvation. The *pdzd*-*8^−/−^*nematodes with re-introduced PDZD8-T527A (**d**), with expression of constitutively active aak-2 (AMPKα homologue in *C*. *elegans*, CA-*aak2*; **e**); or wildtype (N2) nematodes with depletion of glna (GLS homologue in *C*. *elegans*, by knockdown of *glna*-*2* in *glna*-*1* and *glna*-*3* double knockout strain; **f**), and with reintroduction of KAG-33A (by re-introduction of KGA-33A into glna-knockout strain; **g**) were treated with 2-DG. Lifespan data are shown as Kaplan-Meier curves. See also statistical analyses on Supplementary Table 3, and the same hereafter for all lifespan data. **h**, **i**, AMPK-PDZD8 axis induces transient mitochondrial ROS and expression of ROS-depleting enzymes under glucose starvation. The *pdzd*-*8^−/−^ nematodes* with re-introduced PDZD8-T527A were treated with 2-DG for desired durations, followed by determination of mitochondrial ROS using the mitoSOX dye (**h**; data are shown as mean ± s.e.m.; *n* = 10 biological replicates for each condition). At 48 h after 2-DG treatment, RNA-sequencing was performed, and the mRNA levels of ROS-depleting enzymes were shown (**i**; data are shown as mean ± s.e.m.; *n* = 4 biological replicates for each condition). *P* values were determined by one-way ANOVA, followed by Dunn (WT nematodes of **h**), by Tukey (T527A nematodes of **h**), or two-way ANOVA, followed by Tukey (**i**). **j**-**l**, AMPK-PDZD8 axis mediates extension of lifespan in nematodes subjected to CR. Experiments in **j**, **k** and **l** were performed as in **d**, **f** and **g**, except nematodes were subjected to CR for 2 days. **m**, **q**, AMPK-PDZD8 axis induces transient mitochondrial ROS and expression of ROS-depleting enzymes in nematodes subjected to CR for desired duration (**m**, or 2 days in **q**). Experiments in **m** (data are shown as mean ± s.e.m.; with *n* values labelling on the panel) and **q** (data are shown as mean ± s.e.m.; *n* = 4 biological replicates for each condition) were performed as in **h** and **i**, except CR was applied. *P* values were determined by one-way ANOVA, followed by Dunn (WT nematodes of **h**), by Tukey (T527A nematodes of **h**), or two-way ANOVA, followed by Tukey (**q**). **n**, **o**, AMPK-PDZD8 axis promotes pharyngeal pumping rates in nematodes. The *pdzd*-*8*^−/−^ nematodes with re-introduced PDZD8-T527A (**n**), or glna-null nematodes with re-introduced KGA-33A (**o**), were subjected to CR for 2 days. Pharyngeal pumping rates were then determined, and are shown as mean ± s.d.; *n* = 10 biological replicates for each condition. *P* values were determined by two-way ANOVA, followed by Tukey. **p**, **r**, AMPK-PDZD8 axis promotes resistance to oxidative stress. The *pdzd*-*8*^−/−^ nematodes with re-introduced PDZD8-T527A (**p**), or glna-null nematodes with re-introduced KGA-33A (**r**), were subjected to CR for 2 days, followed by treating with 15 mM FeSO_4_ (see experimental timeline on the upper panel of **p**; AL fed; ad libitum fed). Lifespan data are shown as Kaplan-Meier curves. **s**-**u**, AMPK-PDZD8 axis plays a rejuvenating role in mice. Aged (8-month-old) *PDZD8*-MKO mice with muscle-specific re-introduction of wildtype PDZD8 or PDZD8-T527A were CR for 3 months, followed by determination of running distance (**s**; left panel), duration (**s**; right panel), grip strength (**t**) and muscular NAD^+^ levels (**u**). Data are shown as mean ± s.d.; *n* values (labelled on each panel) represents mouse numbers for each condition; *P* values were determined by two-way ANOVA, followed by Tukey. Experiments in this figure were performed three times.

## Discussion

We have pursued further the molecular and metabolic events that entail AMPK activation as a result of calorie restriction for extension of lifespan. The observation of the increase of mitochondria-ER contact has allowed us to identify a new substrate for AMPK, which is PDZD8 located on the ER. PDZD8 phosphorylated by AMPK releases its intramolecular inhibition, allowing its C-terminus to interact with and promotes the activity of GLS1 under physiological concentrations of glutamine (Extended Data Fig. 10c). Glutaminolysis is increased prior to a significant increase of the use of fatty acids, to sustain fuels and energy production, in line with glutamine being the most abundant circulating amino acid and rapidly replenished by other amino acids such as BCAA mobilised from labile proteins in muscle tissues during starvation^22^. Intriguingly, the increased glutaminolysis leads to a mild but rapid (within 1 h of glucose starvation in MEFs and 8 h starvation in mice) burst of mitochondrial ROS, which levels off quickly, conforming to the characteristics of mitohormesis^77,79,82^. The rapid decline of ROS after the burst is likely mediated by the induction of antioxidative genes, such as SOD, which may prepare the cells to prevent further ROS insults when cells have to mainly rely on fatty acids for energy in the case of prolonged starvation^77,79^. Consistently, we observed a T527 phosphorylation-dependent induction of antioxidative genes in nematodes after starvation or caloric restriction (Fig. 5i, q). The rapid increase of ROS along the adaptation to glutaminolysis may also, at least in part, explain the mechanism of Crabtree effect^83^, in that respiration rate is increased in low glucose.

Through isotope chasing experiments, we have shown that the increase of glutamine oxidation occurs prior to the increase of fatty acid oxidation. Glutamine offers several advantages over fatty acids. First of all, glutamine is an abundance amino acid, circulating at around 500 μM in the serum, and even higher in the interstitial space of muscle during fasting^10,84^, while the circulating and muscle interstitial free fatty acid is approximately 20-fold lower^85^. Perhaps also as a way to prevent cells from lipotoxity, levels of free fatty acid are strictly constrained inside cells or tissues, as two-thirds of fatty acid mobilised from adipose tissue after starvation is re-esterified into triglyceride (futile cycle), while the remaining one-third is burned by muscles^12,86^. Second, the rates of glutamine oxidation, at least in the muscle, are much faster than that of fatty acids. As in a simple experiment, glutamine, when infused as a labelled tracer, labels/fuels muscular TCA cycle at a much faster rate than that of palmitic acid (ref. ^10^, also this study). Third, as shown in this study, GLS1 activity is directly boosted upon lysosomal AMPK activation, as a rapid response to glucose/FBP decreases, during which no promotion of FAO was observed. Supporting this notion, we also observed that malonyl-CoA in MEFs that inhibits fatty acid oxidation, started to decrease only after 8 h of glucose starvation, possibly owing to a lack of ACC2 phosphorylation when only the lysosomal pool of AMPK is activated under early glucose starvation condition^53^.

It has been a longstanding question how the ER-mitochondria contact is regulated and what physiological roles will follow. Our work demonstrates that, on phosphorylation by AMPK, PDZD8 known to be involved in ER-mitochondria contacts^61^ interacts with mitochondrial GLS1, thereby tightening the ER-mitochondria contact. Based on the results obtained by us and others^87–89^, PDZD8 likely penetrates across the outer mitochondrial membrane (OMM) to interact with the GLS1 localized on the outer side of the inner mitochondrial membrane (IMM) (Extended Data Fig. 10e; see also Supplementary Note 6 for details). Consistently, we have provided evidence that forced ER-mitochondria contact through an ER-mito linker (mAKAP1-mRFP-yUBC6 linker; comprised of the ER-targeting sequence of yUBC6 and mitochondrial membrane-targeting sequence of mAKAP1, see ref. ^90^) is sufficient to promote glutaminolysis even in high glucose (Extended Data Fig. 10f and Supplementary Note 6). Multiple lines of evidence have clearly demonstrated that AMPK mediates the release of the intramolecular autoinhibition of PDZD8, enabling it to interact with GLS1 to enhance ER-mitochondrial contact, which ultimately leads to promotion of GLS1 activity and glutaminolysis. Together, our study reveals an AMPK-PDZD8-GLS1 axis that transmits low glucose–activated AMPK activity to phosphorylation of PDZD8, and to activation of glutaminolysis via increased activity of GLS1. This axis does not only compensate for the reduction of glucose usage, but also elicits mitohormesis that leads to extension of lifespan and healthspan after fasting or caloric restriction.

## Online content

Any methods, additional references, Nature Portfolio reporting summaries, source data, extended data, supplementary information, acknowledgements, details of author contributions and competing interests; and statements of data and code availability are available at https://doi.org/10.1038/.

## Publisher’s note

Springer Nature remains neutral with regard to jurisdictional claims in published maps and institutional affiliations.

## Supporting information

A list of potential AMPK substrates from the purified ER-mitochondria contact (MAM and the MAM-tethered mitochondria) of glucose-starved MEFs

Km and kcat values for GLS1 enzymatic reactions

Summary of lifespan and analysis in worms

Uncropped gel images

Source data and statistical analysis data

## Methods

### Data reporting

The chosen sample sizes were similar to those used in this field: *n* = 3-6 cells, in which 100-400 mitochondria and 100-400 contact sites were contained, were used to determine the formation of ER-mitochondria contacts through TEM^91,92^; *n* = 14-19 mitochondria to determine the formation of ER-mitochondria contacts through FIB-SEM^60^, *n* = 19-103 cells to determine the formation of ER-mitochondria contacts through SPLICS (split-GFP-based contact site sensors) staining^93^, *n* = 3-10 samples to evaluate the levels of metabolites in cells, tissues^10,12,40,53,94^ and nematodes^95–97^; *n* = 3-9 samples to determine OCR in cells, tissues^94,98^ and nematodes^99–101^, and *n* = 6-10 samples to determine mitochondrial ROS in cells^102^, tissues^103^ and nematodes^104,105^; *n* = 3-4 samples to determine the activity of GLS1 (ref. ^72,106,107^); *n* = 3-4 samples to determine the expression levels and phosphorylation levels of a specific protein^41^; *n* = 20-27 cells to determine protein interaction by FRET-FLIM assay in living cells^108^; *n* = 14-17 cells to determine protein interaction by PLA^109,110^; *n* = 3-4 samples to determine the mRNA levels of a specific gene^41^; *n* = 200 worms were used to determine lifespan^111–113^; and *n* = 60 worms were used to determine healthspan^114–116^. No statistical methods were used to predetermine sample size. All experimental findings were repeated as stated in figure legends, and all additional replication attempts were successful. For animal experiments, mice or nematodes were maintained under the same condition or place. For cell experiments, cells of each genotype were cultured in the same CO_2_ incubator and were parallel seeded and randomly assigned to different treatments. Each experiment was designed and performed along with controls, and samples for comparison were collected and analysed under the same conditions. Randomisation was applied wherever possible. For example, during MS analyses (metabolites and proteins), samples were processed and applied to the mass spectrometer in random orders. For animal experiments, sex-matched (only for mice), age-matched litter-mate animals in each genotype were randomly assigned to pharmacological treatments. Otherwise, randomisation was not performed. For example, when performing immunoblotting, samples needed to be loaded in a specific order to generate the final figures. Blinding was applied wherever possible. For example, samples, cages or agar plates during sample collection and processing were labelled as code names that were later revealed by the individual who picked and treated animals or cells, but did not participate in sample collection and processing, until assessing outcome. Similarly, during microscopy data collection and statistical analyses, the fields of view were chosen on a random basis, and are performed by different operators, preventing potentially biased selection for desired phenotypes. Otherwise, blinding was not performed, such as the measurement of GLS1 activity in vitro, as different reagents were added for particular reactions.

### Mouse strains

Protocols for all rodent experiments were approved by the Institutional Animal Care and the Animal Committee of Xiamen University (XMULAC20180028 and XMULAC20220050). Wildtype C57BL/6J mice (#000664) were obtained from The Jackson Laboratory. *AXIN*^F/F^ and *LAMTOR1*^F/F^ mice were generated and validated as described previously^41^. *AMPK*α*1*^F/F^ (#014141) and *AMPK*α*2*^F/F^ mice (#014142) were obtained from Jackson Laboratory, provided by Dr. Sean Morrison. *PDZD8*^−/−^ (KO-first; *Pdzd8^tm1a(EUCOMM)Wtsi^*) mice were obtained from Wellcome Trust Sanger Institute, and *GLS1*^F/F^ mice (#T015195) from GemPharmatech. *AMPK*α*1*/*2*^F/F^ mice were crossed with *Mck*-*Cre* mice to generate muscle-specific knockout (*AMPK*α-MKO) mice (validated in ref. ^94^). To generate *PDZD8*-MKO mice with muscle specific reintroduction of PDZD8 or its 527A mutant, the *PDZD8*^−/−^ mice were first crossed with FLPo mice (036512-UCD; MMRRC) to generate the *PDZD8*^F/F^ mice. Wildtype PDZD8 or its 527A mutant was then introduced to the *PDZD8*^F/F^ mice under the *Rosa26*-LSL(LoxP-Stop-LoxP) system^117^, followed by crossing with *HSA-CreERT2* mice (#025750; Jackson Laboratory). The removal of endogenous PDZD8 and the LSL cassette ahead of introduced PDZD8 and PDZD8-T527A (to trigger the expression of introduced PDZD8) was achieved by intraperitoneally injecting mice with tamoxifen (dissolved in corn oil) at 200 mg/kg, 3 times a week.

To introduce PDZD8 or PDZD8-T527A into *PDZD8*^F/F^ mice, cDNA fragments encoding PDZD8 or PDZD8-T527A were inserted into the *Rosa26*-CTV vector^118^, followed by purification of the plasmids using CsCl density gradient ultracentrifugation method. Some 100 μg of plasmid was then diluted with 500 μl of di-distilled water, followed by concentrating via centrifuge at 14,000*g* at room temperature in a 30-kDa-cutoff filter (UFC503096, Millipore) to 50 μl of solution. The solution was diluted with 450 μl of di-distilled water, followed by another two rounds of dilution/concentration cycles. The plasmid was then mixed with 50 μl of di-distilled water to a final volume of 100 μl, followed by mixing with 10 μl of NaAc solution (3 M stock concentration, pH 5.2). The mixture was then mixed with 275 μl of ethanol, followed by incubating at room temperature for 30 min to precipitate plasmid. The precipitated plasmid was collected by centrifuge at 16,000*g* for 10 min at room temperature, followed by washing with 800 μl of 75% (v/v) ethanol (in di-distilled water) twice. After evaporating ethanol by placing the plasmid next to an alcohol burner lamp for 10 min, plasmid was dissolved in 100 μl of nuclease-free water. The plasmid, along with SpCas9 mRNA and the sgRNAs against the mouse *Rosa26* locus, was then microinjected into the in vitro fertilised (IVF) embryos of the *PDZD8*^F/F^ mice. To generate the SpCas9 mRNA, 1 ng of pcDNA3.3-hCas9 plasmid (constructed by inserting the Cas9 fragment released from Addgene #41815 (ref. ^119^), into the pcDNA3.3 vector; diluted to 1 ng/μl) was amplified using the Phusion High-Fidelity DNA Polymerase kit on a thermocycler (T100, Bio-Rad) with the following programmes: pre-denaturing at 98 °C for 30 s; denaturing at 98 °C for 10 s, annealing at 68 °C for 25 s, then extending at 72 °C for 2 min in each cycle; and final extending at 72 °C for 2 min; cycle number: 33. The following primer pairs were used: 5’-CACCGACTGAGCTCCTTAAG-3’, and 5’-TAGTCAAGCTTCCATGGCTCGA-3’. The PCR product was then purified using the MinElute PCR Purification Kit following the manufacturer’s instructions. The purified SpCas9 PCR product was then subjected to in vitro transcription using the mMESSAGE mMACHINE T7 Transcription Kit following the manufacturer’s instruction (with minor modifications). Briefly, 5.5 μl (300 ng/μl) of SpCas9 PCR product as the template was mixed with 10 μl of 2× NTP/ARCA solution, 2 μl of 10× T7 Reaction Buffer, 0.5 μl of RNase inhibitor, 2 μl of T7 Enzyme Mix, and 4.5 μl of nuclease-free water, followed by incubating at 37 °C for 2 h. The mixture was then mixed with 1 μl of Turbo DNase, followed by incubating at 37 °C for 20 min to digest the template. The mixture was then mixed with 20 μl of 5× *E*-PAP Buffer, 10 μl of 25 mM MnCl_2_, 10 μl of 10 mM ATP, 4 μl of *E*-PAP enzyme, and 36 μl of nuclease-free water, followed by incubating at 37 °C for 20 min for poly(A) tailing. The tailed product was then purified using the MEGAclear Transcription Clean-Up Kit following the manufacturer’s instruction (with minor modifications). Briefly, some 20 μl of tailed RNA was mixed with 20 μl of Elution Solution, followed by mixing with 350 μl of Binding Solution Concentrate. Some 250 μl of ethanol was then added to the mixture, followed by passing the mixture through the Filter Cartridge and washing with 250 μl of Wash Solution twice. The RNA was then eluted with 50 μl of pre-warmed (at 90 °C) Elution Solution. The sgRNAs was prepared as in the SpCas9 mRNA preparation, except that: a) the gRNA Cloning Vector (Addgene, #41824, ref. ^119^) was used as template, and the following programmes: pre-denaturing at 98 °C for 30 s; denaturing at 98 °C for 10 s, annealing at 60 °C for 25 s, then extending at 72 °C for 20 s in each cycle; and final extending at 72 °C for 2 min; cycle number: 33; and following primers: 5’-GAAATTAATACGACTCACTATAGGCGCCCATCTTCTAGAAAGACGTTTTA GAGCTAGAAATAGC-3’, and 5’-AAAAGCACCGACTCGGTGCC-3’; were used; b) in vitro transcription was performed using the MEGAshortscript T7 Transcription Kit, in which the mixture containing: 7.5 μl (100 ng/μl) of purified PCR product, 2 μl of T7 10× T7 Reaction Buffer, 2 μl of T7 ATP solution, 2 μl of T7 CTP solution, 2 μl of T7 GTP solution, 2 μl of T7 UTP solution, 0.5 μl of RNase inhibitor, 2 μl of T7 Enzyme Mix, and 7.5 μl of nuclease-free water; was prepared. In addition, the poly(A) tailing assay was not performed.

To perform IVF on the *PDZD8*^F/F^ mouse strain (according to ref. ^120^, with modifications), the 4-week-old *PDZD8*^F/F^ female mice were intraperitoneally injected with pregnant mare’s serum gonadotrophin (PMSG) at a dose of 10 U/mouse. At 46 h after the PMSG injection, 10 U/mouse human chorionic gonadotrophin (hCG) was intraperitoneally injected. At 12 h after the hCG injection, oocytes from the oviducts of female mice, along with sperms from cauda epididymides and vasa deferentia of 16-week-old, proven stud *PDZD8*^F/F^ male mice, were isolated. To isolate oocytes, oviducts were briefly left on a filter paper, followed by incubating in a human tubal fluid medium (HTF)/GSH drop on an IVF dish (prepared by placing 200 μl of HTF solution supplemented with 125 mM GSH on a 35-mm dish to form a drop, followed by covering the drop with mineral oil and pre-balancing in a humidified incubator containing 5% CO_2_ at 37 °C for 0.5 h before use). The ampulla was then teared down by forceps, and the cumulus oocyte masses inside was collected and transferred to another HTF/GSH drop. To isolate sperms, cauda epididymides and vasa deferentia were briefly left on a filter paper, followed by penetrating with a 26 G needle on the cauda epididymides for 5 times. Sperms were then released to an HTF drop on sperm capacitation dish (prepared by placing 200 μl of HTF solution on a 35-mm dish to form a drop, followed by covering the drop with mineral oil and pre-balancing in a humidified incubator containing 5% CO_2_ at 37 °C for 12 h before use) by slightly pressing/squeezing the cauda epididymides, followed by incubating in a humidified incubator containing 5% CO_2_ at 37 °C for 0.5 h. The capacitated, motile sperms (located on the edge of each HTF drop) were then collected, followed by adding to the oocyte masses soaked in the HTF/GSH drop, 8 μl per drop. The IVF dishes containing oocyte masses and sperms were then cultured in a humidified incubator containing 5% CO_2_ at 37 °C for 4 h, followed by collecting and washing oocytes in a KSOM drop (freshly prepared by placing 20 μl of KSOM medium on a 35-mm dish to form a drop, followed by covering the drop with mineral oil and pre-balancing in a humidified incubator containing 5% CO_2_ at 37 °C for 0.5 h) twice. The oocytes were then cultured in an HTF/GSH drop on an IVF dish for another 12 h in a humidified incubator containing 5% CO_2_ at 37 °C. The presumptive zygotes (in which 2 pronuclei and an extruded, second polar body could be observed) were then picked up. Some 10 pl of DNA mixture comprising *Rosa26*-CTV-PDZD8 plasmid (20 ng/μl final concentration), SpCas9 mRNA (120 ng/μl final concentration) and *Rosa26* sgRNA (100 ng/μl), was microinjected into each of the zygote, and were cultured in KSOM medium at 37 °C in a humidified incubator containing 5% CO_2_. At 16 h of culturing, the zygotes/embryos at two-cell stage were picked up and transplanted into pseudopregnant ICR female mice (8-10 weeks old, >26 g; prepared by breeding the in-estrus female with a 14-week-old, vasectomised male at a day before the transplantation), 20 zygotes/embryos per mouse, and the offspring carrying the LSL-PDZD8 or LSL-PDZD8-527A allele were further outcrossed 6 times to C57BL/6 mice before experiments.

The *PDZD8*-MKO mice with muscle specific reintroduction of PDZD8 or its 527A mutant were validated as depicted in Extended Data Fig. 5l. For genotyping *Rosa26* locus, the following programmes: pre-denaturing at 98 °C for 300 s; denaturing at 95 °C for 30 s, annealing at 64 °C for 30 s, then extending at 72 °C for 45 s in each cycle for 5 cycles; denaturing at 95 °C for 30 s, annealing at 61 °C for 30 s, then extending at 72 °C for 45 s in each cycle for 5 cycles; denaturing at 95 °C for 30 s, annealing at 58 °C for 30 s, then extending at 72 °C for 45 s in each cycle for 5 cycles; denaturing at 95 °C for 30 s, annealing at 55 °C for 30 s, then extending at 72 °C for 45 s in each cycle for 5 cycles; and final extending at 72 °C for 10 min; were used. For genotyping other genes and elements, the following programmes: pre-denaturing at 95 °C for 300 s; denaturing at 95 °C for 30 s, annealing at 58 °C for 40 s, then extending at 72 °C for 30 s in each cycle; and final extending at 72 °C for 10 min; cycle number: 35; were used. The following primers: 5’-CGCATAACGATACCACGATATCAACAAG-3’ (Primer #1) and 5’-CCGCCTACTGCGACTATAGAGATATC-3’ (Primer #2) for cleaved *FRT*; 5’-ATCACGACGCGCTGTATC-3’ (Primer #3) and 5’-ACATCGGGCAAATAATATCG-3’ (Primer #4) for LacZ; 5’-ACTGTCTGTCCTTCCAGGGG-3’ (Primer #5) and 5’-GTGGAAAAGCCAAGAAAGGC-3’ (Primer 5’-CCACCTTCATGAGCTACAACACC-3’ #6) for LoxP; and 5’-AACAGGAACTGGTACAGGGTCTTGG-3’ for FLPo; 5’-CAGGTAGGGCAGGAGTTGG-3’ and 5’-TTTGCCCCCTCCATATAACA-3’ for *HSA*-*Cre*; 5’-AGTGGCCTCTTCCAGAAATG-3’ and 5’-TGCGACTGTGTCTGATTTCC-3’ for the control of *HSA*-*Cre*; 5’-TCTCCCAAAGTCGCTCTGAG-3’, 5’-AAGACCGCGAAGAGTTTGTC-3’, and 5’-ATGCTCTGTCTAGGGGTTGG-3’ for *Rosa26*, 5’-GGAGTTCTATTAAGACGGTTG-3’ and 5’-GTGCTGGGTCTGTTATCTC-3’ for generating PCR products for sequencing T527.

The following ages of mice were used: 1) for analysing AMPK activation: wild-type, and *AMPK*α-MKO mice, 4 weeks old; 2) for analysing glutaminolysis and FAO in the liver and muscle tissues: wild-type, *AMPK*α-MKO, and *PDZD8*-MKO mice with or without wildtype PDZD8 or PDZD8-527A re-introduced, aged 10 weeks; 3) for analysing OCR and ROS in mouse muscles: wild-type, *AMPK*α-MKO, and *PDZD8*-MKO mice with or without wildtype PDZD8 or PDZD8-527A re-introduced, aged 8 weeks; 4) for determining rejuvenating effects of CR: wild-type, and *PDZD8*-MKO mice with or without wildtype PDZD8 or PDZD8-527A re-introduced, aged 32 weeks.

### CR and starvation treatments of mice

Unless stated otherwise, mice were housed with free access to water and standard diet (65% carbohydrate, 11% fat, 24% protein) under specific pathogen-free conditions. The light was on from 8:00 to 20:00, with the temperature kept at 21-24 °C and humidity at 40-70%. Only male mice were used in the study, and male littermate controls were used throughout the study.

For starvation, mice were individually caged for 1 week before each treatment. The diet was withdrawn from the cage at 5 p.m., and mice were sacrificed at desired time points by cervical dislocation. For CR, mice were individually caged for 1 month before treatment, each mouse was fed with 2.5 g of standard diet (70% of ad libitum food intake for a mouse at 4 months old and above) at 5 p.m. at each day.

### Determination of mouse running capacity and grip strength

The maximal running capacity was determined as described previously^121^, with minor modifications. Briefly, mice were trained on Rodent Treadmill NG (UGO Basile, cat. 47300) at 10 m/min for 5 min for 2 days with normal light-dark cycle, and tests were performed during the dark period. Before the experiment, mice were fasted for 2 h. The treadmill was set at 15° incline, and the speed of treadmill was set to increase in a ramp-mode (10 m/min for 10 min followed by an increase to a final speed of 18 m/min within 15 min). Mice were considered to be exhausted, and removed from the treadmill, following the accumulation of 5 or more shocks (0.1 mA) per minute for two consecutive minutes. The distances travelled were recorded as the running capacity.

Grip strength was determined on a grip strength meter (Ugo Basile, cat. 47200) following the protocol described previously^116^. Briefly, the mouse was held by its tail and lowered (“landed”) until all four limbs grasped the T[bar connected to a digital force gauge. The mouse was further lowered to the extent that the body was horizontal to the apparatus, and was then slowly, steady drawn away from the T[bar until all four limbs were removed from the bar, which gave rise to the peak force in grams. Each mouse was repeated 5 times with 5 min intervals between measurements.

### *Caenorhabditis elegans* strains

Nematodes (hermaphrodites) were maintained on NGM plates spread with *E. coli* OP50 as standard food. All worms were cultured at 20 °C. Wildtype (N2 Bristol) and CA-*aak2* (AGD467; ref. ^122^) strains were obtained from *Caenorhabditis* Genetics Center, and *glna*-*1* (tm6647) and *glna*-*3* (tm8446) from National BioResource Project (NBRP). All mutant strains were outcrossed 6 times to N2 before the experiments. Unless stated otherwise, worms were maintained on nematode growth medium (NGM) plates (1.7% (w/v) agar, 0.3% (w/v) NaCl, 0.25% (w/v) bacteriological peptone, 1 mM CaCl_2_, 1 mM MgSO_4_, 25 mM KH_2_PO_4_-K_2_HPO_4_, pH 6.0, 0.02% (w/v) streptomycin and 5 μg/ml cholesterol) spread with *Escherichia coli* OP50 as standard food.

The *glna*-*1*-knockout and *glna*-*3*-knockout strains were crossed to generate a *glna*-*1* and *glna*-*3* double knockout strain (as an example, and similar procedures were applied to generate the CA-*aak2*;*pdzd*-*8*^−/−^ strain). Before crossing, *glna*-*1*-knockout hermaphrodites were synchronised: worms were washed off from agar plates with 15 ml M9 buffer (22.1 mM KH_2_PO_4_, 46.9 mM Na_2_HPO_4_, 85.5 mM NaCl and 1 mM MgSO_4_) supplemented with 0.05% (v/v) Triton X-100 per plate, followed by centrifugation at 1,000*g* for 2 min. The worm sediments were suspended with 6 ml of M9 buffer containing 50% synchronising bleaching solution (by mixing 25 ml of NaClO solution (5% active chlorine), 8.3 ml of 25% (w/v) NaOH and 66.7 ml of M9 buffer, for a total of 100 ml), followed by vigorous shaking for 2 min and centrifugation for 2 min at 1,000*g*. The sediments were washed with 12 ml of M9 buffer twice, then suspended with 6 ml of M9 buffer, followed by rotating at 20 °C, 30 r.p.m. for 12 h. The synchronised worms were then transferred to the NGM plate and cultured to the L4 stage, followed by heat-shocking at 28 °C for 12 h. The heat-shocked worms were then cultured at 20 °C for 4 days, and the males were picked up for mating with *glna*-*1*-knockout hermaphrodites for another 36 h. The mated hermaphrodites were transferred to new NGM plates and allowed to give birth to more *glna*-*1*-knockout males for another 4 days at 20 °C. The *glna*-*1*-knockout males were then picked up and co-cultured with *glna*-*3*-knockout hermaphrodites at a 1:2 ratio (e.g., 4 males and 8 hermaphrodites on a 10-cm NGM plate) for mating for 36 h at 20 °C, and the mated hermaphrodites (*glna*-*3*-knockout) were picked up for culturing for another 2 days. The offsprings were then picked up and were individually cultured on the 35-mm NGM plate, followed by being individually subjected for genotyping after egg-laying (after culturing for approximately 2 days). For genotyping, individual worms were lysed with 5 μl of Single Worm lysis buffer (50 mM HEPES, pH 7.4, 1 mM EGTA, 1 mM MgCl_2_, 100 mM KCl, 10% (v/v) glycerol, 0.05% (v/v) NP-40, 0.5 mM DTT and protease inhibitor cocktail). The lysates were then frozen at −80 °C for 12 h, followed by incubating at 65 °C for 1 h and 95 °C for 15 min on a thermocycler (XP Cycler, Bioer). The lysates were then cooled to room temperature, followed by amplifying genomic DNA on a thermocycler with the following programmes: pre-denaturing at 95 °C for 10 min; denaturing at 95 °C for 10 s, then annealing and extending at 60 °C for 30 s in each cycle; cycle number: 35. The following primer pairs were used for identifying the *glna*-*1*-knockout: 5’-CCTGGACTGGGAATCGTTCA-3’ and 5’-TACAACTGCGAAACACCGAG-3’; and 5’-CCCTCATTATGCGAACGAAC-3’ and 5’-CCCCCAGAAGTAGATAAACG-3’ for identifying the *glna*-*3*-knockout. The offsprings generated from *glna*-*1*- and *glna*-*3*-knockout-assured individuals were then outcrossed six times to the N2 strain.

The *glna*-*2* was then knocked down in the *glna*-*1* and *glna*-*3* double knockout strain following the previously described procedures^123^. Briefly, synchronised worms (around the L1 stage) were placed on the RNAi plates (NGM containing 1 mg/ml IPTG and 50 μg/ml carbenicillin) spread with HT115 *E*. *coli* stains containing RNAi against *glna*-*2* (well L20 on plate II-5 from the Ahringer *C*. *elegans* RNAi Collection) for 2 days. The knockdown efficiency was then examined by determining the levels of *glna*-*2* mRNA by real-time quantitative PCR (qPCR). Approximately 1,000 worms were washed off from an RNAi plate with 15 ml of M9 buffer containing Triton X-100, followed by centrifugation for 2 min at 1,000*g*. The sediment was then washed with 1 ml of M9 buffer twice, and then lysed with 1 ml of TRIzol. The worms were then frozen in liquid nitrogen, thawed at room temperature and then subjected to repeated freeze-thaw for another two times. The worm lysates were then placed at room temperature for 5 min, then mixed with 0.2 ml of chloroform followed by vigorous shaking for 15 s. After 3 min, lysates were centrifuged at 20,000*g* at 4 °C for 15 min, and 450 μl of the aqueous phase (upper layer) was transferred to a new RNase-free centrifuge tube (Biopur, Eppendorf), followed by mixing with 450 μl of isopropanol, then centrifuged at 20,000*g* at 4 °C for 10 min. The sediments were washed with 1 ml of 75% ethanol (v/v) followed by centrifugation at 20,000*g* for 10 min, and then with 1 ml of anhydrous ethanol followed by centrifugation at 20,000*g* for 10 min. The sediments were then dissolved with 20 μl of RNase-free water after the ethanol was evaporated. The dissolved RNA was then reverse-transcribed to cDNA using ReverTra Ace qPCR RT master mix with a gDNA Remover kit, followed by performing real-time qPCR using Maxima SYBR Green/ROX qPCR master mix on a CFX96 thermocycler (Bio-Rad) with the programmes as described in genotyping the *glna*-knockout strain. Data were analysed using CFX Manager software (v.3.1, Bio-Rad). Knockdown efficiency was evaluated according to the CT value obtained. The primers for *glna-2* are 5’-ACTGTTGATGGTCAAAGGGCA-3’ and 5’-CTTGGCTCCTGCCCAACATA-3’. The primers for *ama-1* (the internal control) are 5’-GACATTTGGCACTGCTTTGT-3’ and 5’-ACGATTGATTCCATGTCTCG-3’.

The *pdzd-8*^−/−^ nematode strains expressing human PDZD8 or its T527A mutant were established as described previously^123^, with minor modifications: a) PDZD8-WT or its T527A mutant was first introduced to the N2 strain; b) such generated strains were then subjected to knockout of the *pdzd8* gene; and c) the *pdzd8*-knockout worms were then picked up for the further outcrossing with N2 strain. Briefly, to generate N2 strain expressing PDZD8 or its T527A mutant, cDNA of PDZD8 or PDZD8-T527A mutant was inserted into a pJM1 vector, with GFP as a selection marker, between the *Nhe* I and *Kpn* I sites (expressed under control by a *sur*-*5* promoter), then injected into the syncytial gonad of the worm (200 ng/µl, 0.5 µl per worm). The injected worms were then recovered on a NGM plate for 2 days, and the F_1_ GFP-expressing hermaphrodites were selected for further culture. The extrachromosomally existed PDZD8 or PDZD8-T527A expression plasmid was then integrated into the nematode genome using UV irradiation to establish nonmosaic transgenic strains as described previously^124^, with minor modifications. Briefly, 70 PDZD8 or PDZD8-T527A mutant-expressing worms at L4 stage were picked up and incubated with 600 µl of M9 buffer, followed by adding 10 µl of TMP solution (3 mg/ml stock concentration in DMSO) and rotating at 30 r.p.m. for 15 min in the dark. Worms were then transferred to a 10-cm NGM plate without OP50 bacteria in the dark, followed by irradiating with UV at a total dose of 35 J/cm^2^ (exposed within 35 s) on a UV crosslinker (CL-508; UVITEC). The irradiated worms were fed with 1 ml of OP50 bacteria at 10^13^/ml concentration, and then cultured at 20 °C for 5 h in the dark, followed by individually cultured on 35-mm NGM plate for 1 week without transferring to any new NGM plate (to make sure that F_1_ was under starvation before further selection). The F_1_ GFP-expressing hermaphrodites were selected and individually cultured for another 2 days, and those F_2_ with 100% GFP-expressing hermaphrodite were selected for further culture. The genomic sequence encoding *pdzd-8* was then knocked out from this strain by injecting a mixture of a pDD122 (Peft-3::Cas9 + ttTi5605 sgRNA) vector carrying sgRNAs against *pdzd-8* (5’-GAGGATCGTATCCAGCATGG-3’, and 5’-GTGAGCACGAAGAAGCGTTG-3’, designed using the CHOPCHOP website http://chopchop.cbu.uib.no/), into young adult worms. The F_1_ hermaphrodite worms were individually cultured on an NGM plate. After egg-laying, worms were lysed using Single Worm lysis buffer, followed by PCR with the programmes as described in genotyping the *glna*-knockout strain, except that the primers 5’-ATCTCCACCACAAACATCACCT-3’ and 5’-CTTCAAAATGCTCGTCAGAGTG-3’ were used. The offsprings generated from knockout-assured individuals were outcrossed six times to the N2 strain, and the expression levels of PDZD8 or PDZD8-T527A were examined by immunoblotting. Strains expressing PDZD8 or PDZD8-T527A at similar levels were chosen for further experiments.

For all nematode experiment, worms at L4 stage were used, except those for CR at 3 days after L4.

### Evaluation of nematode lifespan and healthspan

To determine the lifespan of nematodes, the synchronised worms were cultured to L4 stage before transfer to desired agar plates for determining lifespan. For 2-DG treatment, 4 mM 2-DG (final concentration, and same hereafter) was freshly dissolved in water and was added to warm NGM supplemented with 1.7% (w/v) agar before pouring to make the NGM plates. The plates were stored at 20 °C. For CR, OP50 bacteria were diluted to the concentration of 10^9^/ml (along with 10^12^/ml as the control, ad libitum fed group; see ref. ^80^). The diluted bacteria were isopycnically spread on the NGM plates (for a 35-mm NGM plate, 250 μl of bacteria were used) containing 50 mg/l ampicillin and 50 mg/l kanamycin. Worms were transferred to new plates every 2 d. Live and dead worms were counted during the transfer. Worms that displayed no movement upon gentle touching with a platinum picker were judged as dead. Kaplan-Meier curves were graphed by Prism 9 (GraphPad Software), and the statistical analysis data by SPSS 27.0 (IBM).

Pharyngeal pumping rates, assessed as the numbers of contraction-relaxation cycles of the terminal bulb on nematode pharynx within 1 min, were determined as described previously^125^, with minor modification. Briefly, worms were treated with 2-DG or subjected to CR for 2 days, followed by being picked and placed on a new NGM plate containing *E*. *coli*. After 10 min of incubation at room temperature, the contraction-relaxation cycles of the terminal bulb of each worm were recorded on a stereomicroscope (M165 FC, Leica) through a 63× objective for a consecutive 4 min using the Capture software (v.2021.1.13, Capture Visualisation), and the average contraction-relaxation cycles per min were calculated using the Aimersoft Video Editor software (v.3.6.2.0, Aimersoft).

The resistance of nematodes to the oxidative stress was determined as described previously^114^. Briefly, worms were treated with 2-DG or subjected to CR for 2 days. Some 20 worms were then transferred to an NGM plate containing 15 mM FeSO_4_. Worms were then cultured at 20 °C on such a plate, during which the live and dead worms were counted at every 1 h.

### Determination of mRNA levels of antioxidative genes in nematodes

Levels of antioxidative gene expression were determined through the RNA-sequencing performed by Seqhealth Technology Co., Ltd. (Wuhan, China). Briefly, RNAs from approximately 1,000 worms (treated with 2-DG, or undergone CR) were extracted as described in the section of determining the knockdown efficiency of *glna*-*2*. The residual DNA in each sample was removed by treating with RNase-free DNase I for 30 min at 37 °C, and the quality of RNA was double-checked through agarose gel (1.5%) electrophoresis and the NanoDrop OneC Microvolume UV-Vis Spectrophotometer (Thermo), followed by quantified on a Qubit 3 Fluorometer after staining with the Qubit RNA BR kit. Some 2 µg of total RNAs were then subjected for the construction of cDNA libraries using the Collibri Stranded RNA Library Prep Kit for Illumin Systems following manufacturer’s instruction. The cDNAs in the library with the length 200-500 bps were enriched using KAPA HyperPure magnetic beads following the manufacturer’s instructions, followed by quantification using the Collibri Library Quantification Kit, and sequenced on a DNBSEQ-500 sequencer (MGI Tech Co., Ltd.) under the PEI150 mode. The low-quality sequences, including a) reads containing more than 50% bases with quality lower than 20 in a sequence; b) reads with more than 5% bases unknown; and c) reads containing adaptor sequences were removed from the total reads using the Trimmomatic (version 0.36) software as described previously^126^.

Expression levels of antioxidative gene were quantified through their RPKM (reads per kilobase of transcript per million reads mapped) values. To acquire the RPKM value of each gene, reads were first mapped to the reference sequence of *C*. *elegans* using the STAR software (version 2.5.3a) as described previously^127^ to make sure that reads could be uniquely mapped to the gene chosen to calculate the RPKM values. For genes with more than one alternative transcript, the longest transcript was selected to calculate the RPKM. The RPKM was calculated by the featureCounts software (version 1.5.1) as described previously^128^. RPKM values for each antioxidative gene were plotted using Prism 9 (GraphPad) software.

### Reagents

Rabbit polyclonal antibody against p-T527-PDZD8 (1:1,1000 dilution for immunoblotting (IB)) was raised using the peptide CPLSHSPKRTP(p-T)TLSI (aa 511-525 of human PDZD8) conjugated to the KLH immunogen (linked to the cysteine residue). A rabbit was then biweekly immunised with 100 µg of KLH-conjugated antigen, which is pre-incubated with 500 µg manganese adjuvant (kindly provided by Dr. Zhengfan Jiang from Peking University, see ref. ^129^) for 5 min and then mixed with PBS to a total volume of 500 µl, for 4 times, followed by collecting antiserum. The p-T527-PDZD8 antibody was then purified from the antiserum using the CPLSHSPKRTP(p-T)TLSI peptide-conjugated SulfoLink Coupling resin/column supplied in the SulfoLink Immobilization Kit. To prepare the column, 1 mg of peptide was first dissolved with 2 ml of Coupling Buffer, followed by addition of 0.1 ml of TCEP (25 mM stock concentration) and then incubation at room temperature for 30 min. The mixture was then incubated with SulfoLink Resin in a column, which is pre-calibrated by 2 ml of Coupling Buffer for 2 times, on a rotator at room temperature for 15 min, followed by incubating at room temperature for 30 min without rotating. The excess peptide was then removed, and the resin was washed with 2 ml of Wash Solution for 3 times, followed by 2 ml of Coupling Buffer 2 times. The nonspecific-binding sites on the resin was then blocked by incubating with 2 ml of cysteine solution (by dissolving 15.8 mg of L-cysteine-HCl in 2 ml of Coupling Buffer to make a concentration of 50 mM cysteine) on a rotator for 15 min at room temperature, followed by incubating for another 30 min without rotating at room temperature. After removing the cysteine solution, the resin was washed with 6 ml of Binding/Wash Buffer, followed by incubating with 2 ml of antiserum mixed with 0.2 ml of Binding/Wash Buffer for 2 h on a rotator. The resin was then washed with 1 ml of Binding/Wash Buffer for 5 times, and the antibody was eluted with 2 ml of Elution Buffer. The eluent was then mixed with 100 μl of Neutralization Buffer. The antibody against basal PDZD8 exists in the crude antibody eluent was then removed through a previously described membrane-based affinity purification method^130^. Briefly, the bacterially purified, GST-tagged PDZD8-511-525 was subjected to SDS-PAGE, followed by transferring to a PVDF membrane. The PDZD8-bound-membrane was incubated in 5% (w/v) non-fat milk dissolved in TBST (40 mM Tris, 275 μM NaCl, 0.2% (v/v) Tween-20, pH 7.6) for 2 h, then incubated with the crude antibody preparation for 2 days, and then repeated for another 2 times. Antibody was validated for immunoblotting as shown in Extended Data Fig. 3b.

Rabbit anti-phospho-AMPKα-Thr172 (cat. #2535, RRID: AB_331250; 1:1,000 for IB), anti-AMPKα (cat. #2532, RRID: AB_330331; 1:1,000 for IB), anti-phospho-AMPK substrate motif (cat. #5759, RRID: AB_10949320; 1:1,000 for IB and 1:25 for immunoprecipitation (IP)) anti-phospho-ACC-Ser79 (cat. #3661, RRID: AB_330337; 1:1,000 for IB), anti-ACC (cat. #3662, RRID: AB_2219400; 1:1,000 for IB), anti-cytochrome C (cat. #4280, RRID: AB_10695410; 1:500 for IB), anti-PDI (cat. #3501, RRID: AB_2156433; 1:1,000 for IB), anti-calreticulin (cat. #12238, RRID: AB_2688013; 1:1,000 for IB), anti-erlin2 (cat. #2959, RRID: AB_2277907; 1:1,000 for IB), anti-PDH (cat. #3205, RRID: AB_2277907; 1:1,000 for IB), anti-COXIV (cat. #4850, RRID: AB_2085424; 1:1,000 for IB); anti-GST-tag (cat. #2625, RRID: AB_490796; 1:4,000 for IB), anti-His-tag (cat. #12698, RRID: AB_2744546; 1:1,000 for IB), anti-Myc-tag (cat. #2278, RRID: AB_490778; 1:120 for immunofluorescence (IF)), HRP-conjugated mouse anti-rabbit IgG (conformation-specific, cat. #5127, RRID: AB_10892860; 1:2,000 for IB), HRP-conjugated goat anti-rat IgG (conformation-specific, cat. #98164; 1:2,000 for IB) and mouse anti-Myc-tag (cat. #2276, RRID: AB_331783; 1:500 for IB) were purchased from Cell Signaling Technology. Rabbit anti-calnexin (cat. ab22595, RRID: AB_2069006; 1:1,000 for IB), anti-transferrin (cat. ab1223, RRID: AB_298951; 1:500 for IB), anti-GLS1 (ab202027; 1:120 for IF), and mouse anti-CPT1α (cat. ab128568, RRID: AB_11141632; 1:1,000 for IB), mouse anti-total oxidative phosphorylation (OXPHOS) complex (ab110413, RRID: AB_2629281; 1:1,000 for IB) antibodies were purchased from Abcam. Rabbit anti-PDZD8 (cat. NBP2-58671; 1:1,000 for IB; validated in Extended Data Fig. 2b) was purchased from Novus Biologicals. Mouse anti-ASCL4 (also known as FACL4; cat. sc-365230, RRID: AB_10843105; 1:1,000 for IB) and anti-HA-tag (cat. sc-7392, RRID: AB_2894930; 1:1,000 for IB, 1:500 for IP or 1:120 for IF) antibodies were purchased from Santa Cruz Biotechnology. Rabbit anti-GLS1 (KGA and GAC; cat. 12855-1-AP, RRID: AB_2110381; 1:2,000 for IB and 1:100 for IP), anti-TOMM20 (cat. 11802-1-AP, RRID: AB_2207530; 1:1,000 for IB), anti-PDK4 (cat. 12949-1-AP, RRID: AB_2161499; 1:1,000 for IB), anti-CPT1β (cat. 22170-1-AP, RRID: AB_2713959; 1:1,000 for IB), anti-PDH E1 alpha (PDHA1; cat. 18068-1-AP, RRID: AB_2162931; 1:5,000 for IB), anti-tubulin (cat. 10068-1-AP, RRID: AB_2303998; 1:1,000 for IB nematode tubulin), and mouse anti-tubulin (cat. 66031-1-Ig, RRID: AB_11042766; 1:20,000 for IB mammalian tubulin) antibodies were purchased from Proteintech. Rabbit anti-APEX2 (cat. PA5-72607; 1:1,000 for IB) antibody was purchased from Thermo Scientific. Mouse anti-FLAG M2 (cat. F1804, RRID: AB_262044; 1:1,000 for IB) antibody was purchased from Sigma. Rabbit anti-RMDN3 (also known as PTPIP51; cat. A5820, RRID: AB_2766572; 1:1,000 for IB) antibody was purchased from Abclonal. The horseradish peroxidase (HRP)-conjugated goat anti-mouse IgG (cat. 115-035-003, RRID: AB_10015289; 1:5,000 dilution for IB) and goat anti-rabbit IgG (cat. 111-035-003, RRID: AB_2313567; 1:5,000 dilution for IB and 1:120 dilution for IHC) antibodies were purchased from Jackson ImmunoResearch.

Glucose (cat. G7021), DMSO (cat. D2650), PBS (cat. P5493), NaCl (cat. S7653), KCl (cat. P9333), NaOH (cat. S8045), HCl (cat. 320331), ATP (disodium salt; cat. A6419), ATP (magnesium salt, for kinase assay; cat. A9187), agar (cat. A1296), SDS (cat. 436143), CaCl_2_ (cat. C5670), MgSO_4_ (cat. M2643), KH_2_PO_4_ (cat. P5655), K_2_HPO_4_ (cat. P9666), cholesterol (cat. C3045), Na_2_HPO_4_ (cat. S7907), NaH_2_PO_4_ (cat. S8282), sodium hypochlorite solution (NaClO; cat. 239305), HEPES (cat. H4034), MES (cat. 69889), EDTA (cat. E6758), EGTA (cat. E3889), MgCl_2_ (cat. M8266), CsCl (cat. 289329), NaAc (cat. S7670), ethanol (cat. 459836), isopropanol (cat. 34863), KCl (cat. P9333), glycerol (cat. G5516), IGEPAL CA-630 (NP-40, cat. I3021), Triton X-100 (cat. T9284), Tween-20 (cat. P9416), cholesteryl hemisuccinate (CHS; cat. C6512), sodium deoxycholate (cat. S1827), dithiothreitol (DTT; cat. 43815), IPTG (cat. I6758), carbenicillin (cat. C1613), nuclease-free water (for IVF; cat. W4502), L-glutathione reduced (GSH; cat. G4251), mineral oil (cat. M5310 for IVF, and cat. M5904 for CsCl density gradient), streptomycin (for nematode culture; cat. 85886), Trioxsalen (TMP; cat. T6137), 2-deoxy-D-glucose (2-DG; cat. D8375), ampicillin (cat. A9518), kanamycin (cat. E004000), iron(II) sulfate heptahydrate (FeSO_4_; cat. F8633), agarose (cat. A9539), biotinyl tyramide (biotin-phenol; cat. SML2135), Trizma base (Tris; cat. T1503), hexadimethrine bromide (polybrene; cat. H9268), sodium pyrophosphate (cat. P8135), β-glycerophosphate (cat. 50020), hydrogen peroxide (H_2_O_2_; cat. H1009), sodium azide (NaN_3_; cat. S2002), sodium ascorbate (cat. A4034), 6-Hydroxy-2,5,7,8-tetramethylchromane-2-carboxylic acid (Trolox; cat. 238813), sodium carbonate (Na_2_CO_3_; cat. S7795), urea (cat. U5378), myristic-d27 acid (cat. 68698), glutamine (cat. G8540), carnitine (cat. C0283), BSA (cat. A2153), fatty acid-free BSA (cat. SRE0098), methoxyamine hydrochloride (cat. 89803), MTBSTFA (with 1% t-BDMCS; cat. M-108), hexane (cat. 34859), pyridine (cat. 270970), sodium palmitate (PA; cat. P9767), methanol (cat. 646377), chloroform (cat. C7559), heparin sodium salt (cat. H3149), acetonitrile (cat. 34888), ammonium acetate (cat. 73594), ammonium hydroxide solution (cat. 338818), LC-MS-grade water (cat. 1153332500), mannitol (cat. M4125), L-methionine sulfone (cat. M0876), D-campher-10-sulfonic acid (cat. 1087520), 3-aminopyrrolidine dihydrochloride (cat. 404624), N,N-diethyl-2-phenylacetamide (cat. 384011), trimesic acid (cat. 482749), diammonium hydrogen phosphate (cat. 1012070500), ammonium trifluoroacetate (cat. 56865), oligomycin A (cat. 75351), FCCP (cat. C2920), antimycin A (cat. A8674), rotenone (cat. R8875), gentamycin (cat. 345814), collagenase A (cat. 11088793001), imidazole (cat. I5513), taurine (cat. T8691), ADP (cat. 01897), phosphocreatine (cat. V900832), leupeptin (L2884), saponin (cat. S4521), lactobionate (cat. L3375), glutamate (cat. G8415), malate (cat. M7397), succinate (cat. S9512), sucrose (cat. S7903), digitonin (cat. D141), sodium pyruvate (for Oxygraph-2k measurement; cat. P5280), formaldehyde solution (formalin; F8775), glutaraldehyde solution (cat. G5882), glycine (cat. G8898), K_3_Fe(CN)_6_ (cat. 455946), thiocarbonohydrazide (cat. 223220), Pb(NO_3_)_2_ (cat. 203580), sodium citrate (cat. 71497), potassium acetate (cat. P1190), magnesium acetate (cat. M5661), MEA (cat. 30070), glucose oxidase (cat. G2133), catalase (cat. C40), ammonium hydroxide solution (cat. 338818), OptiPrep (cat. D1556), Percoll (cat. P4937), Coomassie Brilliant Blue R-250 (cat. 1.12553), chymotrypsin (cat. C3142), formic acid (cat. 5.43804), ammonium formate (cat. 70221), β-mercaptoethanol (cat. M6250), MOPS (cat. M3183), acetic acid (cat. 27225), L-glutamic dehydrogenase (GDH; cat. G2626), NAD^+^ (cat. N3014), BPTES (cat. SML0601), Etomoxir (cat. 236020), human tubal fluid (HTF) medium (cat. MR-070-D), KSOM medium (cat. MR-121-D), triple-free DMEM (cat. D5030), Lysosome Isolation Kit (cat. LYSISO1), Endoplasmic Reticulum Isolation Kit (cat. ER0100), Glutamate Assay Kit (cat. MAK004), anti-FLAG M2 affinity gel (cat. A2220; 1:500 for IP), HIS-Select Nickel Affinity Gel (cat. P6611), and Duolink In Situ Red Starter Kit (Mouse/Rabbit; cat. DUO92101) were purchased from Sigma. Penicillin-streptomycin (for DMEM preparation; cat. 15140163), Phusion High-Fidelity DNA Polymerase kit (cat. F530N), mMESSAGE mMACHINE T7 Transcription Kit (cat. AM1344), MEGAclear Transcription Clean-Up Kit (cat. AM1908), MEGAshortscript T7 Transcription Kit (cat. AM1354), TRIzol (cat. 15596018), UltraPure DNase/RNase-Free Distilled Water (RNase-free water; cat. 10977015), Maxima SYBR Green/ROX qPCR master mix (cat. K0223), RNase-free DNase I (cat. EN0523), Qubit RNA BR assay kit (cat. Q10211), Collibri Stranded RNA Library Prep Kit (cat. A39003024), Collibri Library Quantification Kit (cat. A38524100), SulfoLink Immobilization Kit for Peptides (cat. 44999), DMEM, high glucose (DMEM; cat. 11965175), glucose-free DMEM (cat. 11966025), FBS (cat. 10099141C), Lipofectamine 2000 (cat. 11668500), MEM non-essential amino acids solution (cat. 11140050), GlutaMAX (cat. 35050061), sodium pyruvate (cat. 11360070), ProLong Diamond antifade mountant (cat. P36970), ProLong Live Antifade reagent (cat. P36975), Streptavidin Magnetic Beads (cat. 88817; 1:100 for IP), NeutrAvidin agarose (cat. 29204), MitoSOX (cat. M36008), EZ-Link Sulfo-NHS-SS-Biotin (cat. 21331), and Prestained Protein MW Marker (cat. 26612) were purchased from Thermo Scientific. OsO_4_ (cat. 18465) and uranyl acetate (cat. 19481) were purchased from Tedpella. SPI-Pon 812 Embedding Kit (cat. 02660-AB) was purchased from Structure Probe, Inc. n-dodecyl-β-D-maltopyranoside (DDM; cat. D310) was purchased from Anatrace Products, LLC. Difco LB Broth (cat. 240220) was purchased from BD. Bacteriological peptone (cat. LP0137) was purchased from Oxoid. Seahorse XF base medium (cat. 103334) and Seahorse XF Calibrant solution (cat. 100840) were purchased from Agilent. O.C.T. Compound (cat. 4583) was purchased from Sakura Finetek USA, Inc. Antifade Mounting Medium (cat. H-1000-10) was purchased from Vector Laboratories, Inc. PrimeSTAR HS polymerase (cat. R40A) was purchased from Takara. Polyethylenimine (PEI; cat. 23966) was purchased from Polysciences. Nonfat dry milk (cat. #9999) and normal goat serum (NGS; cat. #5425) were purchased from Cell Signaling Technology. ReverTra Ace qPCR RT master mix with a gDNA Remover kit was purchased from TOYOBO. Protease inhibitor cocktail (cat. 70221) and KAPA HyperPure magnetic beads (cat. KK8010) were purchased from Roche. WesternBright ECL and peroxide solutions (cat. 210414-73) were purchased from Advansta. [U-^13^C]-glutamine (cat. 184161-19-1), [U-^13^C]-palmitate ([U-^13^C]-PA; cat. CLM-3943), [alpha-^15^N]glutamine (cat. NLM-1016), [U-^13^C]-pyruvate (cat. CLM-2440), tryptophan-d5 (cat. DLM-1092), and [U-^13^C]-glucose (CLM-1396) were purchased from Cambridge Isotope Laboratories. The isotope-labelled AMP (cat. 123603801), ADP (cat. 129603601) and ATP (cat. 121603801) standards were purchased from Silantes. 3-hydroxynaphthalene-2,7-disulfonic acid disodium salt (2-naphtol-3,6-disulfonic acid disodium salt; cat. H949580) was purchased from Toronto Research Chemicals. Hexakis(1H,1H,3H-perfluoropropoxy)phosphazene (hexakis(1H, 1H, 3H-tetrafluoropropoxy)phosphazine; cat. sc-263379) was purchased from Santa Cruz Biotechnology. MinElute PCR Purification Kit (cat. 28004) was purchased from Qiagen. hCG and PMSG were purchased from Sansheng Biological Technology Co., Ltd. (Ningbo, China). rProtein A Sepharose Fast Flow (cat. 17127904), Protein G Sepharose 4 Fast Flow (cat. 17061806), Glutathione Sepharose 4 Fast Flow (cat. 17513203), and Superdex 200 Increase 10/300 GL (cat. 28990944) were purchased from Cytiva.

### Plasmids

Full-length cDNAs used in this study were obtained either by PCR using cDNA from MEFs, or by purchasing from Origene or Sino Biological. Mutations of PDZD8 and GLS1 were performed by PCR-based site-directed mutagenesis using PrimeSTAR HS polymerase. Expression plasmids for various epitope-tagged proteins were constructed in the pcDNA3.3 vector for transfection (ectopical expression in mammalian cells), in the pBOBI vector for lentivirus packaging (stable expression in mammalian cells), in the in pLVX-IRES for doxycycline-inducible expression, or in the pET-28a and pGEX4T-1 (bacterial expression) vectors. PCR products were verified by sequencing (Invitrogen, China). The lentivirus-based vector pLV-H1-EF1a-puro (for GLS1) and pLL3.7 (for RMDN3) was used for expression of siRNA in MEFs. All plasmids used in this study were purified by CsCl density gradient ultracentrifugation method.

### Cell lines

In this study, no cell line used is on the list of known misidentified cell lines maintained by the International Cell Line Authentication Committee (https://iclac.org/databases/cross-contaminations/). HEK293T cells (cat. CRL-3216) were purchased from ATCC. HEK293T cells and MEFs were maintained in Dulbecco’s modified Eagle’s medium (DMEM) supplemented with 10% fetal bovine serum (FBS), 100 IU penicillin, 100 mg/ml streptomycin at 37 °C in a humidified incubator containing 5% CO_2_. All cell lines were verified to be free of mycoplasma contamination. HEK293T cells were authenticated by STR sequencing. PEI at a final concentration of 10 μM was used to transfect HEK293T cells. Total DNA to be transfected for each plate was adjusted to the same amount by using relevant empty vector. Transfected cells were harvested at 24 h after transfection.

Lentiviruses, including those for knockdown or stable expression, were packaged in HEK293T cells by transfection using Lipofectamine 2000. At 30 h post transfection, medium (DMEM supplemented with 10% FBS and MEM non-essential amino acids; approximately 2 ml) was collected and centrifuged at 5,000*g* for 3 min at room temperature. The supernatant was mixed with 10 μg/ml polybrene, and was added to MEFs or HEK293T cells, followed by centrifuging at 3000*g* for 30 min at room temperature (spinfection). Cells were incubated for another 24 h (MEFs) or 12 h (HEK293T cells) before further treatments.

*AMPK*α^−/−^MEFs and *AMPK*α^−/−^HEK293T cells were generated and validated as described previously^131^. *LAMTOR1*^F/F^ and *AXIN*^F/F^ MEFs were established by introducing SV40 T antigen via lentivirus into cultured primary embryonic cells from mouse litters as described previously^41^, so does *GLS1*^F/F^ MEFs. *LAMTOR1*^−/−^, *AXIN*^−/−^ and *GLS1*^−/−^ MEFs were generated by infecting each of MEFs with adenoviruses expressing the Cre recombinase (cat. 1045, Vector Biolabs) for 12 h. The infected cells were then incubated in the fresh DMEM for another 12 h before further treatments. The *GLS1* (encoding both *KGA* and *GAC*) gene was knocked down and validated in MEFs as described previously^132^. The sequence of siRNA used to knockdown mouse *RMDN3* is: 5’-GAAGCCGACAAGACTTTCT-3’.

The mouse genes (*PDZD8*, *RMDN3*, *PDHA1*, *CPT1A* and *CPT1B*) were deleted from MEFs using the CRISPR-Cas9 system. Nucleotides were annealed to their complements containing the cloning tag aaac, and inserted into the back-to-back *Bsm*B I restriction sites of lentiCRISPRv2 vector (#52961, Addgene). The sequence for each sgRNA is as follows: 5’-CACCCCTCGGCGCCGCCGCCATAA-3’ for *PDZD8*; 5’-TCTTATGGCGCTGCGGCGCG-3’ 5’-GCTGTATCCCGCGTGTTGGC-3’ for for *RMDN3*; *PDHA1*; 5’-GGCGGAGATCGATGCCATCA-3’ for *CPT1A*; and 5’-TCCACCGGAGTCTGGGCGAC-3’ for *CPT1B*. The constructs were then subjected to lentivirus packaging using HEK293T cells that were transfected with 2 µg of DNA in Lipofectamine 2000 transfection reagent per well of a 6-well plate. At 30 h post transfection, the virus (approximately 2 ml) was collected and for infecting MEFs as described above, except cells cultured to 15% confluence were incubated with virus for 72 h. When cells were approaching to confluence, they were single-cell sorted into 96-well dishes. Clones were expanded and evaluated for knockout status by sequencing.

For glucose starvation, cells were rinsed twice with PBS, and then incubated in glucose-free DMEM supplemented with 10% FBS and 1 mM sodium pyruvate for desired periods of time at 37 °C.

### IP and IB assays

To determine the interaction between endogenous GLS1 and PDZD8, four 10-cm dishes of MEFs (grown to 80% confluence) were collected for each experiment. Cells, starved or unstarved, were lysed with 500 μl per dish of ice-cold DDM/CHS lysis buffer (20 mM HEPES, pH 7.4, 50 mM NaCl, 10 mM MgCl_2_, 0.5% (w/v) DDM, 0.1% (w/v) CHS with protease inhibitor cocktail) without pre-washing with PBS (same hereafter for all IP and IB assays), followed by sonication and centrifugation at 4 °C for 15 min. Cell lysates were incubated with GLS1 or PDZD8 antibody overnight. Overnight protein aggregates were pre-cleared by centrifugation at 20,000*g* for 10 min, and protein A/G beads (1:200 dilution, balanced with DDM/CHS lysis buffer) were then added into the lysate/antibody mixtures for another 3 h at 4 °C. The beads were centrifuged and washed with 100 times the volume of ice-cold DDM/CHS wash buffer (20 mM HEPES, pH 7.4, 50 mM NaCl, 10 mM MgCl_2_, 0.01% (w/v) DDM, 0.002% (w/v) CHS) 3 times (by centrifuging at 2,000*g*) at 4 °C and then mixed with an equal volume of 2× SDS sample buffer and boiled for 10 min before IB.

To determine the interaction between ectopically expressed GLS1 and PDZD8, a 6 cm-dish of HEK293T cells were transfected with different expression plasmids. At 24 h after transfection, cells were collected and lysed in 500 µl of ice-cold DDM/CHS lysis buffer, followed by sonication and centrifugation at 4 °C for 15 min. Anti-HA-tag (1:100) or anti-Myc-tag (1:100) antibodies, along with protein A/G beads (1:100), or anti-FLAG M2 Affinity Gel (1:200, pre-balanced in DDM/CHS lysis buffer) was added into the supernatant and mixed for 4 h at 4 °C. The beads were washed with 200 times volume of DDM/CHS wash buffer for 3 times at 4 °C and then mixed with an equal volume of 2× SDS sample buffer and boiled for 10 min before immunoblotting.

To verify the phosphorylation of MAM proteins (listed in Supplementary Table 1) by AMPK (using the anti-pan-phospho-AMPK substrate antibody), a 10 cm-dish of HEK293T cells was transfected with different expression plasmids. IP was performed as in determining the interaction between ectopically expressed GLS1 and PDZD8, except that ice-cold Triton lysis buffer (20 mM Tris-HCl, pH 7.5, 150 mM NaCl, 1 mM EDTA, 1 mM EGTA, 1% (v/v) Triton X-100, 2.5 mM sodium pyrophosphate, 1 mM β-glycerophosphate, with protease inhibitor cocktail) was used to lyse cells and wash protein A/G beads. In particular, antibodies were incubated with cell lysates for a time duration of 15 min to avoid the possible phosphorylation mediated by AMPK in the lysate (even in the unstarved cells).

The APEX2 proximity labeling assay was performed as described previously^133^, with minor modifications. Briefly, protein biotinylation reactions in 60 10-cm dishes of MEFs with stable expression of APEX2-PDZD8 were treated with DMEM (10 ml per dish) containing 500 mM biotinyl tyramide at 37 °C for 30 min, followed by addition of 1 mM hydrogen peroxide and incubated at room temperature for 1 min. The reactions were then terminated by removing the medium and the addition of ice-cold quenching buffer (10 mM sodium azide, 10 mM sodium ascorbate, 5 mM Trolox, in PBS), 10 ml per dish. Cells were washed with PBS, 10 ml per dish, followed by subcellular fractionation (see “Subcellular fractionation” section). Each fraction was lysed with 500 µl of ice-cold RIPA buffer (50 mM Tris-HCl, pH 7.5, 150 mM NaCl, 1% NP-40, 0.5% sodium deoxycholate, 0.1% SDS, with protease inhibitor cocktail), and was centrifuged at 20,000*g* for 10 min at 4 °C. The supernatant was mixed with 20 μl of Streptavidin Magnetic Beads for 12 h on a rotator at 4 °C, followed by washing with 1 ml of ice-cold RIPA buffer twice. The beads were then washed with ice-cold RIPA buffer supplemented with 1 M KCl once, ice-cold RIPA buffer supplemented with 0.1 M Na_2_CO_3_ once, 2 M ice-cold urea dissolved in 10 mM Tris-HCl, pH 8.0 once, and ice-cold RIPA buffer twice. The beads slurry was then mixed with an equal volume of 2× SDS sample buffer and boiled for 10 min before immunoblotting.

To analyse the levels of p-AMPKα, p-ACC and p-PDZD8 in MEFs, cells grown to 70-80% confluence in a well of a 6-well dish were lysed with 250 μl of ice-cold Triton lysis buffer. The lysates were then centrifuged at 20,000*g* for 10 min at 4 °C and an equal volume of 2× SDS sample buffer was added into the supernatant. Samples were then boiled for 10 min and then directly subjected to immunoblotting. To analyse the levels of p-AMPKα, p-ACC and p-PDZD8 in muscle and liver tissues, mice were anesthetised after indicated treatments. Freshly excised (or freeze-clamped) tissues were immediately lysed with ice-cold Triton lysis buffer (10 μl/mg tissue weight for liver, and 5 μl/mg tissue weight for muscle), followed by homogenisation and centrifugation as described above. The lysates were then mixed with 2× SDS sample buffer, boiled, and subjected to immunoblotting. To analyse the levels of PDZD8 in nematodes, about 150 nematodes cultured on the NGM plate were collected for each sample. Worms were quickly washed with ice-cold M9 buffer containing Triton X-100, and were lysed with 150 μl of ice-cold Triton lysis buffer. The lysates were then mixed with 5× SDS sample buffer, followed by homogenisation and centrifugation as described above, and then boiled before subjected to immunoblotting. All samples were subjected to immunoblotting on the same day of preparation, and any freeze-thaw cycle were avoided.

For immunoblotting, the SDS-polyacrylamide gels were prepared in house as described previously^123^. The thickness of gels used in this study was 1.0 mm. Samples of less than 10 μl were loaded into wells, and the electrophoresis was run at 100 V (by PowerPac HC High-Current Power Supply, Bio-Rad) in a Mini-PROTEAN Tetra Electrophoresis Cell (Bio-Rad). In this study, all samples were resolved on 8% resolving gels, except that PDZD8-CT was run on 10% gels, and those smaller than 35 kDa including TOMM20, cytochrome C, ERLIN2 and some PDZD8 truncations on 15% gels. The resolved proteins were then transferred to the PVDF membrane (0.45 μm, cat. IPVH00010, Merck) as described previously^123^. The PVDF membrane was then blocked by 5% (w/v) BSA (for all antibodies against phosphorylated proteins) or 5% (w/v) non-fat milk (for all antibodies against total proteins) dissolved in TBST for 2 h on an orbital shaker at 60 rpm at room temperature, followed by rinsing with TBST for twice, 5 min each. The PVDF membrane was then incubated with desired primary antibody overnight at 4 °C on an orbital shaker at 60 rpm, followed by rinsing with TBST for three times, 5 min each at room temperature, and then the secondary antibodies for 3 h at room temperature with gentle shaking. The secondary antibody was then removed, and the PVDF membrane was further washed with TBST for 3 times, 5 min each at room temperature. PVDF membrane was incubated in ECL mixture (by mixing equal volumes of ECL solution and Peroxide solution for 5 min), then life with Medical X-Ray Film (FUJIFILM). The films were then developed with X-OMAT MX Developer (Carestream), and X-OMAT MX Fixer and Replenisher solutions (Carestream) on a Medical X-Ray Processor (Carestream) using Developer (Model 002, Carestream). The developed films were scanned using a Perfection V850 Pro scanner (Epson) with an Epson Scan software (v.3.9.3.4), and were cropped using Photoshop 2023 software (Adobe). Levels of total proteins and phosphorylated proteins were analysed on separate gels, and representative immunoblots are shown. Uncropped immunoblots are provided as Supplementary Information Figure 1.

### Determination of rates of glutaminolysis and fatty acid oxidation (FAO)

To determine glutaminolysis and FAO rates, MEFs or mice were labelled respectively with [U-^13^C]-glutamine and [U-^13^C]-PA tracers for desired durations, followed by determination of the levels of TCA cycle intermediates through gas chromatography-mass spectrometry (GC-MS). PA was conjugated to BSA after dissolving in 10% fatty acid-free BSA to a stock concentration of 10 mM before use.

To determine the glutaminolysis rates in MEFs, cells from one 10-cm dish (60–70% confluence) were collected for each measurement. MEFs were glucose-starved for desired periods of time by incubating with triple-free (free of glucose, pyruvate and glutamine) DMEM supplemented with 4 mM glutamine, 1 mM sodium pyruvate, 100 µM PA, 1 mM carnitine (according to ref. ^134^), and 10% FBS. At 20 min before sample collection, cells were incubated with pre-warmed triple-free DMEM supplemented with 3 mM glutamine, 1 mM [U-^13^C]-glutamine, 1 mM sodium pyruvate, 100 µM PA, 1 mM carnitine and 10% FBS. Cells were then lysed with 1 ml of 80% methanol (v/v in water) containing 10 µg/ml myristic-d27 acid as an internal standard (IS), followed with 20 s of vortex. After centrifugation at 15,000*g* for 15 min at 4 °C, 600 µl of each supernatant (aqueous phase) was freeze-dried in a vacuum concentrator (a LABCONCO #7310037 centrifuge connected to a LABCONCO #7460037 cold trap and an EDWARDS nXDS15i pump) at 4 °C for 24 h. The lyophilised samples were then subjected for derivatisation by vortexing for 1 min after mixing each with 50 µl of freshly prepared methoxyamine hydrochloride (20 mg/ml in pyridine), followed by incubation at 4 °C for 1 h. The mixtures were sonicated at 0 °C by bathing in ice slurry for 10 min, and were then incubated at 37 °C for 1.5 h, followed by mixing with 50 µl of MTBSTFA and incubated at 55 °C for 1 h. Before subjecting to GC-MS, samples were centrifuged at 15,000*g* for 10 min, and some 60 μl of each supernatant was loaded into an injection vial (cat. 5182-0714, Agilent; with an insert (cat. HM-1270, Zhejiang Hamag Technology)) equipped with a snap cap (cat. HM-0722, Zhejiang Hamag Technology). GC was performed on an HP-5MS column (30 m × 0.25 mm i.d., 0.25 μm film thickness; cat. 19091S-433; Agilent) using a GC/MSD instrument (7890-5977B, Agilent) as described previously^94^. Briefly, the injector temperature of GC/MSD was set at 260 °C. The column oven temperature was first held at 70 °C for 2 min, then increased to 180 °C at the rate of 7 °C/min, then to 250 °C at 5 °C/min, then to 310 °C at 25 °C/min, where it was held for 15 min. The MSD transfer temperature was 280 °C. The MS quadrupole and source temperature were maintained at 150 °C and 230 °C, respectively. Measurements were performed in both a scan mode (to assure the quality and purity of each TCA cycle intermediate peak) and a selected ion monitoring (SIM) mode (to maximise the sensitivity of GC-MS for quantifying each metabolite/isotopomer). In SIM mode, the fragment ion with m/z values of [M-57] (where M is the molecular mass of each derivatised metabolite, and the loss of the 57-Da facile is attributed to the loss of the *tert*-butyl moiety of the metabolite in the GC of each compound) was set as the quantitative ion. To ensure that all possible isotopomer peaks, including those of naturally occurring isotopes of a specific metabolite (with n carbon atoms), were recorded, the m/z values ranging from [M-57] to [M-57] + n + 1 were included during the data collection. In particular, for pyruvate and α-KG, m/z values from [M-57] to [M-57] + n + 2 were recorded, owing to the oximation of these two metabolites during the derivatisation. The following m/z values were used for each compound: 174, 175, 176, 177 and 178 for pyruvate; 289, 290, 291, 292 and 293 for succinate; 287, 288, 289, 291 and 292 for fumarate; 346, 347, 348, 349, 350, 351 and 352 for α-KG; 419, 420, 421, 422 and 423 for malate; 418, 419, 420, 421 and 422 for aspartate; 432, 433, 434, 435, 436 and 437 for glutamate; 431, 432, 433, 434, 435, 436 for glutamine; and 591, 592, 593, 594, 595, 596 and 597 for citrate. Data were collected using the MassHunter GC/MS Acquisition software (v.B.07.04.2260, Agilent). For quantification, peaks were extracted and integrated using GC-MS MassHunter Workstation Qualitative Analysis software (v.B.07.01SP1, Agilent), and were corrected for naturally occurring isotopes using the IsoCor software^135,136^ with the matrix-based method.

Rates of FAO in MEFs were determined as described above for MEFs, except that MEFs were labelled with 100 µM [U-^13^C]-PA for 12 h before collection. Rates of glutamine deamination were also determined as described above, except that MEFs were labelled with 1 mM [alpha-^15^N]glutamine for 20 min, and the following m/z values were used for each compound: 431, 432 and 433 for glutamine; 432 and 433 for glutamate; 418 and 419 for aspartate; and 260 and 261 for alanine. To determine carbon utilisation in MEFs under glucose starvation, cells were separately labelled with the following isotopic tracers: a) 1 mM [U-^13^C]-glutamine, added to the medium as described above; b) 100 µM [U-^13^C]-PA, added to the medium as described above; c) 1 mM [U-^13^C]-pyruvate, added into the triple-free DMEM supplemented with 4 mM glutamine, 100 µM PA, 1 mM carnitine, 10% FBS, and 25 mM glucose or not (for starvation); d) 25 mM [U-^13^C]-glucose, added into the triple-free DMEM supplemented with 4 mM glutamine, 1 mM pyruvate, 100 µM PA, 1 mM carnitine and 10% FBS; all for 24 h (during which the media containing isotopic tracers were refreshed for 4 times, i.e., at 6, 12, 18 and 22 h after labelling) to make sure the isotopic enrichment has reached steady states (been saturated; see ref. ^137,138^ for glutamine, glucose and pyruvate labelling, and ref. ^134^ for PA labelling).

To determine the rates of glutaminolysis and FAO in mouse tissues, mice were cannulated on their right jugular veins to establish a catheter for tracer infusion at 24 h before the experiment^139,140^. Mice were then starved for desired durations, followed by infusion with 6.87 mg/ml [U-^13^C]-glutamine and 2 mM [U-^13^C]-PA (both dissolved in 2.9 mg/ml heparin sodium salt), respectively, both at 3.3 µl/min for 2 h (titrated according to ref. ^12,141^, to achieve a pre-steady-state). At the end of the infusion, the mouse was anaesthetised, and 20 µl of serum, along with 100 mg of liver and muscle tissues, was collected by freeze clamping, immediately followed by freezing in liquid nitrogen. Metabolites of serum and tissues were extracted, followed by subjecting to GC-MS analysis as described above. The levels of each ^13^C-labelled metabolite were then normalised to the levels of [U-^13^C]-glutamine or [U-^13^C]-PA tracer detected in serum (see ref. ^10^). To determine the rates of glutaminolysis in *C*. *elegans*, 1,000 nematodes were incubated with 8 mM [U-^13^C]-glutamine (final concentration, added to a 6-cm NGM plate containing OP50 bacteria) for 24 h. Worms were then washed and collected with M9 buffer, followed by extraction and analysis of metabolites as described above.

### Determination of NAD^+^, malonyl-CoA and glutamine

To determine levels of NAD^+^ and malonyl-CoA, high-performance liquid chromatography-mass spectrometry (HPLC-MS) was performed^94^. Briefly, some 50 mg of fleshly excised (using a freeze-clamp) muscle tissue was immediately frozen in liquid nitrogen, and homogenised in 1 ml of ice-cold methanol. For cells, MEFs collected from one 10-cm dish (grown to 60–70% confluence) were frozen in liquid nitrogen and lysed in 1 ml of ice-cold methanol. The lysates were then mixed with 1 ml of chloroform and 400 µl of water (containing 4 µg/ml [U-^13^C]-glutamine as an IS), followed with 20 s of vortexing. After centrifugation at 15,000*g* for another 15 min at 4 °C, 800 µl of aqueous phase was collected, lyophilised in a vacuum concentrator at 4 °C for 8 h, and then dissolved in 30 µl of 50% (v/v, in water) acetonitrile, followed by loading 20 µl of solution into an injection vial (cat. 5182-0714, Agilent; with an insert (cat. HM-1270, Zhejiang Hamag Technology)) equipped with a snap cap (cat. HM-2076, Zhejiang Hamag Technology). Measurements were based on ref. ^142^ using a QTRAP MS (QTRAP 5500, SCIEX) interfaced with a UPLC system (ExionLC AD, SCIEX). Some 2 µl of each sample were loaded onto a HILIC column (ZIC-pHILIC, 5 μm, 2.1 × 100 mm, PN: 1.50462.0001, Millipore). The mobile phase consisted of 15 mM ammonium acetate containing 3 ml/l ammonium hydroxide (>28%, v/v) in the LC-MS grade water (mobile phase A) and LC-MS grade 90% (v/v) acetonitrile in LC-MS grade water (mobile phase B) run at a flow rate of 0.2 ml/min. Metabolites were separated with the following HPLC gradient elution programme: 95% B held for 2 min, then to 45% B in 13 min, held for 3 min, and then back to 95% B for 4 min. The mass spectrometer was run on a Turbo V ion source in negative mode with a spray voltage of −4,500 V for NAD^+^, or in positive mode with a spray voltage of 5,500 V for malonyl-CoA. Source temperature was set at 550 °C, Gas No.1 at 50 psi, Gas No.2 at 55 psi, and curtain gas at 40 psi. Metabolites were measured using the multiple reactions monitoring mode (MRM), and declustering potentials and collision energies were optimised through using analytical standards. The following transitions (parent ion/daughter ion) were used for monitoring each compound: 662.0/540.1 for NAD^+^, 854/347 for malonyl-CoA, and 149.9/114 (negative mode) or 152/88 (positive mode) for [U-^13^C]-glutamine. Data were collected using Analyst software (v.1.7.1, SCIEX), and the relative amounts of metabolites were analysed using MultiQuant software (v.3.0.3, SCIEX).

To determine the intracellular glutamine levels, samples were prepared as described above, except that tryptophan-d5 (168.9/107.9 on negative mode) at 1 µg/ml final concentration was chosen as an IS. The [U-^13^C]-glutamine dissolved in individual lysates were used to generate corresponding standard curves by plotting the ratios of labelled glutamine (areas) to IS, against the added concentrations of labelled glutamine. The amounts of intracellular glutamine were estimated according to standard curves. The average cell volume of 2,000 μm^3^, as described previously^53^.

### Measurements of adenylates

Levels of AMP, ADP and ATP were analysed by capillary electrophoresis-based mass spectrometry (CE-MS) as described previously^40^, with minor modifications. Briefly, each measurement required MEFs collected from one 10-cm dish (60-70% confluence). Cells were washed with 25 ml of 5% (m/v) mannitol solution (dissolved in water), and were instantly frozen in liquid nitrogen. After thawing, cells were then lysed with 1 ml of methanol containing IS1 (50 µM L-methionine sulfone, 50 µM D-campher-10-sulfonic acid, dissolved in water; 1:500 (v/v) added to the methanol and used to standardise the metabolite intensity and to adjust the migration time). The lysate was then mixed with 1 ml of chloroform and 400 μl of water, followed by 20 s of vortexing. After centrifugation at 15,000*g* for 15 min at 4 °C, 450 μl of aqueous phase was collected and was then filtrated through a 5-kDa cutoff filter (cat. OD003C34, PALL) by centrifuging at 12,000*g* for 3 h at 4 °C. In parallel, quality control samples were prepared by combining 10 μl of the aqueous phase from each sample and then filtered alongside the samples. The filtered aqueous phase was then freeze-dried in a vacuum concentrator at 4 °C, and then dissolved in 100 μl of water containing IS2 (50 µM 3-aminopyrrolidine dihydrochloride, 50 µM N,N-diethyl-2-phenylacetamide, 50 µM trimesic acid, 50 µM 2-naphtol-3,6-disulfonic acid disodium salt, dissolved in methanol; used to adjust the migration time). A total of 20 μl of re-dissolved solution was then loaded into an injection vial (cat. 9301-0978, Agilent; equipped with a snap cap (cat. 5042-6491, Agilent)). Before CE-MS analysis, the fused-silica capillary (cat. TSP050375, i.d. 50 µm × 80 cm; Polymicro Technologies) was installed in a CE/MS cassette (cat. G1603A, Agilent) on the CE system (Agilent Technologies 7100). The capillary was then pre-conditioned with Conditioning Buffer (25 mM ammonium acetate, 75 mM diammonium hydrogen phosphate, pH 8.5) for 30 min, followed by balanced with Running Buffer (50 mM ammonium acetate, pH 8.5; freshly prepared) for another 1 h. CE-MS analysis was run at anion mode, during which the capillary was washed by Conditioning Buffer, followed by injection of the samples at a pressure of 50 mbar for 25 s, and then separation with a constant voltage at −30 kV for another 40 min. Sheath Liquid (0.1 μM hexakis(1H, 1H, 3H-tetrafluoropropoxy)phosphazine, 10 μM ammonium trifluoroacetate, dissolved in methanol/water (50% v/v); freshly prepared) was flowed at 1 ml/min through a 1:100 flow splitter (Agilent Technologies 1260 Infinity II; actual flow rate to the MS: 10 μl/min) throughout each run. The parameters of mass spectrometer (Agilent Technologies 6545) were set as: a) ion source: Dual AJS ESI; b) polarity: negative; c) nozzle voltage: 2,000 V; d) fragmentor voltage: 110 V; e) skimmer voltage: 50 V; f) OCT RFV: 500 V; g) drying gas (N_2_) flow rate: 7 L/min; h) drying gas (N_2_) temperature: 300 °C; i) nebulizer gas pressure: 8 psig; j) sheath gas temperature: 125 °C; k) sheath gas (N_2_) flow rate: 4 L/min; l) capillary voltage (applied onto the sprayer): 3,500 V; m) reference (lock) masses: m/z 1,033.988109 for hexakis(1H, 1H, 3H-tetrafluoropropoxy)phosphazine, and m/z 112.985587 for trifluoroacetic acid; n) scanning range: 50-1,100 m/z; and n) scanning rate: 1.5 spectra/s. Data were collected using MassHunter LC/MS acquisition 10.1.48 (Agilent), and were processed using Qualitative Analysis B.06.00 (Agilent). The peak areas of adenylates were calculated using following parameters (m/z, retention time (min)): a) AMP: 346.0558, 9.302; b) ADP: 426.0221, 10.930; and c) ATP: 505.9885, 11.848. Note that the retention time of each adenylate may vary between each run, and can be adjusted by isotope-labelled standards (dissolved in individual cell or tissue lysates) run between each samples, so do IS1 and IS2.

### Determination of the interaction interface between GLS1 and PDZD8

The interface between GLS1 and PDZD8 was determined through in silico docking using the FRODOCK 2.0 protein docking server (https://frodock.iqfr.csic.es/)143. The reported GAC structure (PDB ID: 3UO9, ref. ^144^; in which the BPTES molecule was removed from the structure) and the AlphaFold-predicted PDZD8 structure (https://alphafold.ebi.ac.uk/entry/Q8NEN9)145,146 were used. Data were then illustrated using the PyMOL (ver. 2.5, Schrodinger) software. The amino acid residues P137, L139, E140, L142, Y145, G150, Q161, E162, K176, E177, D180, Q187, V209, T212, Q213, R216, K218, D223, S226, H230, F439, G444, E445, R446, V447, P450, R534, H535, F536, K538, L540, R544, and E545, a total of 33, which comprise the interface of GLS1 (same between KGA and GAC) for PDZD8 were then mutated (all to alanine) to generate the GLS1-33A mutant.

### Determination of oxygen consumption rates

For measuring oxygen consumption rates (OCR) in MEFs, cells were plated at 10,000 cells per well on a 96-well Seahorse XF Cell Culture Microplate (Agilent) in full medium (DMEM containing 10% FBS) overnight before experiment, followed by glucose starvation for desired time periods. For cells treated with inhibitors of glutaminolysis and FAO, Etomoxir at 20 μM for 10 h and BPTES at 10 μM for 8 h were used. Medium was then changed to Seahorse XF Base Medium supplemented with 10% FBS, 25 mM glucose (not included under starvation condition, and same hereafter), 4 mM glutamine (GlutaMAX) and 1 mM sodium pyruvate 1 h before the experiment. Cells were then placed in a CO_2_-free, XF96 Extracellular Flux Analyzer Prep Station (Agilent) at 37 °C for 1 h. OCR was then measured at 37 °C in an XF96 Extracellular Flux Analyzer (Agilent), with a Seahorse XFe96 sensor cartridge (Agilent) pre-equilibrated in Seahorse XF Calibrant solution in a CO_2_-free incubator at 37 °C overnight. The assay was performed on a Seahorse XFe96 Analyzer (Agilent) at 37 °C following the manufacturer’s instruction. Concentrations of respiratory chain inhibitors used during the assay were: oligomycin at 10 μM, FCCP at 10 μM, antimycin A at 1 μM and rotenone at 1 μM (all final concentrations). Data were collected using Wave 2.6.1 Desktop software (Agilent) and exported to Prism 9 (GraphPad) for further analysis according to manufacturer’s instructions.

The OCR of nematodes was measured as described previously^147^. Briefly, nematodes were washed with M9 buffer for 3 times. Some 15 to 25 nematodes were then suspended in 200 μl of M9 buffer, and were added to a well on a 96-well Seahorse XF Cell Culture Microplate. The measurement was performed as for MEFs, except that 10 μM FCCP and sodium azide (40 mM) were added to nematodes during the assay, and the temperature of Seahorse XFe96 Analyzer was set at 20 °C. Data were collected and analysed as in MEFs. At the end of the assay, the exact number of nematode in each well was determined on a Cell Imaging Multi-Mode Reader (Cytation 1, BioTek) and was used for normalising/correcting OCR results.

The OCR of intact muscle tissue was measured as described previously^98,148^, with modifications. In brief, mice were starved for desired durations, and were sacrificed through cervical dislocation. The gastrocnemius muscles from two hindlegs were then excised, followed by incubating in 4 ml of dissociation media (DM; by dissolving 50 μg/ml gentamycin, 2% (v/v) FBS, 4 mg/ml collagenase A in DMEM (21063-029)) in a 35-mm culture dish in a humidified chamber at 37 °C, 5% CO_2_, for 1.5 h. The digested muscle masses were then washed with 4 ml of pre-warmed collagenase A-free DM, incubated in 0.5 ml of pre-warmed collagenase A-free DM, and dispersed by passing through a 20 G needle for 6 times. Some 20 μl of muscle homogenates was transferred to a well of a Seahorse XF24 Islet Capture Microplate (Agilent). After placing an islet capture screen by a Seahorse Capture Screen Insert Tool (Agilent) into the well, 480 μl of pre-warmed aCSF medium (120 mM NaCl, 3.5 mM KCl, 1.3 mM CaCl_2_, 0.4 mM KH_2_PO_4_, 1 mM MgCl_2_, 5 mM HEPES, 15 mM glucose, 1× MEM non-essential amino acids, 1 mM sodium pyruvate, and 1 mM GlutaMAX; adjust to pH 7.4 before use) was added, followed by equilibrating in a CO_2_-free incubator at 37 °C for 1 h. OCR was then measured at 37 °C in an XFe24 Extracellular Flux Analyzer (Agilent), with a Seahorse XFe24 sensor cartridge (Agilent) pre-equilibrated in Seahorse XF Calibrant solution (Seahorse Bioscience, Agilent) in a CO_2_-free incubator at 37 °C overnight. Respiratory chain inhibitor used during the assay was oligomycin at 10 μM of final concentration. Data were collected using Wave 2.6.3 Desktop software (Agilent) and exported to Prism 9 (GraphPad) for further analysis according to the manufacturer’s instructions.

### Determination of electron transport chain integrity

The integrity of electron transport chain in muscles was determined through measuring the OCR of the permeabilised myocytes supplied with excessive substrate of each mitochondrial respiration complex on an Oxygraph-2k (Oroboros Instruments)^149^. In brief, the gastrocnemius muscle was dissected into thin fibre bundles and then immersed in ice-cold Isolation Solution A (10 mM Ca-EGTA buffer (2.77 mM CaK_2_EGTA and 7.23 mM K_2_EGTA), pH 7.1, 20 mM imidazole, 20 mM taurine, 49 mM K-MES, 3 mM K_2_HPO_4_, 9.5 mM MgCl_2_, 5.7 mM ATP, 15 mM phosphocreatine, and 1 mM leupeptin) and was then permeabilised by addition of 50 μg/ml saponin by gently mixing at 4 °C for 10 min. The permeabilised tissues were then washed three times by Respiration Medium B (0.5 mM EGTA, 3 mM MgCl_2_, pH 7.1, 20 mM taurine, 10 mM KH_2_PO_4_, 20 mM HEPES, 1 g/l BSA, 60 mM K-lactobionate, 110 mM mannitol and 0.3 mM dithiothreitol) before the assay. Some 5 mg of tissue suspended in Respiration Medium B was transferred to an oxygraphy chamber in an Oxygraph-2k (Oroboros Instruments), followed by incubation for 5 min. Glutamate (final 10 mM) and malate (5 mM) were added to the chamber to determine the resting complex I-supported respiration (without ADP addition), followed by addition of 5 mM ADP to determine the maximal complex I-supported respiration. Succinate (10 mM) was then added to the chamber to induce the complex II-supported respiration. Data were collected using DatLab software (v.7.3.0.3, Oroboros Instruments) and exported to Prism 9 for further analysis.

The integrity of electron transport chain in MEFs was determined on an Oxygraph-2k according to a previous study^150^. Briefly, MEFs were glucose-starved, and were then harvested by trypsinising, followed by centrifuged at 1,000 rpm for 3 min at room temperature. The pellets were re-suspended in the mitochondrial respiration medium MiR05 buffer (110 mM sucrose, 60 mM K-lactobionate, 0.5 mM EGTA, 3 mM MgCl_2_, 20 mM taurine, 10 mM KH_2_PO_4_, 20 mM HEPES, pH 7.1 and 0.1% BSA) pre-warmed at 30 °C to a density of 0.5 × 10^6^ cells/ml, followed by transferring 2 ml of cell suspension to a chamber of Oxygraph-2k. After stabilising for 10 min, the basal OCRs for intact cells were recorded. The baseline OCRs in permeabilised cells (also known as leak respiration, given that it is driven by the proton leak) were then determined after addition of 1 μl of digitonin (10 mg/ml stock solution in DMSO) to the chamber to expose the electron transport chain. The resting complex I-supported respiration was then recorded after adding 5 mM pyruvate, 10 mM glutamate and 2 mM malate to the chamber, followed by determining the maximal complex I-supported respiration through addition of 2.5 mM ADP. Succinate (10 mM) was then added to the chamber to induce the complex II-supported respiration. The maximal respiratory capacity was then determined by stepwise addition of FCCP to a final concentration of 0.5 μM. Data were collected using DatLab software (v.7.3.0.3, Oroboros Instruments) and exported to Prism 9 for further analysis.

### Confocal microscopy

The filamentation of GLS1 under glucose or glutamine starvation was determined as described previously^151^. Briefly, MEFs grown to 80% confluence on coverslips in 6-well dishes were fixed for 20 min with 4% (v/v) formaldehyde in PBS, 2 ml per well/coverslip at room temperature. The coverslips were rinsed twice with 2 ml of PBS and permeabilised with 2 ml of 0.1% (v/v) Triton X-100 in PBS for 10 min at 4 °C. After rinsing twice with 2 ml of PBS, the coverslips were incubated with rabbit anti-GLS1 antibody (1:100, diluted in Block Buffer (10% (v/v) NGS in PBS, with 0.1% (w/v) saponin) overnight (by placing 50 μl of antibody solution on a piece of Parafilm M to form a drop, followed by mounting a coverslip on the drop) in a humidified chamber at 4 °C. The cells were then rinsed three times with 2 ml of PBS, and then incubated with secondary antibody (Alexa Fluor 488 donkey anti-rabbit IgG; performed as in primary antibody incubation) for 8 h at room temperature in a humidified chamber in the dark. The coverslips were washed for another 4 times with 2 ml of PBS, and then mounted on slides using ProLong Diamond Antifade Mountant. Confocal microscopic images were taken using an LSM 980 (Zeiss) with a 63× 1.4 NA oil objective, during which a diode laser module (Lasos) at 488-nm was used to excite Alexa Fluor 488 dye. All parameters were kept unchanged for each picture taken. Images were processed and analysed on Zen Blue 3.3 software (Zeiss), and formatted on Photoshop 2023 software (Adobe).

SPLICS staining was performed as for staining the GLS1 filamentation, except that MEFs stably expressing short-range SPLICS were used. Quantification analysis was performed using the FIJI (ver. 1.53q, National Institutes of Health) software according to a previously study^93^, with modifications. In brief, a series of rectangular, identical region of interest (ROI) was created and equally allocated to each image. The SPLICS signal in each ROI was extracted using the “crop” command (by selecting Image > Crop). The Gaussian blur filter was then applied to each cropped image by selecting Process > Filters > Gaussian Blur, during which the “Sigma (Radius)” value was set at 1.5, followed by selecting “yes” on the “apply to all stacks” dialog. The background of each images was then subtracted by selecting Process > Subtract Background, during which the “Rolling ball radius” was set at 50 pixels. The SPLICS count was then calculated by selecting Process > Find Maxima, during which “Strict” and “Exclude edge maxima option” boxes were ticked, and “Prominence” was set at “> 10”.

The PLA/Duolink assay was performed using the Duolink In Situ Red Starter Kit (Mouse/Rabbit) according to the manufacturer’s instruction, with minor changes. In brief, MEFs expressing HA-tagged PDZD8 (or its mutants) were grown to 80% confluence on coverslips in 6-well dishes, followed by fixation for 20 min with 4% (v/v) formaldehyde in PBS, 2 ml per coverslip/well at room temperature. The coverslips were rinsed twice with 2 ml of PBS and permeabilised with 2 ml of 0.1% (v/v) Triton X-100 in PBS for 10 min at 4 °C. Cells were then blocked with Duolink Blocking Solution (50 μl per coverslip) in a humidified chamber at 37 °C for 1 h. Cells were then incubated with primary antibodies (mouse anti-HA-tag and rabbit anti-GLS1; 1:100 diluted with Duolink Antibody Diluent; 50 μl per coverslip) in a humidified chamber at 4 °C for 12 h, followed by washing with two changes of 2 ml of Wash Buffer A, 5 min per change, at room temperature. The coverslip was then incubated with PLUS and MINUS PLA probe solution (freshly prepared by mixing 10 μl of PLA probe MINUS stock, 10 μl of PLA probe PLUS stock with 30 μl of Duolink Antibody Diluent; 50 μl per coverslip) in a humidified chamber at 37 °C for 1 h, followed by washing with two changes of 2 ml of Wash Buffer A, 5 min per change, at room temperature. The coverslip was then incubated with Ligation Solution (freshly prepared by 1:5 diluting Duolink Ligation buffer with water, followed by addition of Ligase stock at a ratio of 1:50; 50 μl per coverslip) in a humidified chamber at 37 °C for 0.5 h, followed by washing with two changes of 2 ml of Wash Buffer A, 5 min per change, at room temperature. The coverslip was then incubated with Amplification Solution (freshly prepared by 1:5 diluting Amplification buffer with water, followed by addition of Polymerase stock at a ratio of 1:80; 50 μl per coverslip) in a humidified chamber at 37 °C for 100 min, followed by washing with two changes of 2 ml of Wash Buffer B, 10 min per change, at room temperature. The coverslip was then washed with 2 ml of 0.01× Wash Buffer B for 1 min at room temperature, followed by mounting with 15 μl of Duolink PLA Mounting Medium with DAPI for 30 min, and then subjected to imaging using an LSM 980 (Zeiss) as described above, except that a DPSS laser module (Lasos) at 594-nm and a diode laser module (Lasos) at 405-nm were used to excite the PLA and DAPI, respectively.

### Determination of mitochondrial ROS

For detecting the mitochondrial ROS levels in MEFs, cells were grown in 35-mm glass-bottom dishes (cat. D35-20-10-N, In Vitro Scientific) to 50% confluence. Cells were treated with 5 μM (final concentration) MitoSOX dye for 0.5 h at 37 °C, then washed three times with 2 ml of pre-warmed culture medium, and incubated in fresh medium containing ProLong™ Live Antifade Reagent before imaging. During imaging, live cells were kept at 37 °C, 5% CO_2_ in a humidified incubation chamber (Incubator PM S1, Zeiss). Images were taken using an LSM 980 (Zeiss) with a 63× 1.4 NA oil objective, during which a DPSS laser module (Lasos) at 594-nm was used to excite mitoSOX. The parameters, including ‘PMT voltage’, ‘Offset’, ‘Pinhole’ and ‘Gain’, were kept unchanged between each picture taken. The resolution of image is 1,024×1,024 pixels. Images were processed and analysed on Zen Blue 3.3 software (Zeiss), and formatted on Photoshop 2023 software (Adobe).

For detecting the mitochondrial ROS levels in nematodes, synchronised nematodes cultured to L4 stage were treated with 2-DG or subjected to CR for 48 h. Nematodes were then treated with 5 μM (final concentration; added into the NGM plate containing the OP50 bacteria) MitoSOX dye for another 12 h at 20 °C, followed by placing on the centre of an injection pad (prepared by placing 2 drops (approximately 50 μl) of boiling 4% agarose (w/v) onto the centre of a glass coverslip (24 × 50 mm, 0.13–0.15 mm thickness), immediately followed by flattening with another coverslip, then dried at room temperature for 24 h). The pad was then subjected to imaging as described in those of MEFs, except that an LSM 900 (Zeiss) with a ×20, 0.8 NA plan-Apochromat air objective was used, during which a laser module URGB (cat. 400102-9301-000, Toptica) using a 10-mW laser line at 561 nm was applied. Images were processed by Zen 3.1 software (Zeiss), and formatted on Photoshop 2023 software (Adobe).

For detecting mitochondrial ROS levels in muscle tissues, mice were starved for desired time periods, and were sacrificed by cervical dislocation. The gastrocnemius muscle was then quickly excised and sliced to 0.05 cm^3^ cubes, followed by immediately soaking in O.C.T. Compound at −20 °C for 20 min. The embedded tissues were then sectioned into 15-μm slices using a CM1950 Cryostat (Leica). Sections were stained with 40 ml of 5 μM (final concentration; by diluting the DMSO stock solution with PBS) MitoSOX dye for 30 min at 37 °C in a Coplin jar, followed by washing for 3 times, 5 min each with 40 ml of PBS at room temperature. Sections were then mounted with Antifade Mounting Medium, and were imaged on a DM4 B (Leica) microscope.

### TEM and FIB-SEM

The 2D-TEM imaging was performed based on the in situ embedding and sectioning method as described previously^152^, with minor modifications. In brief, MEFs were grown in a 3.5-cm dish containing 1 ml of DMEM to approximately 70% confluence. Cells were fixed by gently adding 0.2 ml of 25% (v/v) glutaraldehyde solution into the DMEM, followed by incubating at room temperature for 15 min. Cells were incubated with 1 ml of 2.5% (v/v) glutaraldehyde solution (freshly prepared by diluting 25% (v/v) glutaraldehyde in 0.1 M Phosphate Buffer (by mixing 0.2 M Na_2_HPO_4_ with 0.2 M NaH_2_PO_4_ (both dissolved in water, and adjusted pH to 7.4) at a ratio of 81:19, and then diluted with equal volume of water) at room temperature for 2 h. Cells were then washed with 1 ml of ice-cold 0.1 M Phosphate Buffer for 3 times, 15 min each, followed by the addition of 1 ml of ice-cold 20 mM glycine solution (in 0.1 M Phosphate Buffer) for 5 min on ice, and were then washed with 1 ml of ice-cold 0.1 M Phosphate Buffer for 3 times, 15 min each. Cells were then stained with 1% (w/v) OsO_4_ solution (in 0.1 M Phosphate Buffer, supplemented with 1.5% K_3_Fe(CN)_6_) on ice for 1 h, followed by washing for 5 times, 3 min each with ice-cold water. Cells were then incubated with 1% (w/v) thiocarbonohydrazide (freshly prepared by dissolving thiocarbonohydrazide in water at 60 °C for 1 h with regular shaking, followed by cooling down to room temperature) at 25 °C for 20 min in the dark, followed by washing for 5 times, 3 min each with water at room temperature. Cells were then incubated with 1% (w/v) OsO_4_ solution (dissolved in water) for 40 min at 4 °C, followed by washing for 5 times, 3 min each with water at 4 °C. Cells were stained in ice-cold 2% (w/v) uranyl acetate solution for 12 h at 4 °C in the dark, and were then washed for 5 times, 3 min each, with ice-cold water. Dehydration was then performed by sequentially incubating cells in the following ice-cold solutions: 30, 50, and 70% (v/v) ethanol (in water), each for 7 min at 4 °C, followed by incubating in 90, 100 and 100% v/v) ethanol (in water), each for 7 min at room temperature. Cells were then quickly submerged in ethanol/Spon 812 resin (3:1; the Spon 812 resin was prepared by mixing with 10 ml of SPI-Pon 812, 4.5 ml of DDSA, 6 ml of NMA with 0.6 ml of DMP-30; all supplied in the SPI-Pon 812 Embedding Kit) mixture at room temperature for a 1-h incubation, and then in ethanol/resin (1:1) mixture at room temperature for 2 h, followed by ethanol/resin (1:3) at room temperature for 2 h, and finally 100% resin at room temperature for two rounds: first round overnight, and next round for 6 h. Resin was then completely drained, and the cells were spread with a thin layer of fresh resin (with a total volume of approximately 400 μl, and below the thickness of 1 mm), followed by baking at 60 °C in a hot-wind drying oven for 48 h. The embedded cells were then sectioned into 70-nm slices on an EM UC7 Ultramicrotome (Leica) after cooling down to room temperature. Sections were then stained with lead citrate solution (by dissolving 1.33 g of Pb(NO_3_)_2_ and 1.76 g of sodium citrate in 42 ml of di-distilled water, followed by addition of 8 ml of 1 M NaOH) for 5 min at room temperature before imaging using an AMT-XR81DIR camera on an electron microscope (HT-7800, Hitachi) using TEM system control software (Ver. 01.20, Hitachi). The mitochondria and ER on each image were then segmented, and the measurements such as the length of the ER-mitochondria contact were acquired using the FIJI software. In particular, the “aspect ratio”, a parameter/factor invented to describe the morphology of a mitochondrion (defined as the length of the major axis divided by the length of the minor axis)^153,154^ was calculated using the “Analyze Particles” plugin of FIJI software as described in ref.^155^.

The 3D-FIB-SEM imaging was performed as described previously^156^. Briefly, MEFs were cultured on 35-mm dishes and fixed in 2.5% glutaraldehyde (diluted in 0.1 M Phosphate Buffer) for 2 h at room temperature. The cells were then washed with 0.1 M Phosphate Buffer three times, followed by incubation in 1% (w/v) OsO_4_ solution (in distilled water, supplemented with 1.5% K_3_Fe(CN)_6_) for 1 h at 4 °C, and then dehydration with the following ethanol solutions (50%, 70%, 80%, 90%, 100%, and 100% v/v; in water), each for 2 min on ice. The cells were then quickly rinsed with 0.1 M Phosphate Buffer at room temperature, and then incubated in ethanol/Spon 812 resin (1:1; the Spon 812 resin was prepared as in 2D-SEM) mixture at room temperature for a 30-min incubation, and then in ethanol/resin (1:2) mixture at room temperature for 30 min, followed by ethanol/resin (1:3) at room temperature for 30 min, and finally 100% resin at room temperature for two rounds, each for 1 h. Resin was then polymerised in an oven at 60 °C for 48 h. After sectioning and staining (with uranyl acetate and lead citrate) as in the 2D-TEM, sections were subjected for FIB milling and scanning electron microscopy imaging. During imaging, a layer of block surface was milled by gallium ion beam, and the block surface was imaged using an electron beam with a 2-kV acceleration voltage, 0.4-nA current, and 8-µs dwell time on a Helios NanoLab G3 UC FIB-SEM (FEI). After image collection, the images were imported into the Amira software (ver. 2022.2, Thermo Scientific), followed by aligning using the DualBeam 3D Wizard module, and then exported as an “image-stack”, TIF format. The image stacks were then subjected to structural segmentation. The segmented mitochondria, ER, along with the ER-mitochondria contact, were stitched and reconstructed to yield the 3D-FIB images. The measurements, including the volumes of mitochondria, ER, and the ER-mitochondria contact, were acquired through the Amira software.

### RET-FLIM assay

FRET-FLIM experiments were carried out as described previously^108^, with minor modifications. Briefly, MEFs stably expressing GFP (“donor only”, as a control), RFP-PDZD8-GFP, or different combinations of PDZD8-GFP and GLS1-mCherry were cultured in 35-mm glass-bottom dishes (cat. D35-20-10-N, In Vitro Scientific) to 60-80% confluence. Cells were starved for glucose or not, followed by determining the fluorescence lifetime of GFP in different cells cultured in a humidified chamber with 5% CO_2_ at 37 °C using a STELLARIS 8 FALCON (Leica) systems equipped with HyD X and HyD SMD detectors and an HC PL APO CS2 63x/1.40 OIL objective (Leica). Cells were excited with a 460-nm laser via the systems’ tuneable White Light Laser (WLL), and photon arrival times were recorded with an HyD X detector covering the GFP emission spectrum (460-510 nm). All parameters were kept unchanged between imagings. Images were taken and analysed by LAS X Software (Leica). In all experiments, the position of the focal plane was actively stabilised using the Leica Auto Focus Control (AFC) to prevent any focal drift or focus artefacts.

### SIM and STORM imaging

MEFs grown to 50% confluence in a 35-mm dish (for SIM; cat. D35-20-10-N, In Vitro Scientific), and in Lab-Tek II chambered no. 1.5 German coverglass system (for STORM; cat. 155409, 8 Chamber, Nunc) were treated following the Semi-intact IF protocol as described previously^42,157^, with minor modifications. Briefly, cells were rinsed with PBS once, and treated with Buffer I (25 mM HEPES, pH 7.2, 125 mM potassium acetate, 5 mM magnesium acetate, 1 mM DTT, 1 mg/l glucose and 25 μg/ml digitonin) for 1 min on ice, and then Buffer II (25 mM HEPES, pH 7.2, 125 mM potassium acetate, 5 mM magnesium acetate, 1 mM DTT and 1 mg/l glucose) for another 10 min on ice. The cells were then fixed with ice-cold methanol in PBS on ice for 10 min. The slides were rinsed twice with PBS and cells were then permeabilised with 0.05% Triton X-100 in PBS for 5 min at 4 °C. After rinsing twice with PBS, the slides were blocked in Block Buffer for 30 min. The slides were washed twice with PBS and incubated with primary antibodies diluted in Block Buffer overnight at 4 °C. The cells were then rinsed three times with PBS, and then incubated with secondary antibodies for another 8 h at 4 °C in the dark, followed by washing for four times with PBS before imaging.

SIM images were acquired using a Multi-SIM (multimodality structured illumination microscopy) imaging system (NanoInsights-Tech Co., Ltd.) equipped with a 100× 1.49NA oil objective (CFI SR HP Apo, Nikon), a solid-state, single-mode laser (containing the 488-nm, 561-nm and 640-nm laser beams) and an sCOMS (complementary metal-oxide-semiconductor) camera (ORCA-Fusion C15440-20UP, HAMAMATSU). The immersion oil with a refractive index of 1.518 was chosen for this experiment, and the microscope was calibrated with 100-nm fluorescent spheres before the experiment. The SIM images were taken through the low NA GI-SIM mode with a 50-mw laser power and a 20-ms exposure time via SI-Recon 2.11.19 software (NanoInsights-Tech). Images were then reconstructed using the SI-Recon 2.11.19 software, during which the parameters were set as: a) pixel size: 30.6 nm; b) optical transfer functions: channel-specific; c) Wiener filter: constant 0.01, for the TIRF-SIM mode; and d) negative intensities background: discard. After reconstruction, images were denoised under the total variation (TV) constraint mode. The denoised images were then formatted using Photoshop 2023 software (Adobe).

STORM imaging was performed as described previously^42,158^, with minor modifications. Briefly, the STORM imaging buffer supplemented with MEA was freshly prepared before the experiment by mixing 7 μl of GLOX (14 mg of glucose oxidase, 50 μl of catalase (17 mg/ml), 200 μl of buffer A (10 mM Tris, pH 8.0 and 50 mM NaCl), vortexed to dissolve and cooled on ice) with 70 μl of 1 M MEA (77 mg of MEA dissolved in 1.0 ml of 0.25 M HCl), followed by adding to 620 μl of buffer B (50 mM Tris, pH 8.0, 10 mM NaCl and 10% (m/v) glucose) in a 1.5-ml Eppendorf tube, and followed by brief vortex. The mixture was then added to each well, and images were taken on an N-STORM (Nikon). The imaging was performed using an inverted microscope system (Ti-E Perfect Focus; Nikon) equipped with a monolithic laser combiner (MLC400, Agilent) containing solid-state lasers of wavelengths 405 nm (at 100 mW of maximum fibre output power), 488 nm (200 mW) and 561 nm (150 mW) and a 647-nm laser at 300 mW. After locating a suitable field, a diffraction-limited TIRF image was acquired for reference, followed by a STORM acquisition. The 647-nm laser was then sequentially fired at 100% power to excite all possible fluorophore molecules and photoswitch them into a non-emitting dark state, and then the 561-nm laser. The emitted wavelengths from Alexa Fluor 647 and CF 568 fluorophores were then sequentially collected by the plan-Apochromat 100× 1.49NA TIRF objective (Nikon), filtered by an emission filter set (FF01-586/20-25×3.5 and FF01-692/40-25; Semrock), and detected on an electron-multiplying charge-coupled device camera (iXon DU-897, Andor Technology). During imaging, 20,000 sequential frames of each channel were acquired. The image acquisition, lateral drift correction and data processing were performed using NIS Elements software with STORM package (v.4.30 build 1053, Nikon) as previously described^159,160^.

### Subcellular fractionation

Mitochondria and mitochondria-associated membranes (MAMs) were purified as described previously^161^, with minor modifications^53^. Briefly, 40 10-cm dishes of MEFs (60-80% confluence) were collected by scrapping at room temperature, followed by centrifugation for 5 min at 500*g* at 37 °C. Cells were then resuspended in 20 ml of ice-cold IB_cells_-1 buffer (225 mM mannitol, 75 mM sucrose, 0.1 mM EGTA and 30 mM Tris-HCl, pH 7.4), and dounced for 100 strokes in a 40-ml Dounce homogeniser (using the small clearance pestle, or the pestle B; cat. D9188, Sigma), followed by two times of centrifugation for 5 min at 600*g* at 4 °C. The supernatants were then collected and centrifuged for 10 min at 7,000*g* at 4 °C. The pellets were then washed twice with 20 ml of ice-cold IB_cells_-2 buffer (225 mM mannitol, 75 mM sucrose and 30 mM Tris-HCl pH 7.4). The suspensions were centrifuged at 7,000*g*, and again at 10,000*g*, both for 10 min at 4 °C. The pellets were then resuspended in 2 ml of ice-cold MRB buffer (250 mM mannitol, 5 mM HEPES pH 7.4 and 0.5 mM EGTA), and were loaded on top of 10 ml of Percoll medium (225 mM mannitol, 25 mM HEPES pH 7.4, 1 mM EGTA and 30% Percoll (v/v)) in 14 × 89-mm centrifuge tubes (cat. 344059, Beckman). The tubes were then centrifuged on a SW 41 Ti rotor (Beckman) at 95,000*g* for 0.5 h at 4 °C. After centrifugation, the dense band located near the bottom of each tube was collected as mitochondrial fraction, while the band located at the interface between MRB and Percoll cushions as MAMs. The mitochondrial fractions were diluted with 10 volumes of MRB buffer, followed by centrifugation at 6,300*g* for 10 min at 4 °C; the pellets were resuspended and washed with 2 ml of MRB buffer, followed with centrifugation at 6,300*g* for 10 min at 4 °C to obtain pure mitochondria (the pellets). The MAM fractions were centrifuged at 6,300*g* for 10 min at 4 °C, and the supernatants were combined and transferred to 25 × 83-mm centrifuge tubes (cat. 344367, Beckman), followed by centrifuge at 95,000*g* on a SW 32 Ti rotor (Beckman) for 1 h at 4 °C to obtain the pure MAM (the pellets).

Outer mitochondrial membrane (OMM) was purified by suspending pure mitochondria by 100 μl of MRB buffer containing 0.5 (w/v) digitonin, followed by centrifuge at 10,000*g* for 15 min at 4 °C. The supernatant contains OMM, and the pellet mitoplast. The mitoplasts were suspended with 100 μl of MRB buffer containing 1% (v/v) Triton X-100, followed by centrifuge in an 8 × 34-mm centrifuge tube (cat. 45235-AV, Thermo Scientific) at a S120-AT3 rotor (Thermo Scientific) at 100,000*g* for 30 min at 4 °C, and the pellets and supernatants contain inner mitochondrial membrane (IMM) and matrix, respectively.

ER was purified according to the protocol optimised by combining the traditional microsome-based density gradient isolation method (Endoplasmic Reticulum Isolation Kit developed by Sigma) with the cell surface biotinylation reaction method (developed and optimised by Pierce), and was described previously^42^. Briefly, MEFs from 40 10-cm dishes (80% confluence) were quickly washed with ice-cold PBS (10 ml each dish) twice, followed by incubating with 250 μg/ml of sulfo-NHS-SS-biotin (freshly dissolved in ice-cold PBS, 10 ml each dish) for 30 min with gentle agitation on an orbital shaker at 4 °C. Some 500 μl of 1 M Tris (pH 8.0 at 4 °C) was then added to each dish to quench the biotinylation reaction. Cells were collected afterwards by scrapping, followed by centrifugation at 600*g* for 5 min, and then washed with 40 ml of ice-cold PBS twice. Cells were then re-suspended in 10 ml of 1× Hypotonic Extraction Buffer and then incubated at 4 °C for 30 min, with gentle mixing in the middle. Cells were then centrifuged at 600*g* at 4 °C for 5 min, and the pellet was re-suspended with 6 ml of 1× Isotonic Extraction Buffer, followed by mixing in a 7-ml Dounce homogeniser (using the small clearance pestle; cat. D9063, Sigma) for 10 strokes. The homogenates were centrifuged at 1,000*g* for 10 min at 4 °C, and the supernatants (PNS) were further centrifuged at 12,000*g* for 15 min at 4 °C, yielding the supernatants as the post-mitochondrial fraction (PMF). The PMF was loaded in two 11 × 60 mm centrifuge tubes (cat. 344062, Beckman) and then centrifuged on an SW 60 Ti rotor (Beckman) at 100,000*g* for 1 h at 4 °C. The pellet was re-suspended with 0.5 ml of 1× Isotonic Extraction Buffer, and was mixed in a 2-ml Dounce homogeniser (using the small clearance pestle; cat. D8938, Sigma) for 20 strokes, yielding the microsomal suspension. The suspension was mixed with 0.25 ml of OptiPrep, and was carefully layered on the top of 1 ml of 30% OptiPrep solution (by mixing 0.5 ml of OptiPrep with 0.5 ml of 1× Isotonic Extraction Buffer) in an 11 × 60 mm centrifuge tube. Some 2 ml of 15% OptiPrep solution (by mixing 0.5 ml of OptiPrep with 1.5 ml of 1× Isotonic Extraction Buffer) was then carefully layered on the top of the sample. The tube was then centrifuged on an SW60 Ti rotor at 150,000*g* for 3 h at 4 °C. The top 0.6 ml of 15% OptiPrep solution was discarded, and the remaining 200 μl of fraction was collected as the crude ER fraction. The fraction was then incubated with 100 μl of NeutrAvidin Agarose (pre-balanced by 1× Isotonic Extraction Buffer) for another 2 h. The supernatant contains ER fraction.

### Identification of AMPK substrates in MAM

To identify the substrate(s) of AMPK in the ER-mitochondria contact, the mitochondria and MAM fractions purified from 120 10-cm dishes of glucose-starved MEFs were dissolved with 5 ml of ice-cold Triton lysis buffer, followed by sonication and centrifugation at 4 °C, 20,000*g* for 15 min. Cell lysates were incubated with anti-pan-phospho-AMPK-substrates antibodies overnight. Protein aggregates were pre-cleared by centrifugation at 20,000*g* for 10 min, and protein A/G beads (1:250, pre-balanced with Triton lysis buffer) were then added into the lysate-antibody mixture, and incubated for another 3 h at 4 °C with rotating at 60 r.p.m. The beads were centrifuged and washed with 100 times volume of Triton lysis buffer for 3 times (by centrifuging at 2,000*g*) at 4 °C and then mixed with an equal volume of 2× SDS sample buffer (without bromophenol blue addition), and boiled for 10 min before subjecting to SDS-PAGE. After staining with Coomassie Brilliant Blue R-250 dye, gels were decoloured and the excised gel segments were subjected to in-gel chymotrypsin digestion, and then dried. Samples were analysed on a nanoElute (Bruker) coupled to a timsTOF Pro (Bruker) equipped with a CaptiveSpray source. Peptides were dissolved in 10 μl of 0.1% formic acid (v/v) and were loaded onto a homemade C18 column (35 cm × 75 μm, ID of 1.9 μm, 100 Å). Samples were then eluted with linear gradients of 3–35% acetonitrile (v/v, in 0.1% formic acid) at a flow rate of 0.3 μl min^−1^, for 60 min. MS data were acquired with a timsTOF Pro mass spectrometer (Bruker) operated in PASEF mode, and were analysed using Peaks Studio software (X^+^, Bioinformatics Solutions). The mouse UniProt Reference Proteome database was used for data analysis, during which the parameters were set as: a) precursor and fragment mass tolerances: 20 ppm and 0.05 Da; b) semi-specific digest mode: allowed; c) maximal missed cleavages per peptide: 3; d) variable modifications: oxidation of methionine, acetylation of protein N-termini, and phosphorylation of serine, threonine and tyrosine; e) fixed modification: carbamidomethylation of cysteine.

### Prokaryotic protein expression

Expression plasmid for the His-tagged, heterotrimeric AMPK was kindly provided by Dr. Dietbert Neumann (constructed in ref. ^162^), and the plasmid for rat kinase domain (KD) of CaMKK2 (aa 129-503) by Dr. Anthony Means (constructed in ref. ^163^). The cDNAs encoding human AMPKα1-KD (aa 27-290) (ref. ^164^) and rat CaMKK2-KD^42^ were inserted into the pET-28a (Novagen) vectors for expressing His-tagged recombinant proteins as described previously. The expression plasmids for the two isozymes of GLS1, PDZD8 and mutants were constructed by inserting respective cDNAs into pGEX-4T-1 (Cytiva) vectors for expressing GST-tagged recombinant proteins. KGA and GAC cDNAs were also cloned into the pET-28a vectors for bacterial expression. The pET-28a and pGEX-4T-1 plasmids were transformed into the *E*. *coli* strain BL21 (DE3) (cat. EC0114, Thermo Scientific), followed by culturing in LB medium in a shaker at 200 r.p.m. at 37 °C. The cultures of transformed cells were induced with 0.1 mM IPTG at an OD_600_ of 1.0. After incubating for another 12 h at 160 r.p.m. at 16 °C, the cells were collected. For His-tagged proteins, cells were homogenised in a His binding buffer (50 mM sodium phosphate, pH 7.4, 150 mM NaCl, 1% Triton X-100, 5% glycerol, and 10 mM imidazole), and for GST-tagged proteins, with a GST binding buffer (PBS supplemented with 10 mM β-mercaptoethanol and 1% Triton X-100) on ice. The homogenates were then sonicated on ice, and were subjected to centrifugation at 150,000*g* for 30 min at 4 °C, followed by purification of His-tagged proteins with Nickel Affinity Gel (pre-balanced with His binding buffer), or GST-tagged proteins with Glutathione Sepharose 4 Fast Flow Gel (pre-balanced with GST binding buffer) at 4 °C. The Nickel Affinity Gel was then washed with 100 times the volume of ice-cold His wash buffer (50 mM sodium phosphate, pH 7.4, 150 mM NaCl, and 20 mM imidazole), and the Glutathione Sepharose gel with 100 times the volume of ice-cold PBS. His-tagged proteins were eluted from the resin by His elution buffer (50 mM sodium phosphate, pH 7.4, 150 mM NaCl, and 250 mM imidazole), and GST-tagged proteins by GST elution buffer (50 mM Tris–HCl, pH 8.0, and 10 mM reduced glutathione) at 4 °C. In particular, to avoid the degradation of the full-length PDZD8 protein, a relatively large volume of GST binding buffer (i.e., a diluted bacteria homogenate, e.g., 300 ml of GST binding buffer for 3,600 ml of bacteria suspension at an OD_600_ of 1.0), a low sonication power (e.g., < 25% maximal power output on a VCX 750 (Sonics) sonicator equipped with a 1/4” (6-mm) stepped microtip (630-0435, Sonics)), and a shorter sonication duration (100 cycles of 3 s pulse with 3 s interval for 50 ml of bacteria homogenate) were applied. Proteins were concentrated to approximately 3 mg/ml by ultrafiltration (Millipore, UFC905096) at 4 °C, then subjected to gel filtration (Cytiva, Superdex 200) balanced with a buffer containing 50 mM Tris-HCl, pH 7.4 and 150 mM NaCl.

### Phosphorylation of PDZD8 by AMPK in vitro

For those experiments using heterotrimeric AMPK complex as the kinase in the assay system, AMPK complex was pre-activated by CaMKK2-KD as described previously^42,165,166^, with minor modifications. Briefly, GST-tagged CaMKK2-KD was incubated with Glutathione Sepharose 4 Fast Flow Gel (5 μg of protein/μl gel; pre-balanced with PBS) in PBS at 4°C for 1 h, followed by washing with 100 times the volume of ice-cold CaMKK2 kinase assay buffer (50 mM Tris, pH 8.0, 2 mM DTT, 100 mM NaCl, 10 mM MgCl_2_ and 1 mM ATP) twice. Some 1 μl of gel-immobilised CaMKK2 was then incubated with 5 μg of His-tagged AMPK complex in CaMKK2 kinase assay buffer supplemented with 5 mM ATP (total volume: 60 μl) on a thermomixer at 30 °C for 30 min. CaMKK2 was then removed by centrifugation at 2,000*g*, 4 °C for 30 s, yielding phosphorylated AMPK in the supernatant. For those experiments using His-tagged AMPK-KD as the kinase, proteins were directly subjected to the assay without being pre-phosphorylated by CaMKK2.

Phosphorylated PDZD8 proteins were prepared in an AMPK kinase assay system as described previously^164,167^. Briefly, 50 μg of GST-tagged PDZD8 was incubated with 10 μl of Glutathione Sepharose gel (pre-balanced with PBS) in PBS at 4 °C for 1 h, followed by washing with 100 times the volume of ice-cold PBS twice. The gel was then incubated with 5 μg of AMPK-KD or phosphorylated AMPK complex in 250 μl of AMPK kinase assay buffer (50 mM MOPS, pH 7.0, 100 mM NaCl, 0.1 mM EDTA, 10 mM MgCl_2_, and 5 mM ATP) at 25 °C for 2 h, followed by washing with 100 times the volume of ice-cold PBS twice, and PDZD8 proteins were eluted with GST elution buffer. The eluents were either subjected to immunoblotting or to assays for enzymatic activities of GLS1 (see below).

### Enzymatic activity

Activity of GLS1 in cell-free system was determined through a GLS1-GDH-coupled assay system as described previously^72^. Briefly, in each reaction, 10 nM GLS1 and 100 nM PDZD8 were pre-incubated in 100 μl of Reaction buffer (50 mM Tris-acetate, pH 8.6 and 0.2 mM EDTA) for 2 h on a rotator at 4 °C, followed by mixing with 3 U of GDH and 40 mM NAD^+^ in 10 μl of Reaction buffer at 25 °C for 30 min. The reaction was initiated by pipetting the GLS1-PDZD8 mixture into a well of a glass-bottom, 96-well microplate (cat. 3635, Corning) containing 90 μl of glutamine solution (at a desired concentration used in the assay; dissolved in Reaction buffer) pre-warmed at 25 °C, followed by mixing on a SpectraMax M5 microplate reader (Molecular Devices). For the assays performed in the presence of phosphate, K_2_HPO_4_ (2 M stock, pH 9.4) at a final concentration of 20 mM was added into the glutamine solution. The effects of PDZD8 on GLS1 activity were assessed with the initial velocities of NADH formation during the reaction through which the OD_340_, recorded at 30-s intervals on a SpectraMax M5 microplate reader using the SoftMax Pro software (v.5.4.1.1, Molecular Devices), was increased. All measurements were carried out in triplicate. The catalytic velocities were calculated by the extinction coefficient for NADH at 340 nm, which is 6,220 cm^−1^ M^−1^, and 0.625 cm for path length. Data were collected using the SoftMax Pro software and exported to OriginPro software (v.9.2.0, OriginLab) for further analysis.

Activities of GLS1 in cells were determined through a semi-permeabilised system. Some 24 h before the measurement, MEFs were seeded in 24-well dishes (#142485, Thermo Scientific), and were cultured to 90% confluence. Cells were then starved for glucose for desired periods of time, followed by gentle rinsing with 350 μl of PBS containing 0.01% (v/v) NP-40 (titrated according to the conditions described in ref. ^87^) and 1% (v/v) protease inhibitor cocktail for each well for 60 s at 25 °C. The permeabilised cells were then quickly washed with 300 μl of PBS twice to remove the detergents, and the reaction was initiated by pipetting 200 μl of Reaction Buffer (20 mM glutamine dissolved in PBS), or Reaction Buffer supplemented with 10 μM BPTES for determining baseline, into each well, followed by mixing on a thermomixer (Thermomixer R, Eppendorf) at 37 °C, 50 r.p.m. for 1 h. The glutamate yielded was then measured using the Glutamate Assay Kit according to the manufacturer’s instruction. In brief, 100 μl of Reaction Buffer in each well was collected, followed by centrifugation at 20,000*g* for 10 min. Some 20 μl of supernatant was collected, and then mixed with 30 μl of Glutamate Assay Buffer, followed by incubating with 100 μl of Reaction Mix (prepared by mixing 8 μl of Glutamate Developer and 2 μl of Glutamate Enzyme Mix in 90 μl of Glutamate Assay Buffer) at 37 °C for 30 min in the dark. The OD_450_ was then recorded by a SpectraMax M5 microplate reader using the SoftMax Pro software.

## Statistical analysis

Statistical analyses were performed using Prism 9 (GraphPad Software), except for the survival curves, which were analysed using SPSS 27.0 (IBM). Each group of data was subjected to Kolmogorov-Smirnov test, Anderson-Darling test, D’Agostino-Pearson omnibus test or Shapiro-Wilk test for normal distribution when applicable. An unpaired two-tailed Student’s *t*-test was used to determine significance between two groups of normally distributed data. Welch’s correction was used for groups with unequal variances. An unpaired two-tailed Mann-Whitney test was used to determine significance between data without a normal distribution. For comparisons between multiple groups, an ordinary one-way or two-way ANOVA was used, followed by Tukey, Sidak, Dunnett or Dunn as specified in the legends. The assumptions of homogeneity of error variances were tested using F-test (*P* > 0.05). For comparison between multiple groups with two fixed factors, an ordinary two-way ANOVA was used, followed by Tukey’s or Sidak’s multiple comparisons test as specified in the legends. Geisser-Greenhouse’s correction was used where applicable. The adjusted means and s.e.m., or s.d., were recorded when the analysis met the above standards. Differences were considered significant when *P* < 0.05, or *P* > 0.05 with large differences of observed effects (as suggested in refs. ^168,169^).

## Data availability

The data supporting the findings of this study are available within the paper and its Supplementary Information files. The raw RNA sequencing data corresponding to the expression of ROS-depleting enzymes in nematodes under 2-DG and CR treatments have been deposited in the Genome Sequence Archive^170^ in National Genomics Data Center^171^, China National Center for Bioinformation/Beijing Institute of Genomics, Chinese Academy of Sciences (GSA: CRA011002) that are publicly accessible at https://ngdc.cncb.ac.cn/gsa. The MS proteomics data have been deposited to the ProteomeXchange Consortium (http://proteomecentral.proteomexchange.org) through the iProX partner repository^172,173^ with the dataset identifier PXD041428. Materials, reagents or other experimental data are available upon request. Full immunoblots are provided as Supplementary Information Fig. 1. Source data are provided with this paper.

## Code availability

The analysis was performed using standard protocols with previously described computational tools. No custom code was used in this study.

## Acknowledgements

We thank Dr. S. Morrison (University of Texas Southwestern Medical Center) for providing the *AMPK*α*1*^F/F^ (The Jackson Laboratory, 014141), and *AMPK*α*2*^F/F^ mice (The Jackson Laboratory, 014142); Dietbert Neumann (Maastricht University) for the tricistronic AMPK expression plasmid; Anthony Means (Duke University) for the CaMKK2 expression plasmid; Changchun Xiao (Sanofi China) for the *Rosa26*-CTV vector; Zhengfan Jiang (Peking University) for the manganese adjuvant; Su-Qin Wu and Ying He (Xiamen University) for mouse in vitro fertilisation; Xin Chen (Xiamen University) for adjusting the parameters used for imaging PDZD8 and GLS1 by STORM; Shengrong Xu (Xiamen University) for titrating the conditions used for bacterially expressing and purifying full-length PDZD8; Dijin Xu (Yale University) for critical suggestions on screening the detergent in the lysis buffer to avoid the non-specific binding of PDZD8 during the IP assays; Jingdong Zhuang (Xiamen University) for optimising the protocols for fractioning MAM, OMM and IMM, Xiaoyu Niu (Xiamen University) for the artwork of Extended Data Fig. 10c, and all the other members of the S.-C.L. laboratory for technical assistance. We also acknowledge the *Caenorhabditis* Genetics Center and the National BioResource Project for supplying nematode strains; and the research staff from Guangzhou Computational Super-Resolution Biotech Co., Ltd., and Beijing NanoInsights-tech Co., Ltd. for technical assistance with super-resolution imaging experiments using HIS-SIM and Multi-SIM, respectively. The artworks shown in Extended Data Fig. 5l were modified from elements created by Servier Medical Art (https://smart.servier.com/) licenced under a Creative Commons Attribution 3.0 Unported Licence (https://creativecommons.org/licenses/by/3.0/). This work was supported by grants from the National Key R&D Program of China (2020YFA0803402), the National Natural Science Foundation of China (#92057204, #82088102, #32070753, #31900542, #91854208, and #31922034), the Fundamental Research Funds for the Central Universities (#20720200069), the Project “111” sponsored by the State Bureau of Foreign Experts and Ministry of Education of China (#BP2018017), the XMU Training Programme of Innovation and Entrepreneurship for Undergraduates (2021X1183 and 2022Y123), and the Agilent Applications and Core Technology - University Research Grant (#4769).

## Author contributions

C-S.Z. and S.-C.L. conceived the study and designed the experiments. M.L. identified PDZD8 as an AMPK substrate, and Yu W. and W.-F.C. identified T527 is the site of PDZD8 for AMPK phosphorylation. M.L. and X.W. discovered the roles of AMPK-PDZD8 in promoting contact between ER and mitochondria. M.L. identified that PDZD8 is required for the utilisation of glutamine in cells, and Yongliang W. in mouse muscles. W.-F.C. identified the roles of PDZD8 in enhancing GLS1 activity with the help from B.J. and Q.L. X.W. performed OCR analysis in cells and muscle tissues, with the assistance from M.Z. and G.L., respectively. Yu W. performed nematode experiments (with the help from Y.Y.), and analysed the rejuvenation roles of PDZD8 in aged mouse. M.Z. analysed the levels of TCA cycle intermediate by GC-MS. L.Y. performed 2D-SEM. Y.-H.L. performed the domain mapping experiments, and discovered the autoinhibition of PDZD8, with the assistance from S.C. J.W. generated the muscle-specific, PDZD8-527A-expression mouse. J.X., X.T., and Q.Q. generated the knockout cell lines. R.X. and X.L. performed FIB-SEM with the help from L.Y. X.H. generated the anti-p-T527-PDZD8 antibody. C.X., Yaying W. and Z.X. performed the protein mass spectrometry. C.Z. analysed adenylates, NAD^+^, glutamine, and malonyl-coA levels through CE-MS and HPLC-MS. B.Z. and X.D. performed the in silico modelling assays. Z.-C.W. and H.-L.P. helped interpret the metabolomic data. S.-Y.L. helped supervise the project. C.-S.Z. and S.-C.L. wrote the manuscript.

## Additional information

### Supplementary information

The online version contains supplementary material available at https://doi.org/10.1038/.

### Correspondence and requests for materials

should be addressed to Sheng-Cai Lin.

### Peer review information

Nature thanks anonymous reviewer(s) for their contribution to the peer review of this work. Peer reviewer reports are available.

### Reprints and permissions information

is available at http://www.nature.com/reprints.

## Supplementary Notes

### Supplementary Note 1

AMPK promotes the formation of ER-mitochondrial contact, based on the following lines of evidence: a) Subcellular fractionation yielded smaller amounts of pure mitochondria from glucose-starved cells and tissues than those normally cultured or fed (see methods section of ref. ^53^). Detailed analysis revealed that more mitochondria from starved cells or tissues were associated with ER, as increased amounts of mitochondria were present in the mitochondria-associated ER membrane (MAM)^161^, and knockout of *AMPK*α blocked this effect (Fig. 1a); b) transmission electron microscopic (TEM) images showed a significantly increased proportion of ER-associated mitochondria (Fig. 1b and Extended Data Fig. 1a; defined as membrane appositions between the two organelles with less than 30 nm distance, as reviewed in ref. ^174^), a reduced average distance between ER and mitochondria (Extended Data Fig. 1b), and an increased number of contact sites within a cell (Extended Data Fig. 1c), all of which are AMPK-dependent; c) cyro-focused ion beam scanning electron microscopy (FIB-SEM) images gave an AMPK-dependent increase of volumes (and surface area) of the ER-mitochondrial contacts (Fig. 1c, d); and d) live-cell imaging using a split-GFP-based ER-mitochondrial contact site sensor (the short-range SPLICS^175^) suggested an increase of 8-10 nm distance between the E{R and mitochondria, which is also dependent on AMPK (Fig. 1e). As controls, we did not observe significant change of ER and mitochondrial morphology, including the “aspect ratios” of mitochondria, which is a parameter/factor invented to describe the morphology of a mitochondrion (defined as the length of the major axis divided by the length of the minor axis)^153,154^ (as assessed by TEM; Extended Data Fig. 1d), cristae lengths of mitochondria (assessed by TEM; Extended Data Fig. 1d), and the volumes and surface areas of ER and mitochondria (assessed by FIB-SEM; Extended Data Fig. 1e).

### Supplementary Note 2

As described previously^54–57^, a typical AMPK substrate motif meets at least one of the following properties:

a) A hydrophobic residue (valine, isoleucine, leucine, methionine, phenylalanine, tryptophan and cysteine) in −5 position relative to the phosphoacceptor site (serine or threonine), and the following sites on PDZD8 meet this criteria: S10, T18, T135, S144, T153, T159, T171, S215, S223, T234, S269, T284, T300, S338, S352, S354, S386, T427, S503, S530, T662, S666, S672, S733, T746, T749, T790, S801, S822, S867, S889, T912, S925, S943, T961, S989, S1056, T1103, S1106, S1132 and S1153. We also included those residues show less hydrophobic properties at −5 position (alanine, tyrosine, histidine, threonine, serine, proline and glycine): S15, T79, T81, T86, T91, T94, T101, T120, T239, S244, T268, T326, S331, T348, S353, S403, T425, S426, S476, T489, S491, S496, T527, S538, S558, T569, S570, S631, S663, S673, T678, T696, S699, S747, T767, T769, S775, T837, S842, T894, S927, T932, S957, T971, S980, S991, S996, S1011, T1029, S1080, S1108, S1113, S1137, S1142, and S1144.

b) A basic residue (arginine, lysine, histidine) in −4 position, and sites T120, T234, T239, S244, T284, T326, S363, S386, T427, S497, T527, T528, S663, T696, S699, T767, S801, T807, T863, T901, T941, S957, T974, S996, T997, T1029, and S1074 conform to this criterion.

c) A basic residue in −3 position, and sites T268, S269, T284, T300, T319, T326, S331, S338, T348, S352, S353, S354, S362, S363, T366, S376, T380 and S386 conform to this criterion.

d) A hydrophobic residue in +4 position, and sites T425, S426, T427, S452, S471, S472, S473, S476, T486, T489, S491, S497, S503, S517, S519, S521, T527, T528, S530, S538, S549, S558, T569, S570, T573, S579, T582, S585, S603, S631, T662, S663, T665, S666, T669, S672, S673, T678, S681, S682, T694, T696, S699, S733, T746, S747, T749, S753, S761, T767, T769, S775, T790, S801, T807, S822, T837, S842, T846, T852, T863, S867, T888, S889, T894, T901, T912, S925, S927, T931, T932, T935, T941, S942, S943, S952, T954, S957, T961, S967, T971, T974, S975, S980, T982, S989, S991, S996, T997, S1011, T1029, S1056, T1064, T1065, T1067, S1071, S1074, S1080, T1088, and T1103 conform to this criterion.

Among these predicted sites, S10, S15, T18, T81, T101, T120, T135, S144, T153, T159, T171, S215, S223, T234, T239, S244, T319, T326, S331, S338, T348, S352, S362, S363, T366, S376, T380, S386, T425, S426, T427, S496, S519, S521, T527, T528, S530, S538, S549, S558, S579, S585, T696, S733, S747, S753, S761, T767, T769, S775, T790, S822, S842, T894, T901, S927, T931, T935, T941, S942, S943, S952, T954, S967, S996, S1011, T1029, S1056, T1064, S1071, S1074, S1080, T1088, T1103, S1108, S1113, S1132 and S1144 were hit in mass spectrometry analysis (Supplementary Table 1; all converted to human PDZD8 amino acid positions) and were evolutionarily conserved in human. After individually mutating these sites and the other predicted sites as well, we found that T527 is the site of PDZD8 that is phosphorylated by AMPK (Fig. 1h, i). T527 also fits the AMPK substrate motif refined by a very recent study^58^, in which a hydrophobic residue resides in the −2 position, and a proline residue in −1.

### Supplementary Note 3

We conclude that under glucose starvation, the utilisation of glutamine through glutaminolysis was promoted, based on following two lines of evidence:

a) In a dynamic labelling assay in which MEFs were pre-treated with [U-^13^C]glutamine for 20 min, a time duration lay in a phase during which the ^13^C incorporation in the pool of TCA cycle intermediates was increased in a time-dependent manner (not saturated; see “Determination of rates of glutaminolysis and FAO” in Methods section), the levels of ^13^C-isotopologs of TCA cycle intermediates were increased (Fig. 2a), indicating the overall rates of glutamine carbon entry into the TCA cycle were increased.

b) The deamination reaction, which reflects the conversion of glutamine to glutamate and then α-KG catalysed separately by GLS1, and glutamic-pyruvic transaminase 2 (GPT2) and glutamic-oxaloacetic transaminase 2 (GOT2), as determined by the levels of ^15^N-labelled alanine and aspartate (both are m+1) in MEFs pre-treated with [alpha-^15^N]glutamine, was significantly promoted in low glucose (Extended Data Fig. 4d). Furthermore, knockdown of *GLS1* or treatment of GLS1 inhibitor BPTES blocked the effects of enhanced glutaminolysis on OCR (Fig. 2q, Extended Data Fig. 5m). These data indicate that the channelling of glutamine to the TCA cycle though glutaminolysis is promoted in low glucose.

Data shown in Fig. 2a and Extended Data Fig. 4b also indicate an elevated, glutamine-derived cataplerosis from the TCA cycle in low glucose, including an elevated reductive carboxylation (determined by the levels of m+5 and m+3 citrate), an elevated citrate-pyruvate cycle (determined by the levels of m+3 malate), and an elevated malate-aspartate shuttle (determined by the levels of m+4 aspartate). The cataplerosis-mediated dissipation of the TCA cycle intermediates prevents the accumulation of anions in the mitochondrial matrix brought about by the increased glutaminolysis, which may inhibit TCA reactions (reviewed in ref. ^176^), thereby sustaining the high rates of TCA reactions observed in Fig. 2a.

We found that FAO was promoted much later than that of glutaminolysis in low glucose, as determined by levels of [U-^13^C]palmitate-labelled, ^13^C-isotopologs of TCA cycle intermediates during the starvation periods. One may argue that it is the utilisation of stored (unlabelled) TAG first, and labelled palmitate next, in low glucose, that may lead to the delayed elevation of ^13^C-isotopologs of TCA cycle intermediates. We therefore determined the levels of free glycerol in culture medium to reflect the rates of lipolysis in MEFs, and found that glucose starvation did not elevate free glycerol contents (Extended Data Fig. 4e), ruling out the possibility that stored TAG utilisation leads to the delayed promotion of FAO in low glucose. Consistently, we have shown that knockout of *CPT1* or treatment of CPT1 inhibitor etomoxir in low glucose did not block the promotion of glutaminolysis and OCR (Fig. 2r, Extended Data Fig. 5m).

### Supplementary Note 4

To determine the carbon source shift during low glucose, we separately labelled MEFs with [U-^13^C]glutamine, [U-^13^C]palmitate, [U-^13^C]pyruvate, and [U-^13^C]glucose (only in high glucose condition) until isotopic enrichment has reached steady states (been saturated, see “Determination of rates of glutaminolysis and fatty acid oxidation (FAO)” of Methods section, and ref. ^137,138^ for glutamine, glucose and pyruvate labelling, and ref. ^134^ for PA labelling), and then determined the contribution of these carbon sources to the pool of TCA cycle intermediates. As shown in Fig. 2c, in high glucose, the contribution of glucose, glutamine, palmitate and pyruvate to the TCA cycle was approximately 15%, 54%, 8% and 23%, respectively. Under 2-h glucose starvation, glutamine contributes more than 72% pool of the TCA cycle intermediates, while pyruvate 24%, and palmitate 10%.

### Supplementary Note 5

As shown in Extended Data Fig. 7a, the truncate PDZD8-CT lacking the N-terminal region showed a significantly higher affinity towards GLS1 than that of full-length PDZD8, indicative of an intramolecular autoinhibition of the C-terminus of PDZD8 by the N-terminus for interacting with GLS1. Indeed, the truncate protein PDZD8-NT showed a strong interaction with PDZD8-CT regardless of glucose concentrations (Extended Data Fig. 8c). We also found that phosphorylation of full-length PDZD8 by AMPK led to an increased affinity towards GLS1, to a similar extent to that between PDZD8-CT and GLS1 (Fig. 4i), suggesting that phosphorylation of T527 removes the intramolecular autohinibition. The FRET-FLIM experiment in live cells also indicated that the N-terminus of PDZD8 was moved away from the its C-terminus under glucose starvation, as the fluorescent lifetimes of GFP fused to the N-terminus of PDZD8 were significantly increased, due to the removal of FRET brought about by mCherry fused to PDZD8-NT (Fig. 4j); knockout of *AMPK*α, or re-introduction of PDZD8-T527A mutant into *PDZD8*^−/−^ cells abolished the low glucose-induced conformational change of PDZD8 (Fig. 4j). Given that PDZD8 is anchored in the ER membrane through its transmembrane (TM) domain located near the N-terminus^60^, and that the C-terminus of PDZD8 participates in interacting with the mitochondrion-localised GLS1, it is reasonable to suggest that GLS1 might be involved in PDZD8-mediated promotion of ER-mitochondria contact in low glucose. Indeed, we found that knockdown of *GLS1* significantly blocked the glucose starvation-induced tightening of the contact between ER and mitochondria (Fig. 4k, l, Extended Data Fig. 8d-g). Together, we stand to reason that upon phosphorylation at T527, the C-terminal region of PDZD8 is no longer inhibited by the N-terminus, and exhibits stronger affinity for GLS1, which consequentially promotes GLS1 activity along with tightened tethering of mitochondria to ER.

### Supplementary Note 6

In mitochondria, GLS1 has been reported to be localised on both the external^87–89^ and internal sides of IMM^177,178^, as well as the mitochondrial matrix^76,89,179^. To determine the pool of GLS1 that interacts with PDZD8 in low glucose, we performed the APEX2 proximity labelling experiments^180^ using MEFs stably expressing a chimera between the biotinylating enzyme ascorbate peroxidase 2 (APEX2) fused to the C-terminus of PDZD8 under the control of a doxycycline-inducible promoter. We found that a significant enrichment of biotinylated GLS1 in purified IMM from starved cells, while GLS1 was hardly biotinylated in the purified mitochondria matrix regardless of starvation (Extended Data Fig. 10e). If PDZD8-APEX2 interacted with GLS1 localised on the internal side of IMM, the matrix GLS1 may probably be biotinylated, but this did not happen. Given this, we conclude that PDZD8 interacts with GLS1 located on the external side of IMM. Interestingly, it has been suggested that the enzymatically active GLS1 is localised on the outer face of the IMM, because the high concentrations of glutamate in the matrix will inhibit the GLS1 localised in the internal sides of IMM and matrix ^87,89,181–183^. As for how PDZD8 approaches the outer face of IMM, it is reasonable to speculate that PDZD8 likely penetrates across through the outer mitochondrial membrane, given that the ER-mitochondria contact site is closely associated with the protein sorting and assembly machinery (SAM) of mitochondria. In yeast, for example, the ERMES integral member Mdm10 is also a component of the SAM complex on the OMM^184,185^. Therefore, the promotion of ER-mitochondria contact may facilitate the penetration of PDZD8 through the OMM to interact with GLS1, leading to its activation, which is, indeed observed by us (Extended Data Fig. 10f).

## Extended Data figure legends

**Extended Data Fig. 1.**
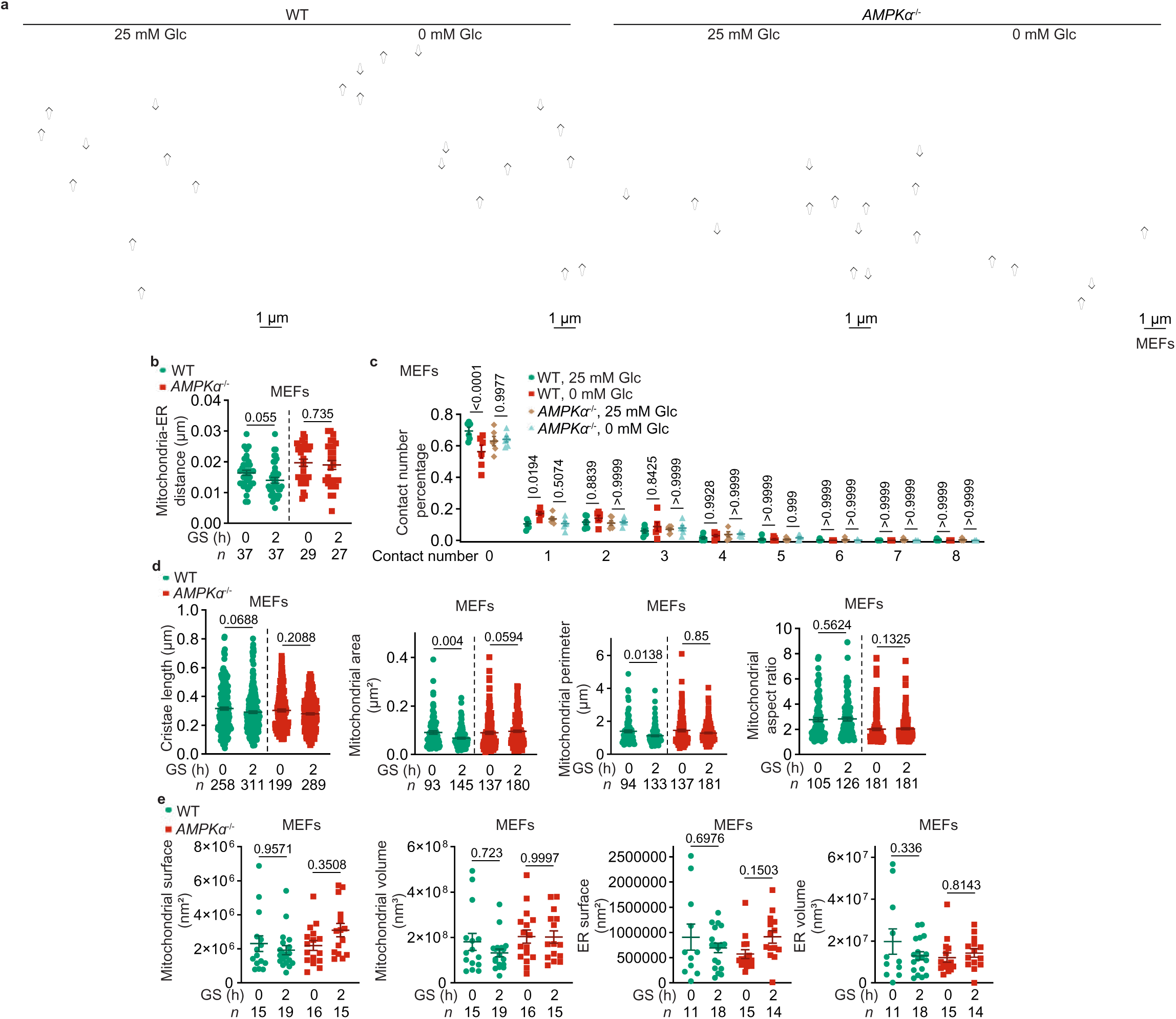
AMPK promotes ER-mitochondria association. **a**, Representative TEM images of Fig. 1b. The white arrowheads point to ER-mitochondria contacts (defined as membrane appositions between the two organelles with less than 30 nm distance). **b**-**d**, Statistical analysis of TEM images in **a**. The average distance of each mitochondrion between the nearest ER (**b**), the numbers of ER that form contacts with a specific mitochondrion (**c**; shown as the percentage of each “kind”, i.e., contact with 0, 1, 2, 3…or 8 pieces of ER sheets, of mitochondria), the total length of cristae of a mitochondrion (**d**), the area and the perimeter of a mitochondrion (**d**), and the aspect ratio (calculated as described in “TEM and FIB-SEM” of Methods section) of a mitochondrion (**d**) were calculated according to the TEM images. Data are shown as mean ± s.e.m.; *n* = 6 (**c**) cells, or labelled on each panel representing contact numbers **(b)** or mitochondria numbers (**d**, except the leftmost panel cristae) for each condition. **e**, Statistical analysis of FIB-SEM images in Fig. 1c. The surface area (left panel) or volume (right panel) of a mitochondrion to the ER that forms a contact with, were determined. Data are shown as mean ± s.e.m.; *n* values represent mitochondria numbers for each condition. Note that mitochondria and ER partially covered in the image/field were not calculated. *P* values in this figure were determined by Mann-Whitney test (**b**, KO and **d**), unpaired two-tailed Student’s *t*-test (**b**, WT), or two-way ANOVA, followed by Sidak (c) or Tukey (**e**). Experiments in this figure were performed three times.

**Extended Data Fig. 2.**
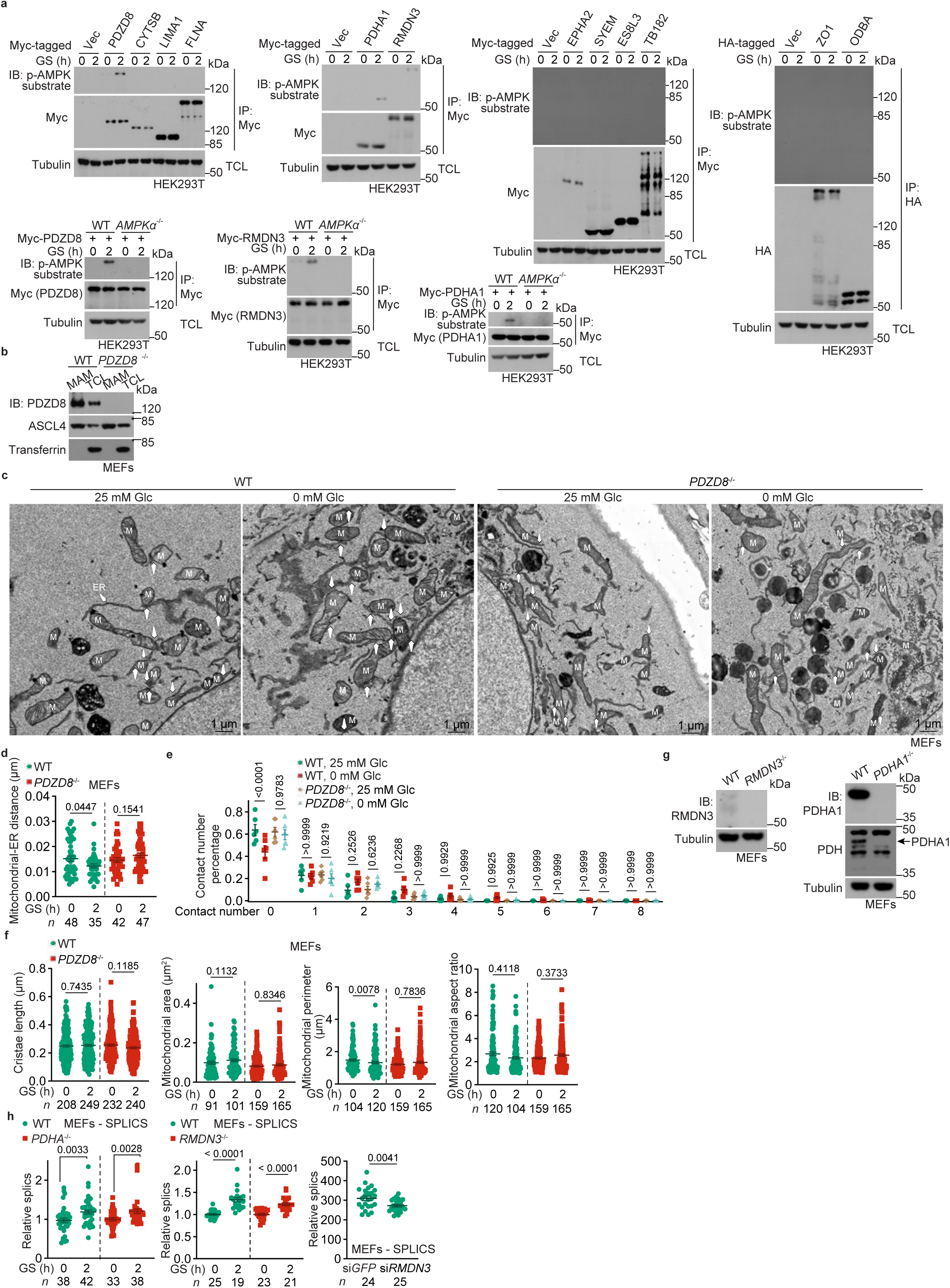
PDZD8 is a substrate of AMPK. **a**, Verification of possible AMPK substrate(s) in MAM. In the upper panel, HEK293T cells transfected with different constructs of potential AMPK substrates (HA- or Myc-tagged) hit by mass spectrometry (listed in Supplementary Table 1) were glucose-starved for 2 h, followed by immunoprecipitation using antibodies against the HA-tag or Myc-tag, and followed by immunoblotting using the antibody for pan-phospho-AMPK-substrates. In the lower panel, wildtype and *AMPK*α^−/−^ HEK293T cells were transfected with Myc-tagged, PDZD8, RMDN3 and PDHA1, three phosphoproteins hit by the mass spectrometry, followed by glucose starvation, immunoprecipitation and immunoblotting as in the upper panel. **b**, Validation of *PDZD8* knockout in MEFs. Cells were lysed or subjected for the purification of MAM, followed by immunoblotting. **c**, Representative TEM images of Fig. 1f. **d**-**f**, Statistical analysis of TEM images in **c**. Data were analysed as in Extended Data Fig. 1b-d, and are shown as mean ± s.e.m.; *n* = 6 cells (**e**), or labelled on each panel indicate contact numbers (**d**) or mitochondria numbers (**f**, except the leftmost panel cristae) for each condition. **g**, Validation of *RMDN3* and *PDHA1* knockout MEFs. **h**, *RMDN3* and *PDHA1* do not regulate the formation of ER-mitochondria contact. *PDHA1*^−/−^ MEFs (left panel), *RMDN3*^−/−^ MEFs (middle panel), or wildtype MEFs with *RMDN3* knocked down (right panel), all stably expressing SPLICS reporter, were glucose-starved for 2 h, followed by determination of the contact formation through quantifying the puncta of SPLICS as in Fig. 1e. Data are shown as mean ± s.e.m.; *n* (labelled on each panel) values represent cell numbers for each condition. *P* values in this figure were determined by unpaired two-tailed Student’s *t*-test (**d**, KO cells of middle panel and the right panel in **h**), by Mann-Whitney test (**f**, left panel and WT cells of the middle panel in **h**), or by two-way ANOVA, followed by Sidak (**e**). Experiments in this figure were performed three times.

**Extended Data Fig. 3.**
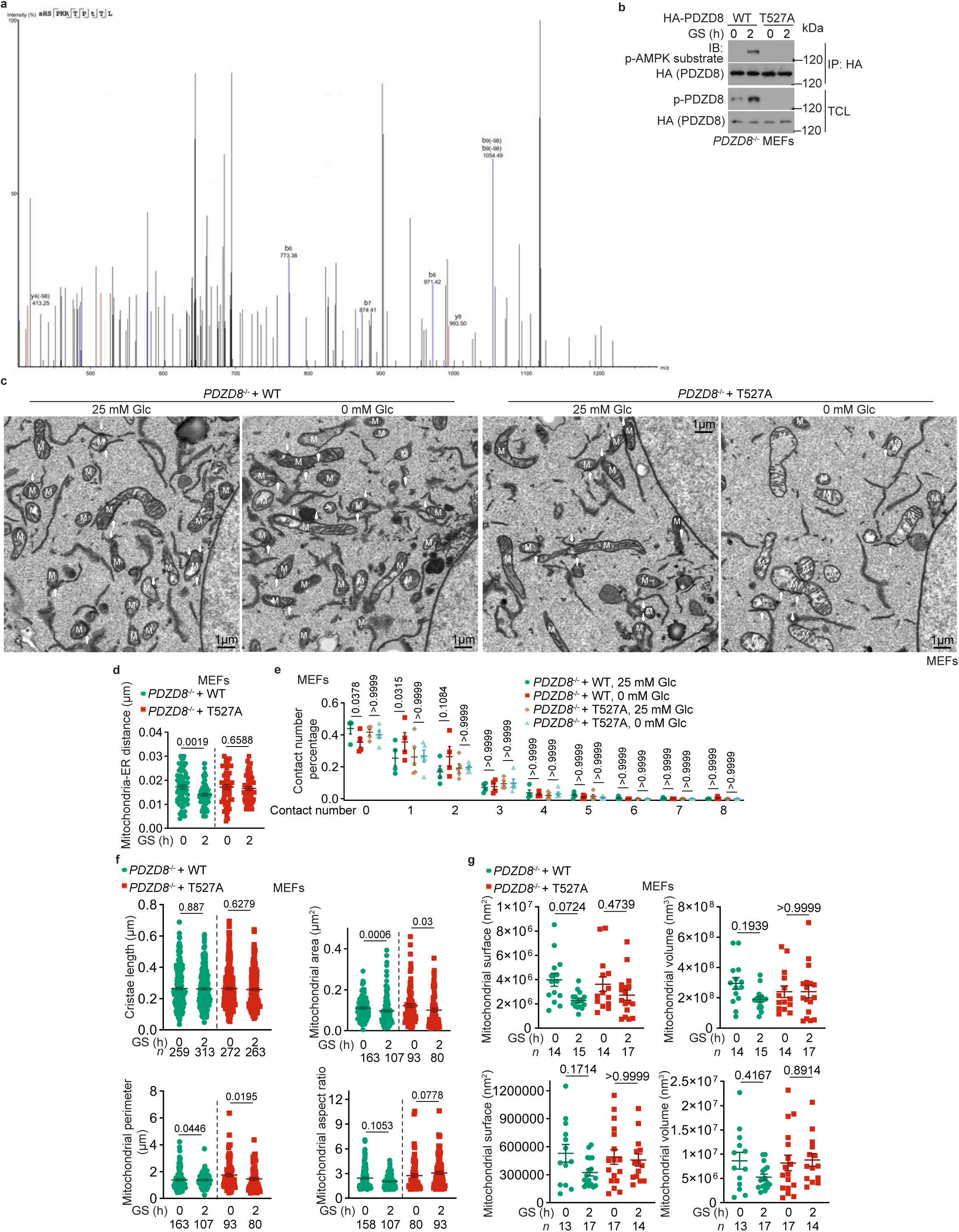
AMPK phosphorylates PDZD8 at T527 residue. **a**, Typical spectrogram showing that T527 site of PDZD8 is phosphorylated. **b**, Validation of p-T527-PDZD8 antibody. *PDZD8*^−/−^ MEFs stably expressing HA-tagged PDZD8 or PDZD8-T527A were glucose-starved for 2 h, followed by immunoblotting using the p-T527-PDZD8 antibody. As a control, HA-tagged PDZD8 was immunoprecipitated, followed by immunoblotting using the pan-phospho-AMPK-substrates antibody. **c**, Representative TEM images of Fig. 1m. **d**-**f**, Statistical analysis of TEM images in **c**. Data were analysed as in Extended Data Fig. 1b-d, and are shown as mean ± s.e.m.; *n* = 6 mitochondria (**e**), or labelled on each panel, contact numbers (**d**) or mitochondria numbers (**f**, except the upper left panel cristae) for each condition. **g**, Statistical analysis of FIB-SEM images in Fig. 1n. Data were analysed as in Extended Data Fig. 1e, and are shown as mean ± s.e.m.; *n* (labelled on each panel) values represent mitochondria numbers for each condition. *P* values were determined by unpaired two-tailed Student’s *t*-test (**d**), by Mann-Whitney test (**f**), or two-way ANOVA, followed by Sidak (**e**) or Tukey (**g**). Experiments in this figure were performed three times.

**Extended Data Fig. 4.**
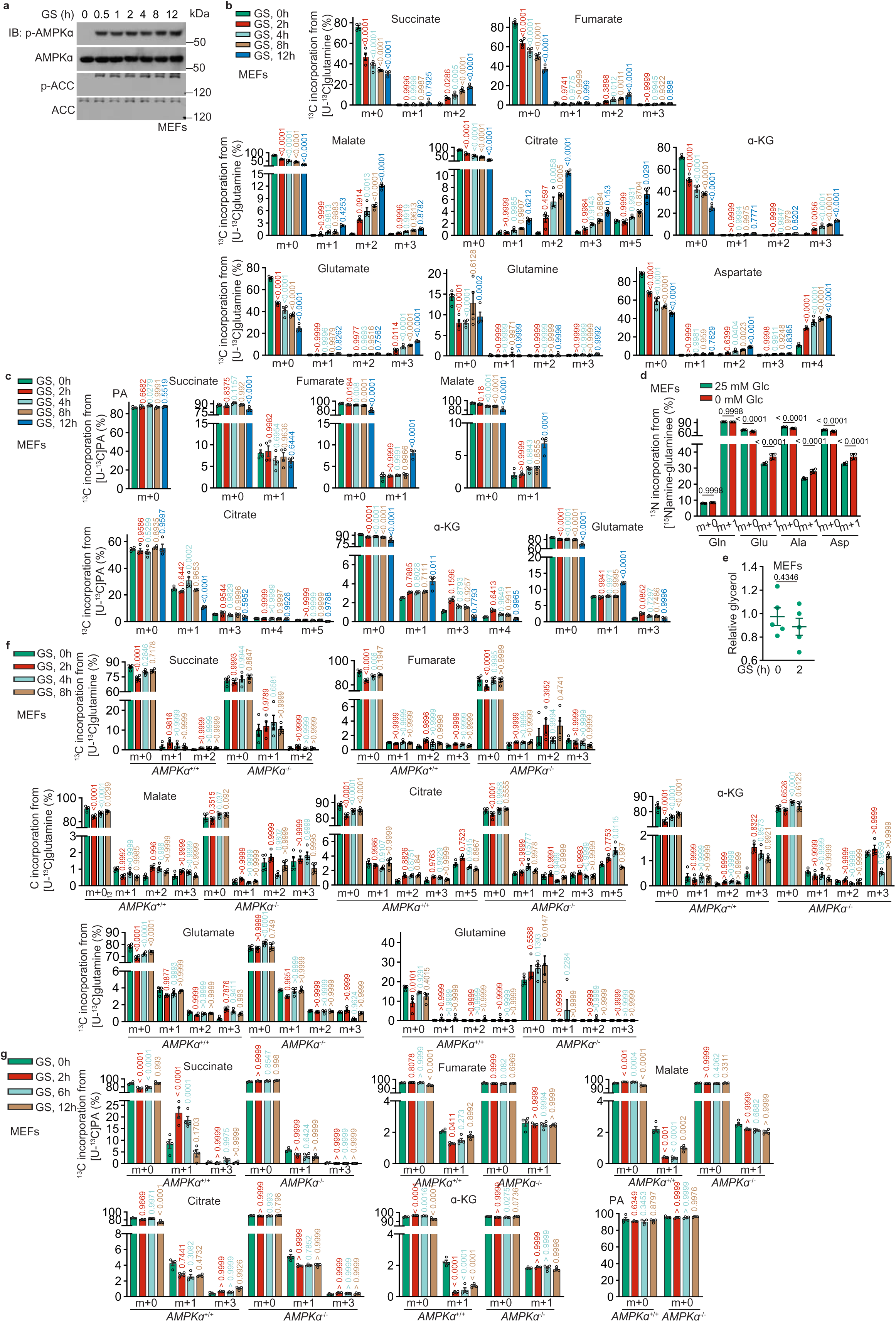
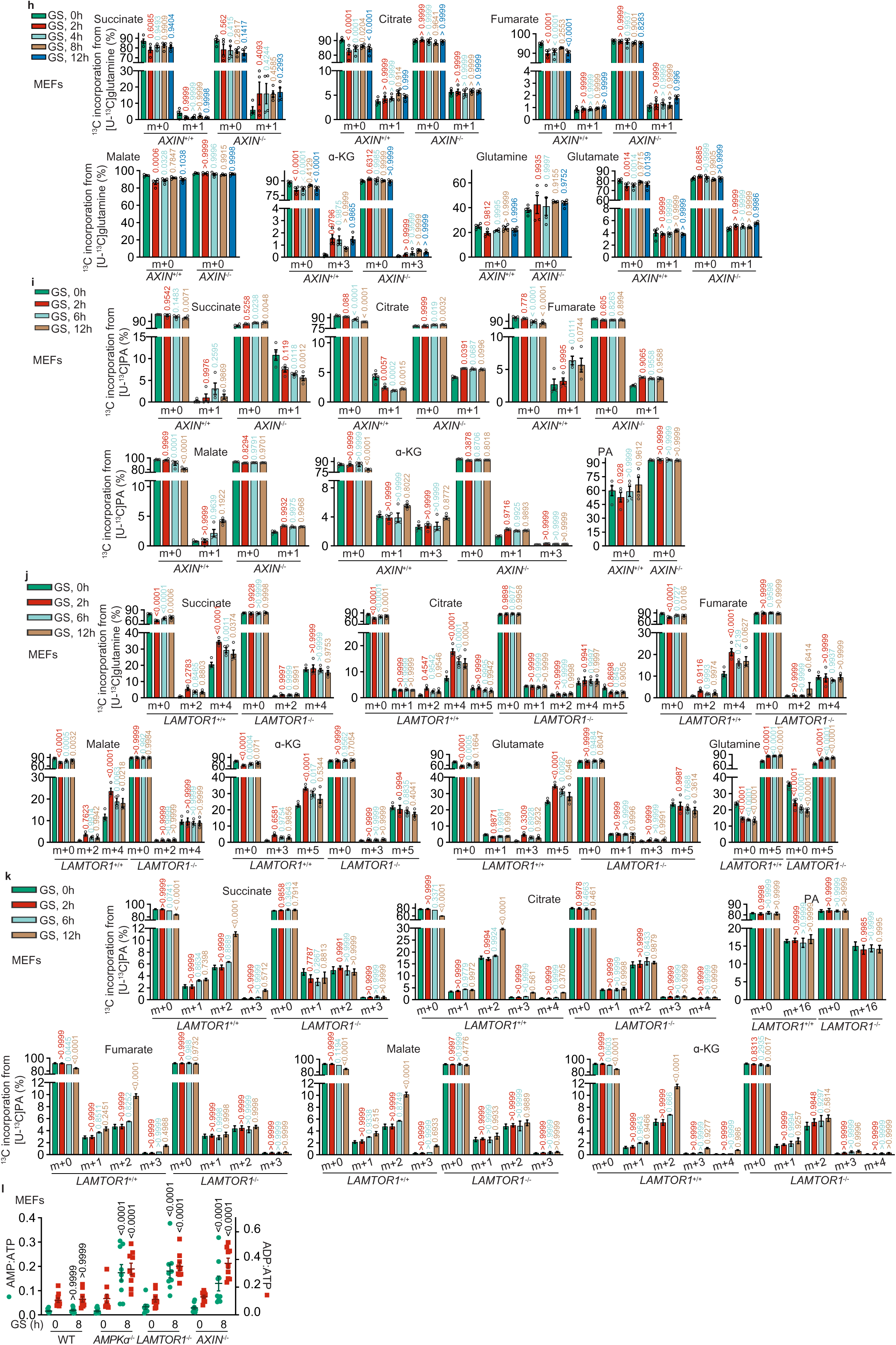
Lysosomal AMPK promotes the utilisation of glutamine. **a**, Glucose starvation leads to a fast and persistent activation of AMPK. MEFs were glucose-starved for desired durations, followed by immunoblotting for p-AMPKα and p-ACC. **b**, **c**, **f**-**i**, Levels of other isotopomers of the labelled TCA cycle intermediates shown in Fig. 2a (**b**), 2b (**c**), 2d (**f**), 2e (**h**), 2f (**g**) and 2g (**i**). Data are shown as mean ± s.e.m.; *n* = 4 for each condition; and *P* values were determined by one-way ANOVA, followed by Tukey (PA of **c**, **g**, **i**; and malate and glutamine of **h**), or two-way ANOVA, followed by Tukey (others), all compared to the unstarved group of each genotype. **d**, Deamination reaction is promoted under glucose starvation. MEFs were glucose starved for 2 h. At 20 min before sample collection, cells were labelled with [alpha-^15^N]glutamine, followed by determination of the levels of m+1 glutamate (Glu), alanine (Ala) and aspartate (Asp), all indicators to the rates of deamination reactions, along with glutamine (Gln). Data are shown as mean ± s.e.m.; *n* = 4 for each condition; *P* values were determined by two-way ANOVA, followed by Tukey. **e**, Glucose starvation does not promote lipolysis in MEFs. MEFs were glucose starved for 2 h, followed by determining free glycerol in culture medium to reflect the rates of lipolysis. Data are shown as mean ± s.e.m.; *n* = 5 biological replicates for each condition; *P* values were determined by unpaired two-tailed Student’s *t*-test. **j**, **k**, LAMTOR1 is required for the promotion of glutamine utilisation. Experiments were performed as in Fig. 2a (**h**) and 2b (**i**), respectively, except that *LAMTOR1*^−/−^ MEFs were used. Data are shown as mean ± s.e.m.; *n* = 4 for each condition; and *P* values were determined by two-way ANOVA, followed by Tukey, all compared to the unstarved group of each genotype. **l**, Ablation of lysosomal AMPK activation that blocks the promotion of both glutaminolysis and FAO in low glucose, caused energy deficiencies. MEFs with *AMPK*α, *AXIN* or *LAMTOR1* knocked out were glucose-starved for 8 h, followed by determining the AMP:ATP and ADP:ATP ratios by CE-MS. Data are shown as mean ± s.e.m.; *n* = 9 for each condition; and *P* values were determined by two-way ANOVA, followed by Tukey. Experiments in this figure were performed three times.

**Extended Data Fig. 5.**
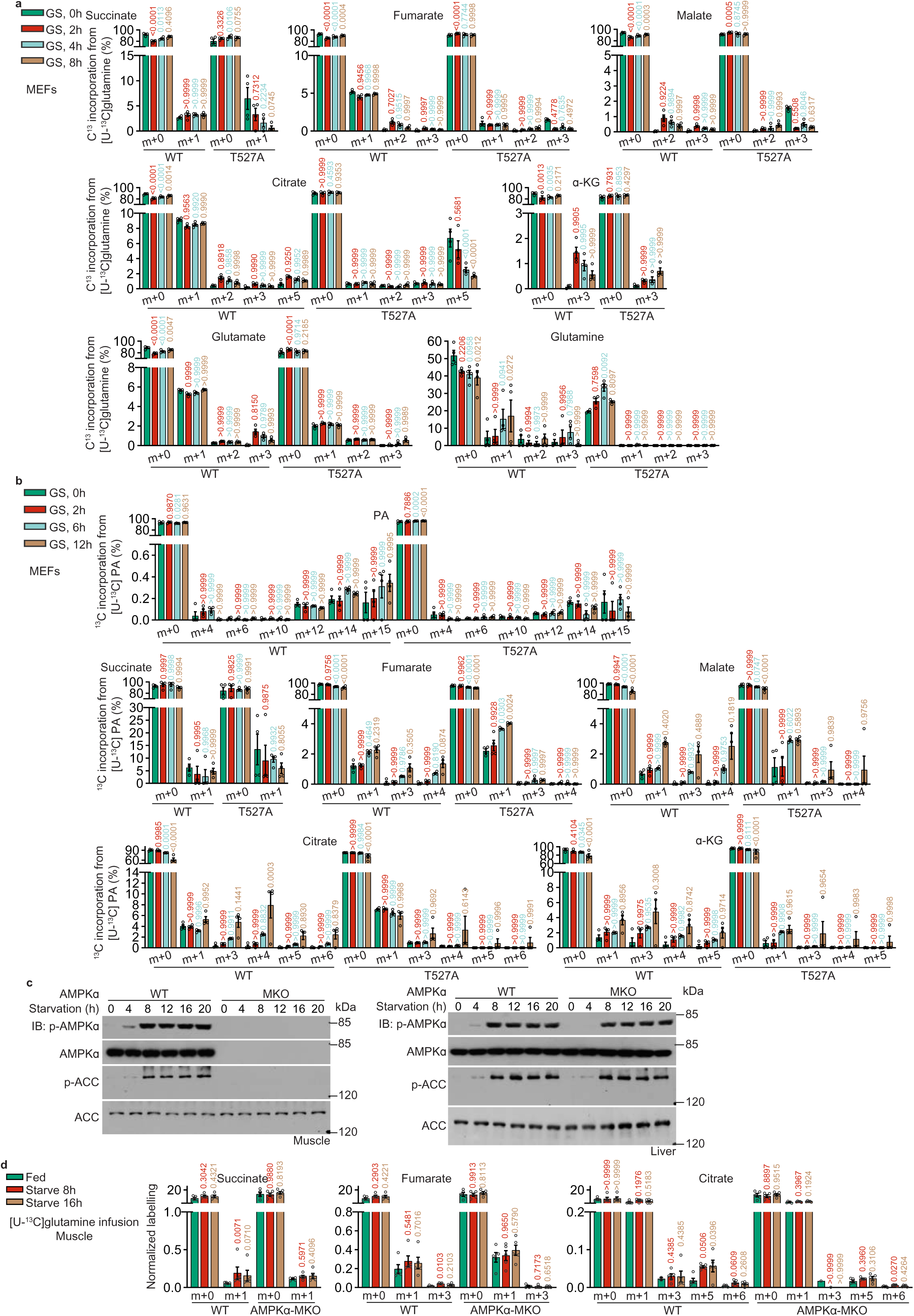
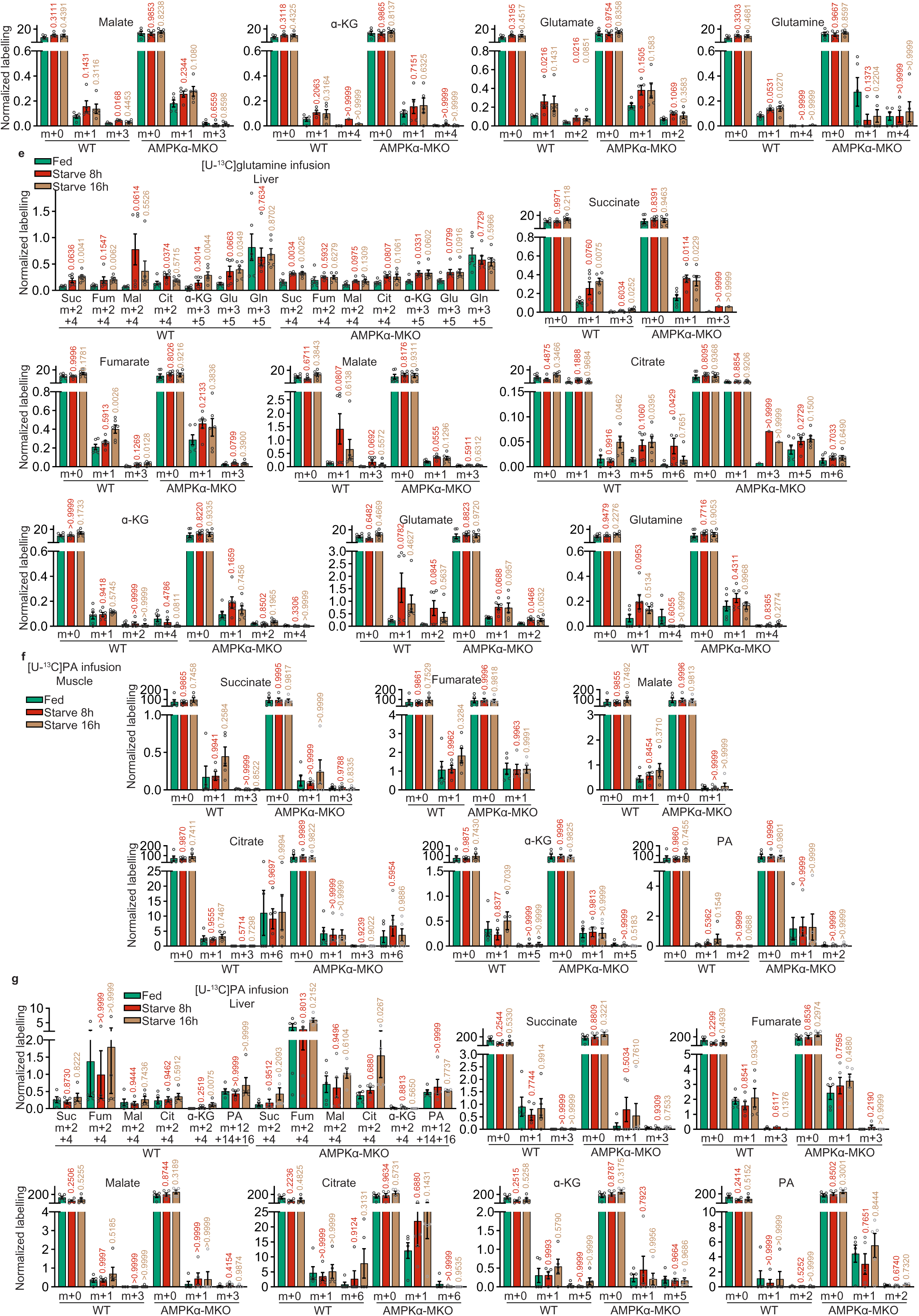
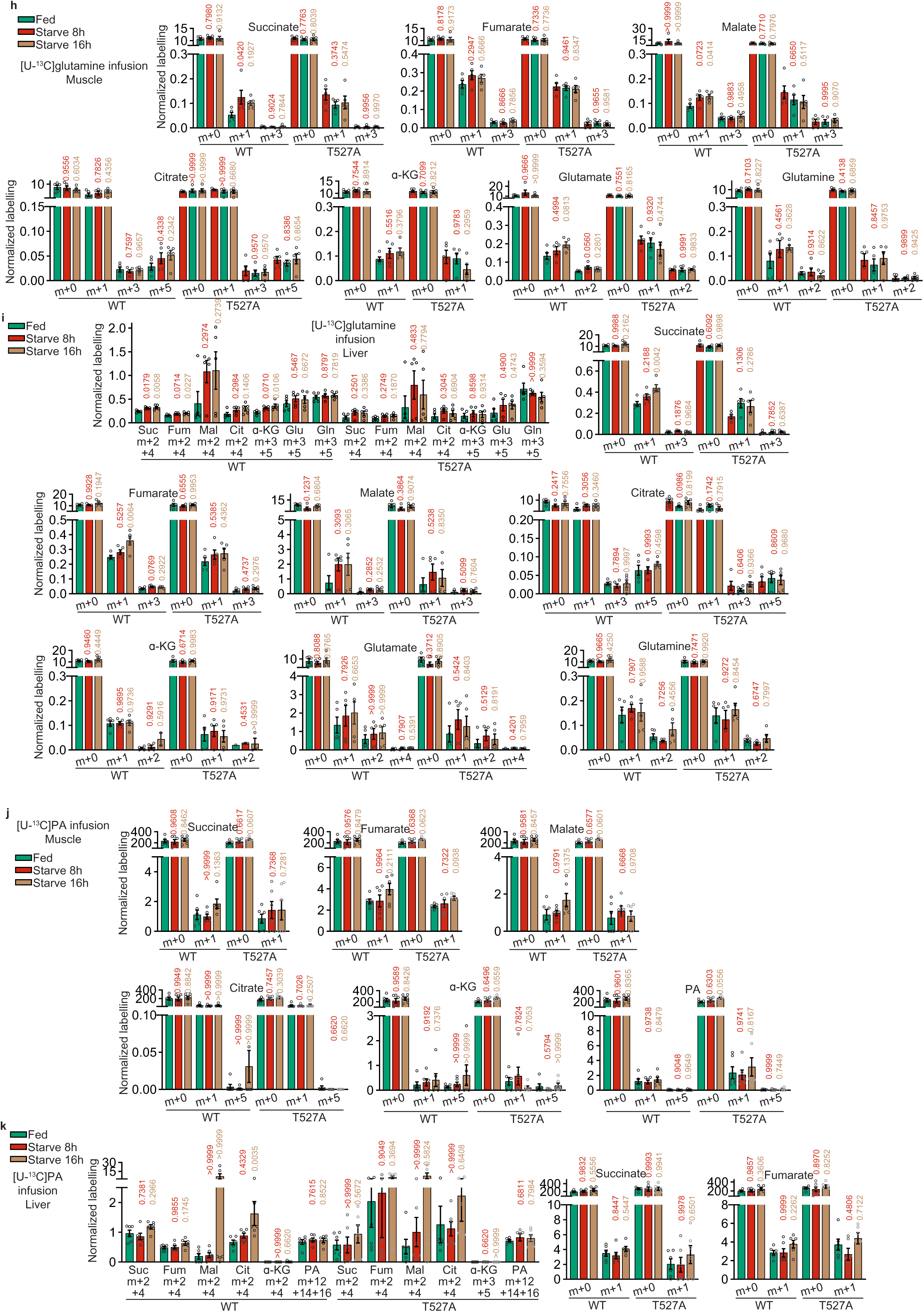
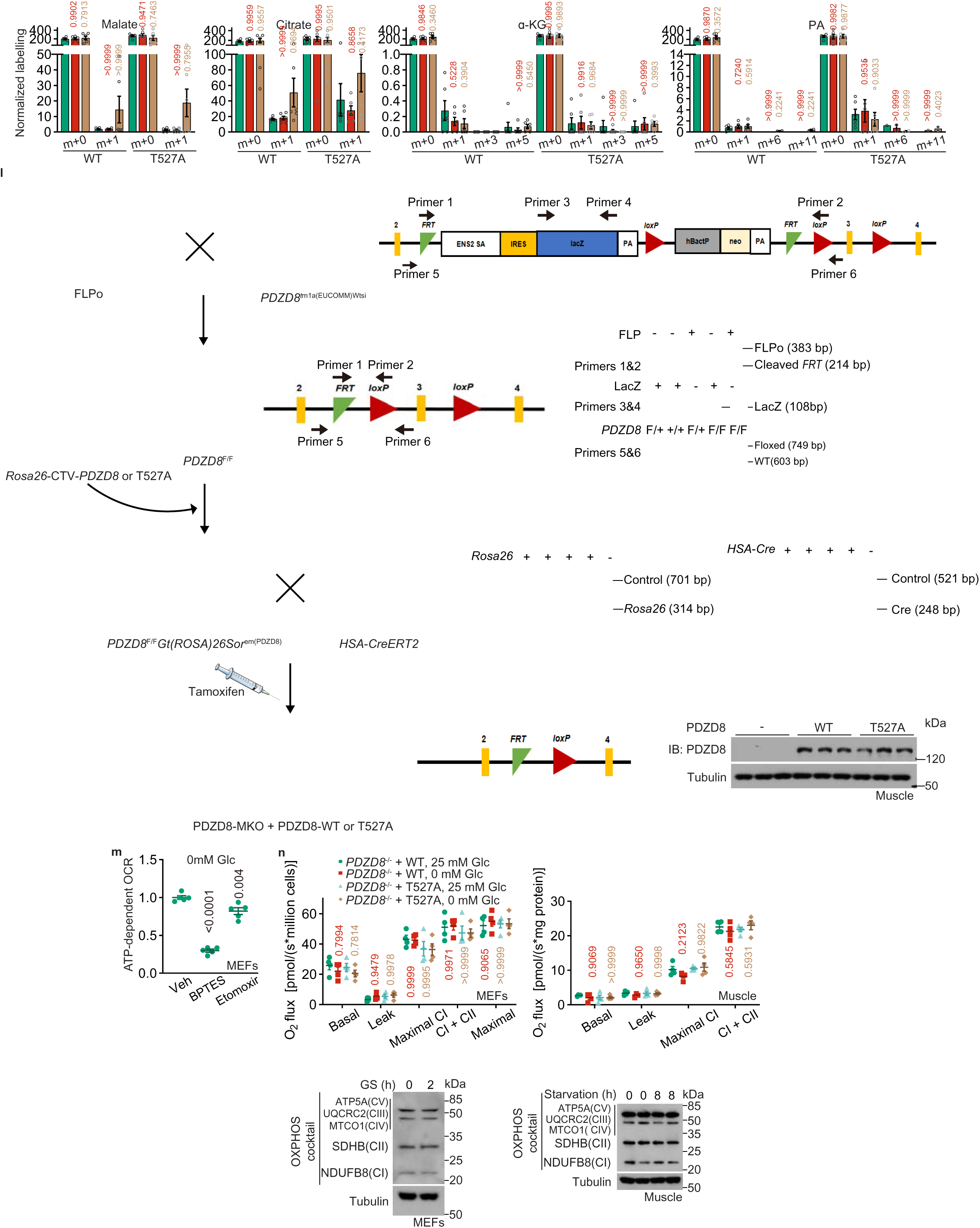
PDZD8 specifically promotes the utilisation of glutamine. **a**, **b**, Levels of other isotopomers of the labelled TCA cycle intermediates shown in Fig. 2h (**a**) and 2i (**b**). Data are shown as mean ± s.e.m.; *n* = 4 biological replicates for each condition; and *P* values were determined by two-way ANOVA, followed by Tukey, all compared to the unstarved group of each genotype. **c**, Validation of *AMPK*α-MKO mice. *AMPK*α-MKO mice were starved for desired durations, and the muscle (left panel) and liver (right panel) tissues were excised, followed by immunoblotting. **d**-**k**, Levels of other isotopomers of the labelled TCA cycle intermediates shown in Fig. 2j (**d**; see also **e** for the rates of hepatic glutaminolysis, as a control), 2k (**f**; see also **g** for the rates of hepatic FAO, as a control), 2l (**h**; see also **i** for the rates of hepatic glutaminolysis), 2m (**j**; see also **k** for the rates of hepatic FAO). Data are shown as mean ± s.e.m.; *n* = 5 (**d**, **f**, **g**, **h** and **i**), or 6 (**e**, **j** and **k**) biological replicates for each condition. *P* values in **d** were determined by: a) one-way ANOVA, followed by Dunn: succinate (m+1), citrate (m+0), malate (m+1), glutamate (m+1 and m+2) of WT MEFs in **d**; citrate (m+3) of KO MEFs in **d**; and α-KG (m+4) and glutamine (m+4) of both WT and KO MEFs in **d**; b) one-way ANOVA, followed by Sidak: citrate (m+3) of WT MEFs in **d**; and c) one-way ANOVA, followed by Tukey: others. *P* values in **e** were determined by one-way ANOVA, followed by: a) Dunn, for fumarate (m+2+4; means that the sum of m+2 and m+4 isotopomers of fumarate; same hereafter), α-KG (m+2) and glutamine (m+4) of WT mice; and α-KG (m+3+5 and m+4), glutamate (m+3+5), succinate (m+3), fumarate (m+3), and citrate (m+3) of KO mice; and b) Tukey, for others. *P* values in **f** were determined by one-way ANOVA, followed by: a) Dunn, for succinate (m+3) of WT mice; succinate (m+1), malate (m+1) and citrate (m+1) of KO mice; and α-KG (m+5) and PA (m+1 and m+2) for both WT and KO mice; and b) Tukey: for others. *P* values in **g** were determined by one-way ANOVA, followed by: a) Dunn: fumarate (m+2+4), α-KG (m+2+4 and m+5), succinate (m+3), malate (m+3) and PA (m+1 and m+2) of WT mice; citrate (m+2+4 and m+6) and malate (m+1) of KO mice; and PA (m+12+14+16), fumarate (m+3), and citrate (m+1) for both WT and KO mice; and b) Tukey: for others. *P* values in **h** were determined by one-way ANOVA, followed by: a) Dunn: malate (m+0) and glutamate (m+0) of WT mice and citrate (m+0 and m+1) of T527A mice; b) Sidak: citrate (m+3 and m+5) of T527A mice; and c) Tukey: for others. *P* values in **i** were determined by one-way ANOVA, followed by: a) Dunn: glutamate (m+2 and m+4) of WT mice; and glutamine (m+3+5) and α-KG (m+2) of T527A mice; and b) Tukey: for others. *P* values in **j** were determined by one-way ANOVA, followed by: a) Dunn: succinate (m+1) of WT mice; and citrate (m+1 and m+5) and α-KG (m+5) of both WT and KO mice; and b) Tukey: for others. *P* values in **k** were determined by one-way ANOVA, followed by: a) Sidak: fumarate (m+2+4) of T527A mice; b) Dunn: citrate (m+1) of WT mice; α-KG (m+3) of T527A mice; and malate (m+2+4 and m+1), citrate (m+2+4), α-KG (m+2+4, m+5), and PA (m+6, m+11) of both WT and T527A mice; and c) Tukey: for others. *P* values in these panels represents the comparisons between the starved and the unstarved groups of each genotype **l**, Validation of *PDZD8*-MKO mice with muscle specific reintroduction of PDZD8 or its 527A mutant. The *PDZD8*^F/F^ mice, generated through breeding the *PDZD8*-KO-first mice (*Pdzd8^tm1a(EUCOMM)Wtsi^*) with the FLPo mice (to remove the FRT-flanked Stop element ahead of the *PDZD8* locus), were validated through: a) determining FRT cleavage (the “cleaved FRT” band; genotyped through using Primers #1 and #2); and b) determining the LacZ (of the Stop element) removal (genotyped through using Primers #3 and #4). See also genotyping results for determining the existence of FLPo. The floxed PDZD8 was then validated through genotyping using Primers #5 and #6. After introducing the PDZD8 and PDZD8-T527A into the *PDZD8*^F/F^ mice through the *Rosa26*-LSL system (see “Mouse strains” of Methods section; validated by genotyping the *ROSA26* sequence), mice were bred with *HSA-CreERT2* mice (validated by genotyping the *HSA-Cre* sequence). The muscle-specific expression of PDZD8 was then induced by tamoxifen injection, followed by validation through immunoblotting. See also primer sequences and PCR programmes used for genotyping in the “Mouse strains” of Methods section. **m**, Inhibition of glutaminolysis, but not FAO, prevents OCR increases at early starvation. MEFs were pre-treated with 20 μM BPTES for 10 h, or 10 μM Etomoxir for 8 h, and then glucose-starved for 2 h (early starvation), followed by determination of OCR by Seahorse Analyzer. Data are shown as mean ± s.e.m.; *n* = 5 biological replicates for each condition; and *P* values were determined by one-way ANOVA, followed by Tukey. **n**, Glucose starvation does not affect the protein contents or the efficiency of the mitochondrial electron transport chain. MEFs or muscle tissues were permeabilised with digitonin to expose the electron transport chain, followed by addition of substrate of each mitochondrial respiratory complex to determine its activity (see “Determination of electron transport chain integrity” in Methods section; left panel; data are shown as mean ± s.e.m.; *n* = 4 for each condition); and *P* values were determined by two-way ANOVA, followed by Tukey. See also right panel for the protein levels of each mitochondrial respiratory complex before and after glucose starvation. Experiments in this figure were performed three times.

**Extended Data Fig. 6.**
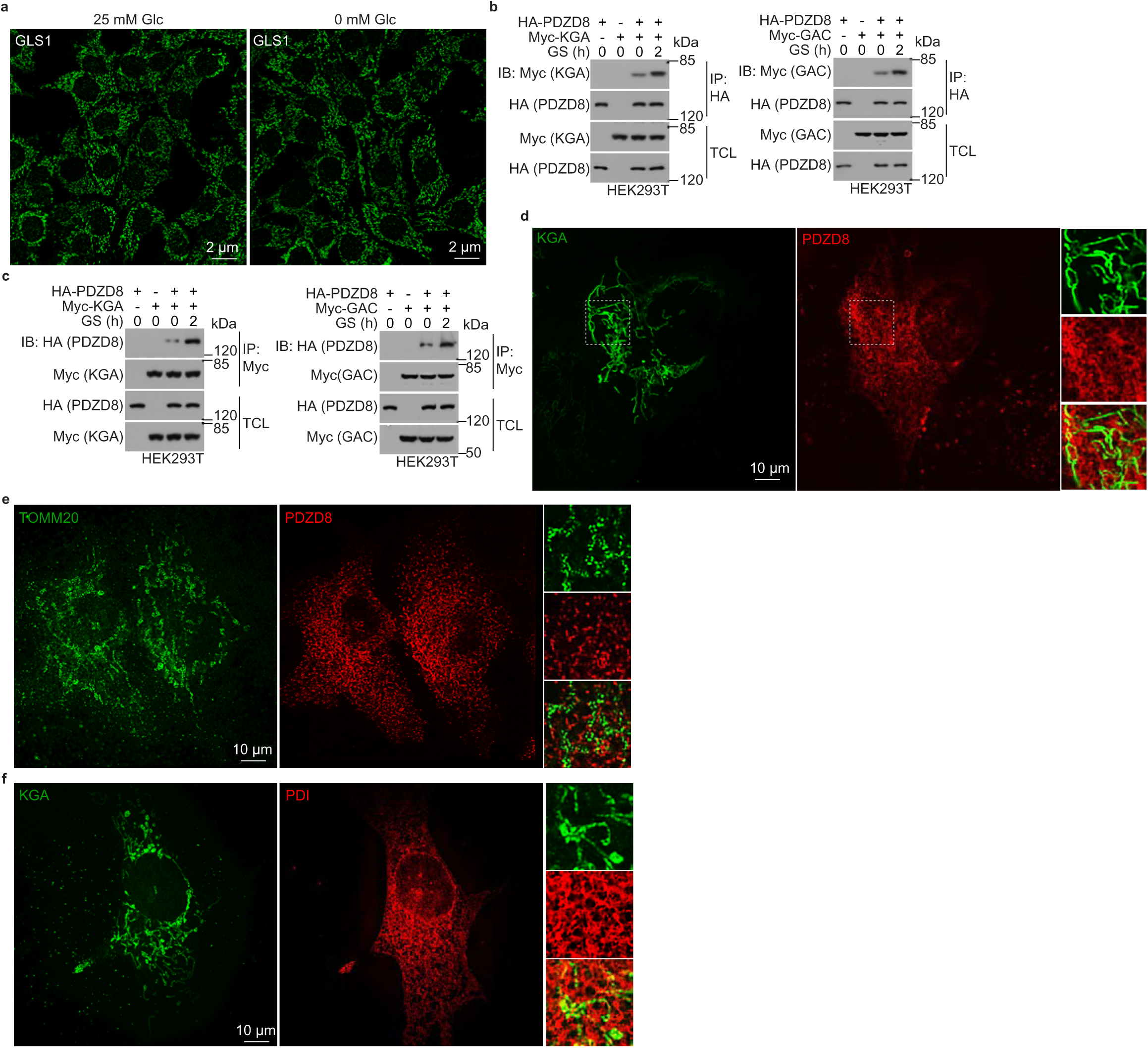
PDZD8 interacts with GLS1. **a**, Glucose starvation does not cause GLS1 filamentation. MEFs were starved for glucose for 2 h, followed by determining the filamentation of GLS1 by immunofluorescent staining. **b**, **c**, Ectopically expressed PDZD8 and GLS1 interact with each other. HEK293T cells were transfected with different combinations of PDZD8 and GLS1 (GAC or KGA), and then glucose-starved for 2 h, followed by immunoprecipitation and immunoblotting. **d**-**f**, PDZD8 is juxtaposed with GLS1 in cells. MEFs stably expressing FLAG-tagged KGA and Myc-tagged PDZD8 were stained and subjected to SIM imaging (**d**). The ER marker PDI (**f**) and the mitochondrial marker TOMM20 (**e**) were also co-stained with KGA (**f**) and PDZD8 (**e**), respectively. Experiments in this figure were performed three times.

**Extended Data Fig. 7.**
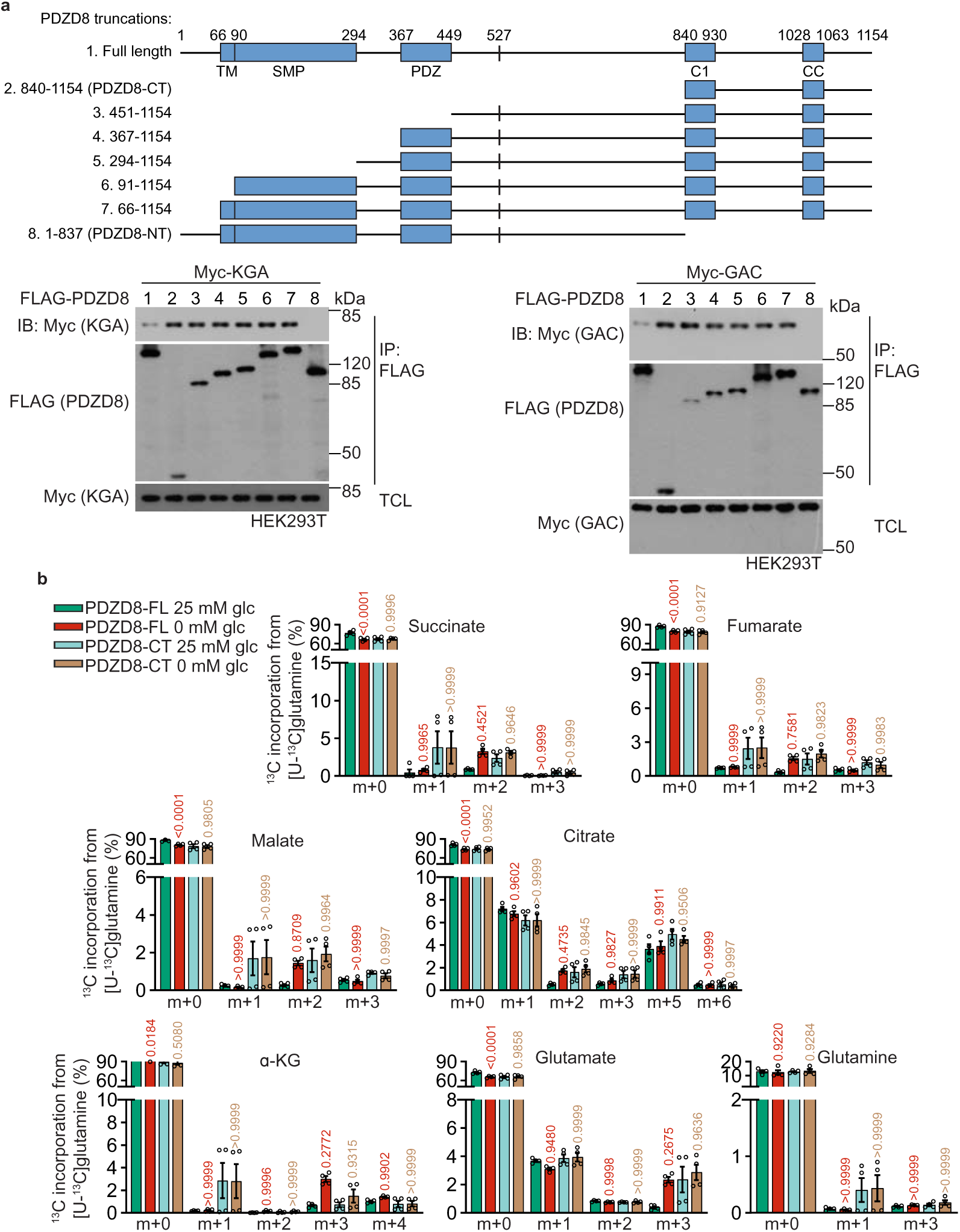
PDZD8-CT dominantly interact with and activates GLS1. **a**, Domain mapping for the region on PDZD8 responsible for interacting with GLS1. Myc-tagged KGA (left panel) or GAC (right panel) was co-transfected with FLAG-tagged PDZD8, or deletion mutants into HEK293T cells. Immunoprecipitation was performed using antibody against FLAG-tag, followed by immunoblotting. **b**, Levels of other isotopomers of the labelled TCA cycle intermediates shown in Fig. 4c. Data are shown as mean ± s.e.m.; *n* = 4 biological replicates for each condition; and *P* values were determined by two-way ANOVA, followed by Tukey. Experiments in this figure were performed three times.

**Extended Data Fig. 8.**
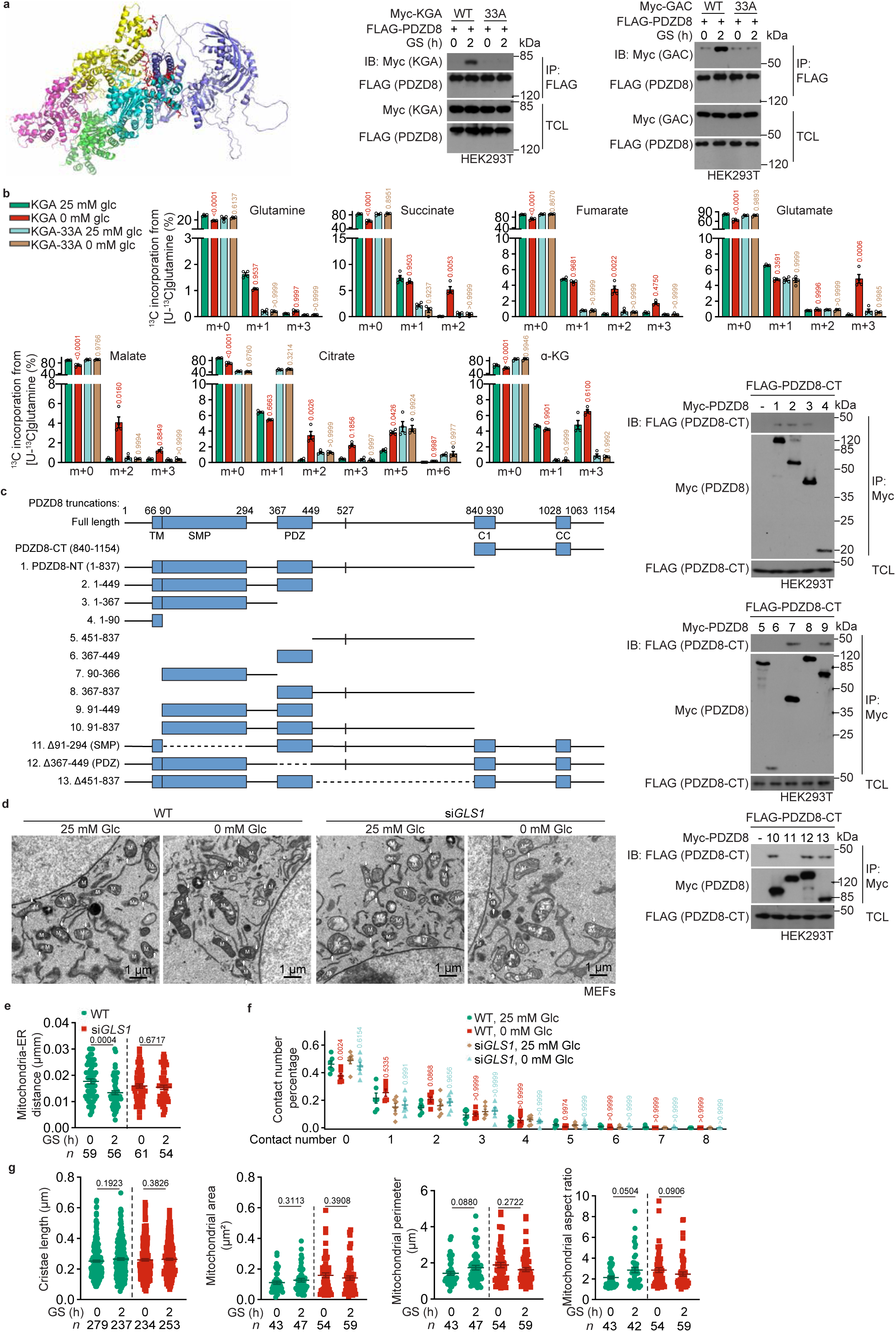
PDZD8 promotes GLS1 activity through interacting with GLS1. **a**, In silico modelling of PDZD8 (blue) bound to GAC (as a tetramer, coloured in magenta, yellow, green and cyan each). The interface is shown as stick structures, and is coloured in red. See detailed list of the 33 residues of GLS1 involved in the interface in “Determination of GLS1-PDZD8 interface” in Methods section. See also left panel for the validation of GLS1-PDZD8 interface, in which HEK293T cells were transfected with FLAG-tagged PDZD8 and Myc-tagged KGA or GAC. Immunoprecipitation was then performed using anti-FLAG antibody, followed by immunoblotting. **b**, Levels of other isotopomers of the labelled TCA cycle intermediates shown in Fig. 4g. Data are shown as mean ± s.e.m.; *n* = 4 for each condition; and *P* values were determined by two-way ANOVA, followed by Tukey, all compared to the unstarved group of each genotype. **c**, PDZD8-NT interacts with PDZD8-CT. Various Myc-tagged PDZD8 deletion mutants were co-transfected with FLAG-tagged PDZD8-CT into HEK293T cells. Immunoprecipitation was performed using anti-Myc antibody, followed by immunoblotting. **d**, Representative TEM images of Fig. 4k. **e**-**g**, Statistical analysis of TEM images in **d**. Data were analysed as in Extended Data Fig. 1b-d, and are shown as mean ± s.e.m.; *n* = 6 (**f**) cells, or labelled on each panel indicates contact numbers (**e**) or mitochondria numbers (**g**, except the leftmost panel cristae) for each condition; and *P* values were determined by two-way ANOVA, followed by Sidak (**f**), by unpaired two-tailed Student’s *t*-test (**e**, WT cells), or by Mann-Whitney test (others). Experiments in this figure were performed three times.

**Extended Data Fig. 9.**
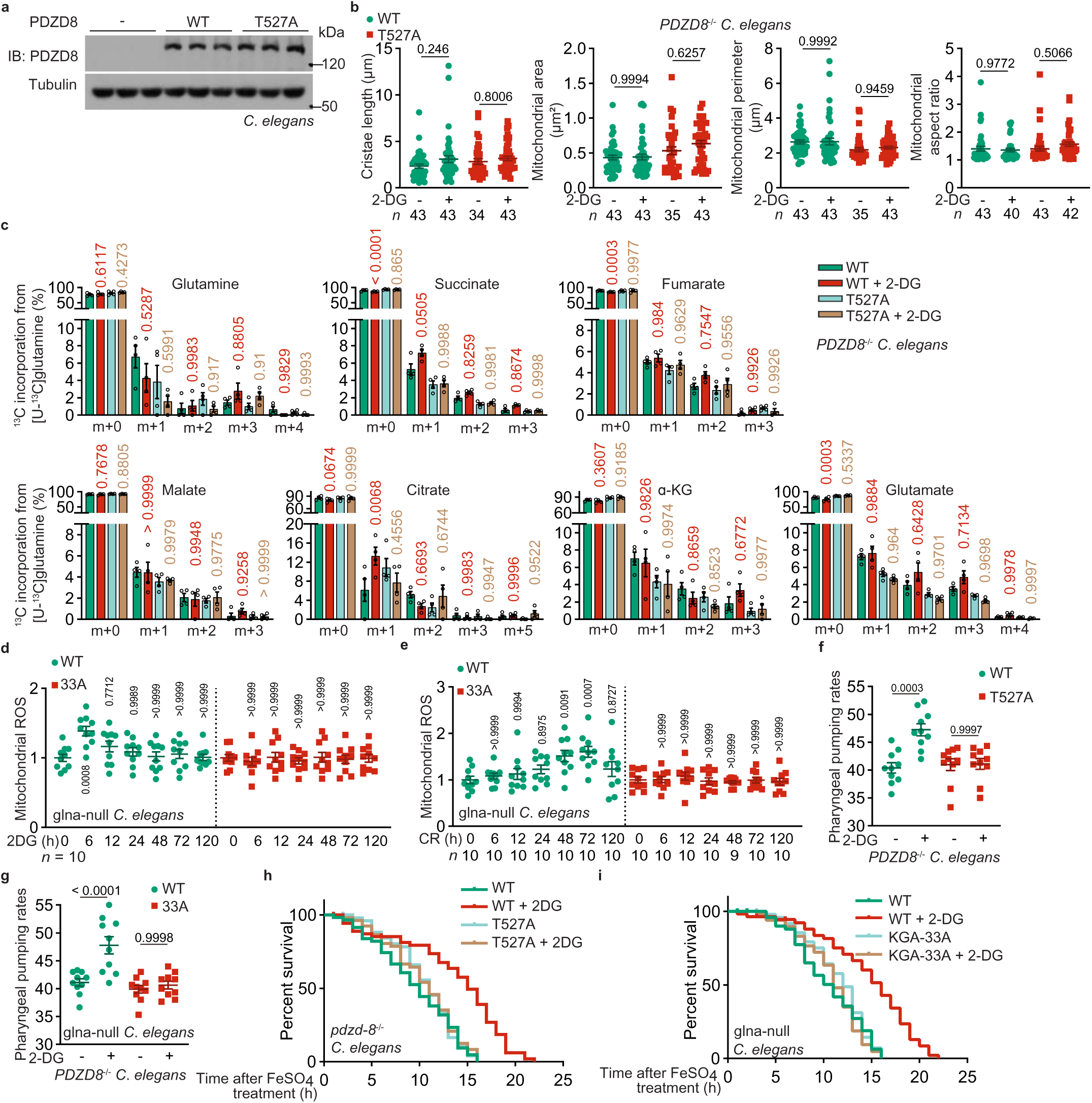
PDZD8 exerts rejuvenating effects in nematodes. **a**, Validation of *pdzd-8*^−/−^ nematodes re-introduced with PDZD8 or its 527A mutant. **b**, Diameter and area of mitochondria in *pdzd-8*^−/−^ nematodes with PDZD8 or its 527A mutant re-introduced. Data are shown as mean ± s.d.; with *n* labelling on the panel. See also Fig. 5a for the ratios of contact length:mitochondrial perimeter. *P* values were determined by two-way ANOVA, followed by Tukey. **c**, Levels of other isotopomers of the labelled TCA cycle intermediates shown in Fig. 5b. Data are shown as mean ± s.d.; *n* = 4 biological replicates for each condition; *P* values were calculated by two-way ANOVA, followed by Tukey, all compared to the unstarved group of each genotype. **d**, **e**, AMPK-PDZD8 axis induces transient mitochondrial ROS in nematodes. Experiments were performed as in Fig. 5h (**d**) and 5m (**e**), respectively, except that the glna-depletion strain re-introduced with KAG-33A was used. Data are shown as mean ± s.d.; *n* values labelled on each panel. *P* values were calculated by two-way ANOVA, followed by Tukey, all compared to the untreated group of each genotype. **f**, **g**, AMPK-PDZD8 axis promotes pharyngeal pumping rates in nematodes. Experiments were performed as in Fig. 5n (**f**) and 5o (**g**), respectively, except that nematodes were treated with 2-DG for 2 days. Data are shown as mean ± s.d.; *n* = 10 for each condition. *P* values were calculated by two-way ANOVA, followed by Tukey, all compared to the untreated group of each genotype. **h**, **i**, AMPK-PDZD8 axis promotes resistance of nematodes to oxidative stress. Experiments were performed as in Fig. 5p (**h**) and 5r (**i**), respectively, except that nematodes were treated with 2-DG for 2 days before the FeSO_4_ treatment. Experiments in this figure were performed three times.

**Extended Data Fig. 10.**
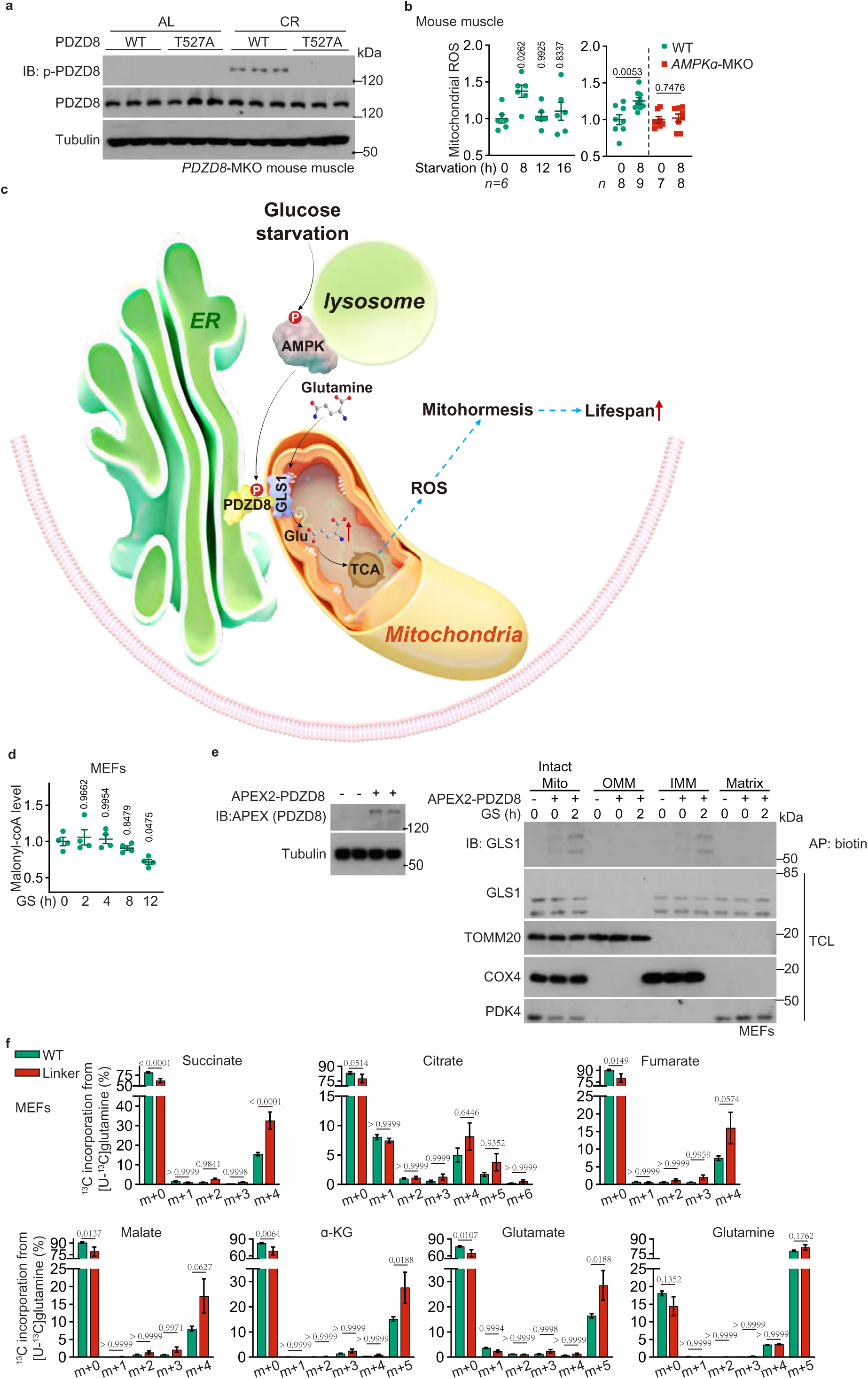
PDZD8 exerts rejuvenating effects in mice. **a**, Validation of PDZD8 phosphorylation in the *PDZD8*-MKO mice with muscle-specific re-introduction of wildtype PDZD8 or PDZD8-T527A. Mice at 8-month-old were subjected to CR for another 3 months, followed by immunoblotting to determine PDZD8-phosphorylation in muscle tissues at 4 p.m. (1 h before the feeding time of each day during CR). **b**, AMPK-PDZD8 axis induces transient mitochondrial ROS in mouse muscle. Wildtype mice and *AMPK*α-MKO mice were starved for desired durations, followed by determination of muscle mitochondrial ROS using the mitoSOX dye. Data are shown as mean ± s.e.m.; *n* (labelled on each panel) values indicate biological replicates for each condition; and *P* values were determined by one-way ANOVA, followed by Tukey (left panel), or unpaired two-tailed Student’s *t*-test (right panel). **c**, Schematic diagramme showing that AMPK-PDZD8 plays a crucial role in the shift of carbon utilisation from glucose to glutamine. In low glucose, the ER-localised PDZD8 is phosphorylated at T527 by AMPK activated via the glucose sensing pathway, which leads to the release of intramolecular autoinhibition (NT towards CT) of PDZD8. As a result, PDZD8 (CT) interacts with and activates the mitochondrial GLS1, promotes glutaminolysis, and also strengthens the ER-mitochondria contact. The promoted glutaminolysis elicits a burst of mitochondrial ROS, which levels off soon owing to the induction of anti-oxidative enzymes (conforming to the characteristics of mitohormesis), thereby executing the anti-ageing effects of starvation and calorie restriction. **d**, Levels of malonyl-CoA in MEFs decreases only after prolonged glucose starvation. MEFs were glucose-starved desired time, followed by determining the levels of malonyl-CoA through HPLC-MS. Data are shown as mean ± s.e.m.; *n* = 4 for each condition; and *P* values were determined by one-way ANOVA, followed by Tukey. **e**, PDZD8 interacts with GLS1 located on the external side of IMM. MEFs stably expressing PDZD8-APEX2 (induced by incubating with doxycycline at a final concentration of 100 ng/ml for 24 h; see validation data in the left panel) were treated with biotinyl tyramide and hydrogen peroxide, followed by purification of OMM, IMM and matrix. The affinity pull-down (AP) of biotinylated proteins was then performed by using Streptavidin Magnetic Beads, followed by immunoblotting. **f**, Forced ER-mitochondria contact formation promotes glutaminolysis in high glucose. Glutaminolysis rates in MEFs with ER-mito linker (the mAKAP1-mRFP-yUBC6 linker) expression (induced by incubating with doxycycline for 12 h) were determined as in Fig. 2a. Data are shown as mean ± s.e.m.; *n* = 4 for each condition; and *P* values were determined by two-way ANOVA, followed by Sidak. Experiments in this figure were performed three times.

## References

1 Cahill, G. F., Jr. Fuel metabolism in starvation. Annu Rev Nutr 26, 1–22, doi:10.1146/annurev.nutr.26.061505.111258 (2006).

2 Gonzalez, A., Hall, M. N., Lin, S. C. & Hardie, D. G. AMPK and TOR: The Yin and Yang of Cellular Nutrient Sensing and Growth Control. Cell metabolism 31, 472–492, doi:10.1016/j.cmet.2020.01.015 (2020).

3 Schoolwerth, A. C., Nazar, B. L. & LaNoue, K. F. Glutamate dehydrogenase activation and ammonia formation by rat kidney mitochondria. The Journal of biological chemistry 253, 6177–6183 (1978).

4 Schoolwerth, A. C. & LaNoue, K. F. The role of microcompartmentation in the regulation of glutamate metabolism by rat kidney mitochondria. The Journal of biological chemistry 255, 3403–3411 (1980).

5 Rej, R. Measurement of aminotransferases: Part 1. Aspartate aminotransferase. Crit Rev Clin Lab Sci 21, 99–186, doi:10.3109/10408368409167137 (1984).

6 Cahill, G. F., Jr., et al. Hormone-fuel interrelationships during fasting. J Clin Invest 45, 1751–1769, doi:10.1172/JCI105481 (1966).

7 Owen, O. E. et al. Brain metabolism during fasting. J Clin Invest 46, 1589–1595, doi:10.1172/JCI105650 (1967).

8 Owen, O. E., Felig, P., Morgan, A. P., Wahren, J. & Cahill, G. F., Jr. Liver and kidney metabolism during prolonged starvation. J Clin Invest 48, 574–583, doi:10.1172/JCI106016 (1969).

9 Moir, A. M. & Zammit, V. A. Monitoring of changes in hepatic fatty acid and glycerolipid metabolism during the starved-to-fed transition in vivo. Studies on awake, unrestrained rats. The Biochemical journal 289 (Pt 1), 49–55, doi:10.1042/bj2890049 (1993).

10 Hui, S. et al. Glucose feeds the TCA cycle via circulating lactate. Nature 551, 115–118, doi:10.1038/nature24057 (2017).

11 Liu, S., Dai, Z., Cooper, D. E., Kirsch, D. G. & Locasale, J. W. Quantitative Analysis of the Physiological Contributions of Glucose to the TCA Cycle. Cell metabolism 32, 619–628 e621, doi:10.1016/j.cmet.2020.09.005 (2020).

12 Hui, S. et al. Quantitative Fluxomics of Circulating Metabolites. Cell metabolism 32, 676–688 e674, doi:10.1016/j.cmet.2020.07.013 (2020).

13 Leloir, L. F. & Munoz, J. M. Fatty acid oxidation in liver. The Biochemical journal 33, 734–746, doi:10.1042/bj0330734 (1939).

14 Grafflin, A. L. & Green, D. E. Studies on the cyclophorase system; the complete oxidation of fatty acids. The Journal of biological chemistry 176, 95–115 (1948).

15 Skrede, S. & Bremer, J. The compartmentation of CoA and fatty acid activating enzymes in rat liver mitochondria. Eur J Biochem 14, 465–472, doi:10.1111/j.1432-1033.1970.tb00312.x (1970).

16 Knoop, F. Der Abbau aromatischer fettsäuren im tierkörper, Verlag nicht ermittelbar, (1904).

17 Krebs, H. A. & Johnson, W. A. Metabolism of ketonic acids in animal tissues. The Biochemical journal 31, 645–660, doi:10.1042/bj0310645 (1937).

18 Novelli, G. D. & Lipmann, F. The Catalytic Function of Coenzyme a in Citric Acid Synthesis. Journal of Biological Chemistry 182, 213–228, doi:10.1016/s0021-9258(18)56541-8 (1950).

19 Bergstrom, J., Furst, P., Noree, L. O. & Vinnars, E. Intracellular free amino acid concentration in human muscle tissue. J Appl Physiol 36, 693–697, doi:10.1152/jappl.1974.36.6.693 (1974).

20 Stumvoll, M., Perriello, G., Meyer, C. & Gerich, J. Role of glutamine in human carbohydrate metabolism in kidney and other tissues. Kidney Int 55, 778–792, doi:10.1046/j.1523-1755.1999.055003778.x (1999).

21 Perry, R. J. et al. Leptin Mediates a Glucose-Fatty Acid Cycle to Maintain Glucose Homeostasis in Starvation. Cell 172, 234–248 e217, doi:10.1016/j.cell.2017.12.001 (2018).

22 Ruderman, N. B. & Berger, M. The formation of glutamine and alanine in skeletal muscle. The Journal of biological chemistry 249, 5500–5506 (1974).

23 Marliss, E. B., Aoki, T. T., Pozefsky, T., Most, A. S. & Cahill, G. F., Jr. Muscle and splanchnic glutmine and glutamate metabolism in postabsorptive andstarved man. J Clin Invest 50, 814–817, doi:10.1172/JCI106552 (1971).

24 Aikawa, T., Matsutaka, H., Yamamoto, H., Okuda, T. & Ishikawa, E. Gluconeogenesis and amino acid metabolism. II. Inter-organal relations and roles of glutamine and alanine in the amino acid metabolism of fasted rats. J Biochem 74, 1003–1017 (1973).

25 Moreadith, R. W. & Lehninger, A. L. The pathways of glutamate and glutamine oxidation by tumor cell mitochondria. Role of mitochondrial NAD(P)+-dependent malic enzyme. The Journal of biological chemistry 259, 6215–6221 (1984).

26 Ross, B. D., Hems, R. & Krebs, H. A. The rate of gluconeogenesis from various precursors in the perfused rat liver. The Biochemical journal 102, 942–951, doi:10.1042/bj1020942 (1967).

27 Consoli, A., Kennedy, F., Miles, J. & Gerich, J. Determination of Krebs cycle metabolic carbon exchange in vivo and its use to estimate the individual contributions of gluconeogenesis and glycogenolysis to overall glucose output in man. J Clin Invest 80, 1303–1310, doi:10.1172/JCI113206 (1987).

28 Nurjhan, N. et al. Glutamine: a major gluconeogenic precursor and vehicle for interorgan carbon transport in man. J Clin Invest 95, 272–277, doi:10.1172/JCI117651 (1995).

29 Stumvoll, M. et al. Uptake and release of glucose by the human kidney. Postabsorptive rates and responses to epinephrine. J Clin Invest 96, 2528–2533, doi:10.1172/JCI118314 (1995).

30 Hankard, R. G., Haymond, M. W. & Darmaun, D. Role of glutamine as a glucose precursor in fasting humans. Diabetes 46, 1535–1541, doi:10.2337/diacare.46.10.1535 (1997).

31 Elwyn, D. H., Parikh, H. C. & Shoemaker, W. C. Amino acid movements between gut, liver, and periphery in unanesthetized dogs. Am J Physiol 215, 1260–1275, doi:10.1152/ajplegacy.1968.215.5.1260 (1968).

32 Welbourne, T. C. Ammonia production and glutamine incorporation into glutathione in the functioning rat kidney. Can J Biochem 57, 233–237, doi:10.1139/o79-029 (1979).

33 Sato, H. et al. Redox imbalance in cystine/glutamate transporter-deficient mice. The Journal of biological chemistry 280, 37423–37429, doi:10.1074/jbc.M506439200 (2005).

34 DeBerardinis, R. J. et al. Beyond aerobic glycolysis: transformed cells can engage in glutamine metabolism that exceeds the requirement for protein and nucleotide synthesis. Proceedings of the National Academy of Sciences of the United States of America 104, 19345–19350, doi:10.1073/pnas.0709747104 (2007).

35 Mullen, A. R. et al. Reductive carboxylation supports growth in tumour cells with defective mitochondria. Nature 481, 385–388, doi:10.1038/nature10642 (2011).

36 Son, J. et al. Glutamine supports pancreatic cancer growth through a KRAS-regulated metabolic pathway. Nature 496, 101–105, doi:10.1038/nature12040 (2013).

37 Carling, D., Zammit, V. A. & Hardie, D. G. A common bicyclic protein kinase cascade inactivates the regulatory enzymes of fatty acid and cholesterol biosynthesis. FEBS letters 223, 217–222, doi:10.1016/0014-5793(87)80292-2 (1987).

38 Xiao, B. et al. Structure of mammalian AMPK and its regulation by ADP. Nature 472, 230–233, doi:nature09932 [pii] 10.1038/nature09932 (2011).

39 Salt, I. P., Johnson, G., Ashcroft, S. J. H. & Hardie, D. G. AMP-activated protein kinase is activated by low glucose in cell lines derived from pancreatic b cells, and may regulate insulin release. Biochem. J. 335, 533–539 (1998).

40 Zhang, C. S. et al. Fructose-1,6-bisphosphate and aldolase mediate glucose sensing by AMPK. Nature 548, 112–116, doi:10.1038/nature23275 (2017).

41 Zhang, C. S. et al. The lysosomal v-ATPase-Ragulator complex Is a common activator for AMPK and mTORC1, acting as a switch between catabolism and anabolism. Cell Metab. 20, 526–540, doi:10.1016/j.cmet.2014.06.014 (2014).

42 Li, M. et al. Transient Receptor Potential V Channels Are Essential for Glucose Sensing by Aldolase and AMPK. Cell metabolism 30, 508–524 e512, doi:10.1016/j.cmet.2019.05.018 (2019).

43 Zhang, Y. L. et al. AMP as a low-energy charge signal autonomously initiates assembly of AXIN-AMPK-LKB1 complex for AMPK activation. Cell Metab. 18, 546–555, doi:10.1016/j.cmet.2013.09.005 (2013).

44 Davies, S. P., Sim, A. T. & Hardie, D. G. Location and function of three sites phosphorylated on rat acetyl-CoA carboxylase by the AMP-activated protein kinase. Eur J Biochem 187, 183–190 (1990).

45 McGarry, J. D., Mannaerts, G. P. & Foster, D. W. A possible role for malonyl-CoA in the regulation of hepatic fatty acid oxidation and ketogenesis. J Clin Invest 60, 265–270, doi:10.1172/JCI108764 (1977).

46 Krause, U., Bertrand, L. & Hue, L. Control of p70 ribosomal protein S6 kinase and acetyl-CoA carboxylase by AMP-activated protein kinase and protein phosphatases in isolated hepatocytes. Eur J Biochem 269, 3751–3759, doi:10.1046/j.1432-1033.2002.03074.x (2002).

47 Inoki, K., Zhu, T. & Guan, K. L. TSC2 mediates cellular energy response to control cell growth and survival. Cell 115, 577–590 (2003).

48 Horman, S. et al. Activation of AMP-activated protein kinase leads to the phosphorylation of elongation factor 2 and an inhibition of protein synthesis. Current biology : CB 12, 1419–1423, doi:10.1016/s0960-9822(02)01077-1 (2002).

49 Meley, D. et al. AMP-activated protein kinase and the regulation of autophagic proteolysis. The Journal of biological chemistry 281, 34870–34879, doi:10.1074/jbc.M605488200 (2006).

50 Egan, D. F. et al. Phosphorylation of ULK1 (hATG1) by AMP-activated protein kinase connects energy sensing to mitophagy. Science 331, 456–461, doi:10.1126/science.1196371 (2011).

51 Kim, J., Kundu, M., Viollet, B. & Guan, K. L. AMPK and mTOR regulate autophagy through direct phosphorylation of Ulk1. Nature cell biology 13, 132–141, doi:10.1038/ncb2152 (2011).

52 Nakashima, K. & Yakabe, Y. AMPK activation stimulates myofibrillar protein degradation and expression of atrophy-related ubiquitin ligases by increasing FOXO transcription factors in C2C12 myotubes. Biosci Biotechnol Biochem 71, 1650–1656, doi:10.1271/bbb.70057 (2007).

53 Zong, Y. et al. Hierarchical activation of compartmentalized pools of AMPK depends on severity of nutrient or energy stress. Cell Res 29, 460–473, doi:10.1038/s41422-019-0163-6 (2019).

54 Weekes, J., Ball, K. L., Caudwell, F. B. & Hardie, D. G. Specificity determinants for the AMP-activated protein kinase and its plant homologue analysed using synthetic peptides. FEBS letters 334, 335–339, doi:10.1016/0014-5793(93)80706-z (1993).

55 Dale, S., Wilson, W. A., Edelman, A. M. & Hardie, D. G. Similar substrate recognition motifs for mammalian AMP-activated protein kinase, higher plant HMG-CoA reductase kinase-A, yeast SNF1, and mammalian calmodulin-dependent protein kinase I. FEBS letters 361, 191–195 (1995).

56 Scott, J. W., Norman, D. G., Hawley, S. A., Kontogiannis, L. & Hardie, D. G. Protein kinase substrate recognition studied using the recombinant catalytic domain of AMP-activated protein kinase and a model substrate. J Mol Biol 317, 309–323, doi:10.1006/jmbi.2001.5316 (2002).

57 Gwinn, D. M. et al. AMPK phosphorylation of raptor mediates a metabolic checkpoint. Mol. Cell 30, 214–226 (2008).

58 Johnson, J. L. et al. An atlas of substrate specificities for the human serine/threonine kinome. Nature, doi:10.1038/s41586-022-05575-3 (2023).

59 Cai, Z. et al. Phosphorylation of PDHA by AMPK Drives TCA Cycle to Promote Cancer Metastasis. Molecular cell 80, 263–278 e267, doi:10.1016/j.molcel.2020.09.018 (2020).

60 Hirabayashi, Y. et al. ER-mitochondria tethering by PDZD8 regulates Ca(2+) dynamics in mammalian neurons. Science 358, 623–630, doi:10.1126/science.aan6009 (2017).

61 Guillen-Samander, A., Bian, X. & De Camilli, P. PDZD8 mediates a Rab7-dependent interaction of the ER with late endosomes and lysosomes. Proceedings of the National Academy of Sciences of the United States of America 116, 22619–22623, doi:10.1073/pnas.1913509116 (2019).

62 De Vos, K. J. et al. VAPB interacts with the mitochondrial protein PTPIP51 to regulate calcium homeostasis. Hum Mol Genet 21, 1299–1311, doi:10.1093/hmg/ddr559 (2012).

63 Stoica, R. et al. ER-mitochondria associations are regulated by the VAPB-PTPIP51 interaction and are disrupted by ALS/FTD-associated TDP-43. Nat Commun 5, 3996, doi:10.1038/ncomms4996 (2014).

64 Sakamoto, K. et al. Deficiency of LKB1 in skeletal muscle prevents AMPK activation and glucose uptake during contraction. The EMBO journal 24, 1810–1820, doi:10.1038/sj.emboj.7600667 (2005).

65 Jorgensen, S. B. et al. Effects of alpha-AMPK knockout on exercise-induced gene activation in mouse skeletal muscle. FASEB journal : official publication of the Federation of American Societies for Experimental Biology 19, 1146–1148, doi:10.1096/fj.04-3144fje (2005).

66 McGarry, J. D. & Foster, D. W. Regulation of hepatic fatty acid oxidation and ketone body production. Annu Rev Biochem 49, 395–420, doi:10.1146/annurev.bi.49.070180.002143 (1980).

67 Perry, R. J., Peng, L., Cline, G. W., Petersen, K. F. & Shulman, G. I. A Non-invasive Method to Assess Hepatic Acetyl-CoA In Vivo. Cell metabolism 25, 749–756, doi:10.1016/j.cmet.2016.12.017 (2017).

68 Robinson, M. M. et al. Novel mechanism of inhibition of rat kidney-type glutaminase by bis-2-(5-phenylacetamido-1,2,4-thiadiazol-2-yl)ethyl sulfide (BPTES). The Biochemical journal 406, 407–414, doi:10.1042/BJ20070039 (2007).

69 Wolf, H. P., Eistetter, K. & Ludwig, G. Phenylalkyloxirane carboxylic acids, a new class of hypoglycaemic substances: hypoglycaemic and hypoketonaemic effects of sodium 2-[5-(4-chlorophenyl)-pentyl]-oxirane-2-carboxylate (B 807-27) in fasted animals. Diabetologia 22, 456–463, doi:10.1007/BF00282590 (1982).

70 Ferreira, A. P. et al. Active glutaminase C self-assembles into a supratetrameric oligomer that can be disrupted by an allosteric inhibitor. The Journal of biological chemistry 288, 28009–28020, doi:10.1074/jbc.M113.501346 (2013).

71 Errera, M. & Greenstein, J. P. Phosphate-activated glutaminase in kidney and other tissues. The Journal of biological chemistry 178, 495–502 (1949).

72 Cassago, A. et al. Mitochondrial localization and structure-based phosphate activation mechanism of Glutaminase C with implications for cancer metabolism. Proceedings of the National Academy of Sciences of the United States of America 109, 1092–1097, doi:10.1073/pnas.1112495109 (2012).

73 Haussinger, D. et al. Role of plasma membrane transport in hepatic glutamine metabolism. Eur J Biochem 152, 597–603, doi:10.1111/j.1432-1033.1985.tb09237.x (1985).

74 Lenzen, C., Soboll, S., Sies, H. & Haussinger, D. pH control of hepatic glutamine degradation. Role of transport. Eur J Biochem 166, 483–488, doi:10.1111/j.1432-1033.1987.tb13541.x (1987).

75 Chen, W. W., Freinkman, E., Wang, T., Birsoy, K. & Sabatini, D. M. Absolute Quantification of Matrix Metabolites Reveals the Dynamics of Mitochondrial Metabolism. Cell 166, 1324–1337 e1311, doi:10.1016/j.cell.2016.07.040 (2016).

76 Crompton, M., McGivan, J. D. & Chappell, J. B. The intramitochondrial location of the glutaminase isoenzymes of pig kidney. The Biochemical journal 132, 27–34, doi:10.1042/bj1320027 (1973).

77 Schulz, T. J. et al. Glucose restriction extends Caenorhabditis elegans life span by inducing mitochondrial respiration and increasing oxidative stress. Cell metabolism 6, 280–293, doi:10.1016/j.cmet.2007.08.011 (2007).

78 Apfeld, J., O’Connor, G., McDonagh, T., Distefano, P. S. & Curtis, R. The AMP-activated protein kinase AAK-2 links energy levels and insulin-like signals to lifespan in C. elegans. Genes Dev 18, 3004–3009 (2004).

79 Zarse, K. et al. Impaired insulin/IGF1 signaling extends life span by promoting mitochondrial L-proline catabolism to induce a transient ROS signal. Cell metabolism 15, 451–465, doi:10.1016/j.cmet.2012.02.013 (2012).

80 Greer, E. L. et al. An AMPK-FOXO pathway mediates longevity induced by a novel method of dietary restriction in C. elegans. Current biology : CB 17, 1646–1656, doi:10.1016/j.cub.2007.08.047 (2007).

81 Greer, E. L. & Brunet, A. Different dietary restriction regimens extend lifespan by both independent and overlapping genetic pathways in C. elegans. Aging cell 8, 113–127, doi:10.1111/j.1474-9726.2009.00459.x (2009).

82 Cox, C. S. et al. Mitohormesis in Mice via Sustained Basal Activation of Mitochondrial and Antioxidant Signaling. Cell metabolism 28, 776–786 e775, doi:10.1016/j.cmet.2018.07.011 (2018).

83 Crabtree, H. G. Observations on the carbohydrate metabolism of tumours. The Biochemical journal 23, 536–545, doi:10.1042/bj0230536 (1929).

84 Maggs, D. G. et al. Interstitial fluid concentrations of glycerol, glucose, and amino acids in human quadricep muscle and adipose tissue. Evidence for significant lipolysis in skeletal muscle. J Clin Invest 96, 370–377, doi:10.1172/JCI118043 (1995).

85 Sjostrand, M. et al. Measurements of interstitial muscle glycerol in normal and insulin-resistant subjects. J Clin Endocrinol Metab 87, 2206–2211, doi:10.1210/jcem.87.5.8495 (2002).

86 Landau, B. R. et al. Glycerol production and utilization in humans: sites and quantitation. Am J Physiol 271, E1110–1117, doi:10.1152/ajpendo.1996.271.6.E1110 (1996).

87 Kvamme, E., Torgner, I. A. & Roberg, B. Evidence indicating that pig renal phosphate-activated glutaminase has a functionally predominant external localization in the inner mitochondrial membrane. The Journal of biological chemistry 266, 13185–13192 (1991).

88 Roberg, B., Torgner, I. A. & Kvamme, E. The orientation of phosphate activated glutaminase in the inner mitochondrial membrane of synaptic and non-synaptic rat brain mitochondria. Neurochem Int 27, 367–376, doi:10.1016/0197-0186(95)00018-4 (1995).

89 Roberg, B. et al. Properties and submitochondrial localization of pig and rat renal phosphate-activated glutaminase. American journal of physiology. Cell physiology 279, C648–657, doi:10.1152/ajpcell.2000.279.3.C648 (2000).

90 Csordas, G. et al. Structural and functional features and significance of the physical linkage between ER and mitochondria. The Journal of cell biology 174, 915–921, doi:10.1083/jcb.200604016 (2006).

## References for Methods section

91 Demeter-Haludka, V. et al. Examination of the Role of Mitochondrial Morphology and Function in the Cardioprotective Effect of Sodium Nitrite Administered 24 h Before Ischemia/Reperfusion Injury. Front Pharmacol 9, 286, doi:10.3389/fphar.2018.00286 (2018).

92 Thoudam, T. et al. PDK4 Augments ER-Mitochondria Contact to Dampen Skeletal Muscle Insulin Signaling During Obesity. Diabetes 68, 571–586, doi:10.2337/db18-0363 (2019).

93 Cali, T. & Brini, M. Quantification of organelle contact sites by split-GFP-based contact site sensors (SPLICS) in living cells. Nature protocols 16, 5287–5308, doi:10.1038/s41596-021-00614-1 (2021).

94 Zhang, C. S. et al. The aldolase inhibitor aldometanib mimics glucose starvation to activate lysosomal AMPK. Nat Metab, doi:10.1038/s42255-022-00640-7 (2022).

95 Perez, C. L. & Van Gilst, M. R. A 13C isotope labeling strategy reveals the influence of insulin signaling on lipogenesis in C. elegans. Cell metabolism 8, 266–274, doi:10.1016/j.cmet.2008.08.007 (2008).

96 Falk, M. J. et al. Stable isotopic profiling of intermediary metabolic flux in developing and adult stage Caenorhabditis elegans. J Vis Exp, doi:10.3791/2288 (2011).

97 Vergano, S. S. et al. In vivo metabolic flux profiling with stable isotopes discriminates sites and quantifies effects of mitochondrial dysfunction in C. elegans. Mol Genet Metab 111, 331–341, doi:10.1016/j.ymgme.2013.12.011 (2014).

98 Schuh, R. A., Jackson, K. C., Khairallah, R. J., Ward, C. W. & Spangenburg, E. E. Measuring mitochondrial respiration in intact single muscle fibers. Am J Physiol Regul Integr Comp Physiol 302, R712–719, doi:10.1152/ajpregu.00229.2011 (2012).

99 Koopman, M. et al. A screening-based platform for the assessment of cellular respiration in Caenorhabditis elegans. Nature protocols 11, 1798–1816, doi:10.1038/nprot.2016.106 (2016).

100 Sarasija, S. & Norman, K. R. Measurement of Oxygen Consumption Rates in Intact Caenorhabditis elegans. J Vis Exp, doi:10.3791/59277 (2019).

101 Ng, L. F. & Gruber, J. Measurement of Respiration Rate in Live Caenorhabditis elegans. Bio Protoc 9, e3243, doi:10.21769/BioProtoc.3243 (2019).

102 Yang, S. H. et al. Estrogen receptor beta as a mitochondrial vulnerability factor. The Journal of biological chemistry 284, 9540–9548, doi:10.1074/jbc.M808246200 (2009).

103 Vaccaro, A. et al. Sleep Loss Can Cause Death through Accumulation of Reactive Oxygen Species in the Gut. Cell 181, 1307–1328 e1315, doi:10.1016/j.cell.2020.04.049 (2020).

104 Dingley, S. et al. Mitochondrial respiratory chain dysfunction variably increases oxidant stress in Caenorhabditis elegans. Mitochondrion 10, 125–136, doi:10.1016/j.mito.2009.11.003 (2010).

105 Yang, W. & Hekimi, S. A mitochondrial superoxide signal triggers increased longevity in Caenorhabditis elegans. PLoS Biol 8, e1000556, doi:10.1371/journal.pbio.1000556 (2010).

106 Curthoys, N. P. & Weiss, R. F. Regulation of renal ammoniagenesis. Subcellular localization of rat kidney glutaminase isoenzymes. The Journal of biological chemistry 249, 3261–3266 (1974).

107 Tong, Y. et al. SUCLA2-coupled regulation of GLS succinylation and activity counteracts oxidative stress in tumor cells. Molecular cell 81, 2303–2316 e2308, doi:10.1016/j.molcel.2021.04.002 (2021).

108 Li, M. et al. Aldolase is a sensor for both low and high glucose, linking to AMPK and mTORC1. Cell Res, doi:10.1038/s41422-020-00456-8 (2020).

109 Kirsch, N. et al. Angiopoietin-like 4 Is a Wnt Signaling Antagonist that Promotes LRP6 Turnover. Developmental cell 43, 71–82 e76, doi:10.1016/j.devcel.2017.09.011 (2017).

110 Benavente, F. et al. Novel C1q receptor-mediated signaling controls neural stem cell behavior and neurorepair. Elife 9, doi:10.7554/eLife.55732 (2020).

111 Espada, L. et al. Loss of metabolic plasticity underlies metformin toxicity in aged Caenorhabditis elegans. Nat Metab 2, 1316–1331, doi:10.1038/s42255-020-00307-1 (2020).

112 Wu, L. et al. An Ancient, Unified Mechanism for Metformin Growth Inhibition in C. elegans and Cancer. Cell 167, 1705–1718 e1713, doi:10.1016/j.cell.2016.11.055 (2016).

113 Martin-Montalvo, A. et al. Metformin improves healthspan and lifespan in mice. Nat Commun 4, 2192, doi:10.1038/ncomms3192 (2013).

114 De Rosa, M. J. et al. The flight response impairs cytoprotective mechanisms by activating the insulin pathway. Nature 573, 135–138, doi:10.1038/s41586-019-1524-5 (2019).

115 Yuan, J. et al. Two conserved epigenetic regulators prevent healthy ageing. Nature 579, 118–122, doi:10.1038/s41586-020-2037-y (2020).

116 Zhang, H. et al. NAD(+) repletion improves mitochondrial and stem cell function and enhances life span in mice. Science 352, 1436–1443, doi:10.1126/science.aaf2693 (2016).

117 Chu, V. T. et al. Efficient generation of Rosa26 knock-in mice using CRISPR/Cas9 in C57BL/6 zygotes. BMC Biotechnol 16, 4, doi:10.1186/s12896-016-0234-4 (2016).

118 Xiao, C. et al. Lymphoproliferative disease and autoimmunity in mice with increased miR-17-92 expression in lymphocytes. Nat Immunol 9, 405–414, doi:10.1038/ni1575 (2008).

119 Mali, P. et al. RNA-guided human genome engineering via Cas9. Science 339, 823–826, doi:10.1126/science.1232033 (2013).

120 Taft, R. In Vitro Fertilization in Mice. Cold Spring Harbor protocols 2017, pdb prot094508, doi:10.1101/pdb.prot094508 (2017).

121 Park, S. J. et al. DNA-PK Promotes the Mitochondrial, Metabolic, and Physical Decline that Occurs During Aging. Cell metabolism 26, 447, doi:10.1016/j.cmet.2017.07.005 (2017).

122 Mair, W. et al. Lifespan extension induced by AMPK and calcineurin is mediated by CRTC-1 and CREB. Nature 470, 404–408, doi:10.1038/nature09706 (2011).

123 Ma, T. et al. Low-dose metformin targets the lysosomal AMPK pathway through PEN2. Nature, doi:10.1038/s41586-022-04431-8 (2022).

124 Han, M. et al. A Systematic RNAi Screen Reveals a Novel Role of a Spindle Assembly Checkpoint Protein BuGZ in Synaptic Transmission in C. elegans. Front Mol Neurosci 10, 141, doi:10.3389/fnmol.2017.00141 (2017).

125 Fang, E. F. et al. NAD(+) Replenishment Improves Lifespan and Healthspan in Ataxia Telangiectasia Models via Mitophagy and DNA Repair. Cell metabolism 24, 566–581, doi:10.1016/j.cmet.2016.09.004 (2016).

126 Bolger, A. M., Lohse, M. & Usadel, B. Trimmomatic: a flexible trimmer for Illumina sequence data. Bioinformatics 30, 2114–2120, doi:10.1093/bioinformatics/btu170 (2014).

127 Dobin, A. et al. STAR: ultrafast universal RNA-seq aligner. Bioinformatics 29, 15–21, doi:10.1093/bioinformatics/bts635 (2013).

128 Liao, Y., Smyth, G. K. & Shi, W. featureCounts: an efficient general purpose program for assigning sequence reads to genomic features. Bioinformatics 30, 923–930, doi:10.1093/bioinformatics/btt656 (2014).

129 Fang, R., Jiang, Q., Jia, X. & Jiang, Z. ARMH3-mediated recruitment of PI4KB directs Golgi-to-endosome trafficking and activation of the antiviral effector STING. Immunity 56, 500–515 e506, doi:10.1016/j.immuni.2023.02.004 (2023).

130 Lin, S. C. & Morrison-Bogorad, M. Cloning and characterization of a testis-specific thymosin beta 10 cDNA. Expression in post-meiotic male germ cells. The Journal of biological chemistry 266, 23347–23353 (1991).

131 Li, M. et al. Hierarchical inhibition of mTORC1 by glucose starvation-triggered AXIN lysosomal translocation and by AMPK. Life Metabolism, doi:10.1093/lifemeta/load005 (2023).

132 Cai, W. F. et al. Glutaminase GLS1 senses glutamine availability in a non-enzymatic manner triggering mitochondrial fusion. Cell Res 28, 865–867, doi:10.1038/s41422-018-0057-z (2018).

133 Jang, W. et al. Endosomal lipid signaling reshapes the endoplasmic reticulum to control mitochondrial function. Science 378, eabq5209, doi:10.1126/science.abq5209 (2022).

134 Vacanti, N. M. et al. Regulation of substrate utilization by the mitochondrial pyruvate carrier. Molecular cell 56, 425–435, doi:10.1016/j.molcel.2014.09.024 (2014).

135 Millard, P., Letisse, F., Sokol, S. & Portais, J. C. IsoCor: correcting MS data in isotope labeling experiments. Bioinformatics 28, 1294–1296, doi:10.1093/bioinformatics/bts127 (2012).

136 Millard, P., Letisse, F., Sokol, S. & Portais, J. C. Correction of MS data for naturally occurring isotopes in isotope labelling experiments. Methods Mol Biol 1191, 197–207, doi:10.1007/978-1-4939-1170-7_12 (2014).

137 Wiechert, W. & de Graaf, A. A. In vivo stationary flux analysis by 13C labeling experiments. Adv Biochem Eng Biotechnol 54, 109–154, doi:10.1007/BFb0102334 (1996).

138 Schell, J. C. et al. A role for the mitochondrial pyruvate carrier as a repressor of the Warburg effect and colon cancer cell growth. Molecular cell 56, 400–413, doi:10.1016/j.molcel.2014.09.026 (2014).

139 Wang, Y. et al. Uncoupling Hepatic Oxidative Phosphorylation Reduces Tumor Growth in Two Murine Models of Colon Cancer. Cell reports 24, 47–55, doi:10.1016/j.celrep.2018.06.008 (2018).

140 Wang, Y. et al. AdipoRon exerts opposing effects on insulin sensitivity via fibroblast growth factor 21-mediated time-dependent mechanisms. The Journal of biological chemistry 298, 101641, doi:10.1016/j.jbc.2022.101641 (2022).

141 TeSlaa, T. et al. The Source of Glycolytic Intermediates in Mammalian Tissues. Cell metabolism 33, 367–378 e365, doi:10.1016/j.cmet.2020.12.020 (2021).

142 Bajad, S. U. et al. Separation and quantitation of water soluble cellular metabolites by hydrophilic interaction chromatography-tandem mass spectrometry. Journal of chromatography. A 1125, 76–88, doi:10.1016/j.chroma.2006.05.019 (2006).

143 Ramirez-Aportela, E., Lopez-Blanco, J. R. & Chacon, P. FRODOCK 2.0: fast protein-protein docking server. Bioinformatics 32, 2386–2388, doi:10.1093/bioinformatics/btw141 (2016).

144 DeLaBarre, B. et al. Full-length human glutaminase in complex with an allosteric inhibitor. Biochemistry 50, 10764–10770, doi:10.1021/bi201613d (2011).

145 Jumper, J. et al. Highly accurate protein structure prediction with AlphaFold. Nature 596, 583–589, doi:10.1038/s41586-021-03819-2 (2021).

146 Varadi, M. et al. AlphaFold Protein Structure Database: massively expanding the structural coverage of protein-sequence space with high-accuracy models. Nucleic Acids Res 50, D439–D444, doi:10.1093/nar/gkab1061 (2022).

147 Preez, G. D. et al. Oxygen consumption rate of Caenorhabditis elegans as a high-throughput endpoint of toxicity testing using the Seahorse XF(e)96 Extracellular Flux Analyzer. Scientific reports 10, 4239, doi:10.1038/s41598-020-61054-7 (2020).

148 Wang, Q. et al. IL-27 signalling promotes adipocyte thermogenesis and energy expenditure. Nature 600, 314–318, doi:10.1038/s41586-021-04127-5 (2021).

149 Kuznetsov, A. V. et al. Analysis of mitochondrial function in situ in permeabilized muscle fibers, tissues and cells. Nature protocols 3, 965–976, doi:10.1038/nprot.2008.61 (2008).

150 Makrecka-Kuka, M., Krumschnabel, G. & Gnaiger, E. High-Resolution Respirometry for Simultaneous Measurement of Oxygen and Hydrogen Peroxide Fluxes in Permeabilized Cells, Tissue Homogenate and Isolated Mitochondria. Biomolecules 5, 1319–1338, doi:10.3390/biom5031319 (2015).

151 Jiang, B. et al. Filamentous GLS1 promotes ROS-induced apoptosis upon glutamine deprivation via insufficient asparagine synthesis. Molecular cell 82, 1821–1835 e1826, doi:10.1016/j.molcel.2022.03.016 (2022).

152 Martell, J. D., Deerinck, T. J., Lam, S. S., Ellisman, M. H. & Ting, A. Y. Electron microscopy using the genetically encoded APEX2 tag in cultured mammalian cells. Nature protocols 12, 1792–1816, doi:10.1038/nprot.2017.065 (2017).

153 Koopman, W. J. et al. Inhibition of complex I of the electron transport chain causes O2-. -mediated mitochondrial outgrowth. American journal of physiology. Cell physiology 288, C1440–1450, doi:10.1152/ajpcell.00607.2004 (2005).

154 De Vos, K. J., Allan, V. J., Grierson, A. J. & Sheetz, M. P. Mitochondrial function and actin regulate dynamin-related protein 1-dependent mitochondrial fission. Current biology : CB 15, 678–683, doi:10.1016/j.cub.2005.02.064 (2005).

155 Filadi, R. et al. Mitofusin 2 ablation increases endoplasmic reticulum-mitochondria coupling. Proceedings of the National Academy of Sciences of the United States of America 112, E2174–2181, doi:10.1073/pnas.1504880112 (2015).

156 Sun, L. et al. Katanin p60-like 1 sculpts the cytoskeleton in mechanosensory cilia. The Journal of cell biology 220, doi:10.1083/jcb.202004184 (2021).

157 Du, W. et al. Kinesin 1 Drives Autolysosome Tubulation. Developmental cell 37, 326–336, doi:10.1016/j.devcel.2016.04.014 (2016).

158 Chen, X. et al. Mosaic composition of RIP1-RIP3 signalling hub and its role in regulating cell death. Nature cell biology 24, 471–482, doi:10.1038/s41556-022-00854-7 (2022).

159 Dempsey, G. T., Vaughan, J. C., Chen, K. H., Bates, M. & Zhuang, X. Evaluation of fluorophores for optimal performance in localization-based super-resolution imaging. Nature methods 8, 1027–1036, doi:10.1038/nmeth.1768 (2011).

160 Jones, S. A., Shim, S. H., He, J. & Zhuang, X. Fast, three-dimensional super-resolution imaging of live cells. Nature methods 8, 499–508, doi:10.1038/nmeth.1605 (2011).

161 Wieckowski, M. R., Giorgi, C., Lebiedzinska, M., Duszynski, J. & Pinton, P. Isolation of mitochondria-associated membranes and mitochondria from animal tissues and cells. Nature protocols 4, 1582–1590, doi:10.1038/nprot.2009.151 (2009).

162 Neumann, D., Woods, A., Carling, D., Wallimann, T. & Schlattner, U. Mammalian AMP-activated protein kinase: functional, heterotrimeric complexes by co-expression of subunits in Escherichia coli. Protein expression and purification 30, 230–237 (2003).

163 Anderson, K. A. et al. Components of a calmodulin-dependent protein kinase cascade. Molecular cloning, functional characterization and cellular localization of Ca2+/calmodulin-dependent protein kinase kinase beta. The Journal of biological chemistry 273, 31880–31889, doi:10.1074/jbc.273.48.31880 (1998).

164 Chen, L. et al. Structural insight into the autoinhibition mechanism of AMP-activated protein kinase. Nature 459, 1146–1149, doi:10.1038/nature08075 (2009).

165 Woods, A. et al. Identification of phosphorylation sites in AMP-activated protein kinase (AMPK) for upstream AMPK kinases and study of their roles by site-directed mutagenesis. The Journal of biological chemistry 278, 28434–28442, doi:10.1074/jbc.M303946200 (2003).

166 Hawley, S. A. et al. Calmodulin-dependent protein kinase kinase-beta is an alternative upstream kinase for AMP-activated protein kinase. Cell Metab. 2, 9–19 (2005).

167 Davies, S. P., Carling, D. & Hardie, D. G. Tissue distribution of the AMP-activated protein kinase, and lack of activation by cyclic-AMP-dependent protein kinase, studied using a specific and sensitive peptide assay. Eur J Biochem 186, 123–128, doi:10.1111/j.1432-1033.1989.tb15185.x (1989).

168 Amrhein, V., Greenland, S. & McShane, B. Scientists rise up against statistical significance. Nature 567, 305–307, doi:10.1038/d41586-019-00857-9 (2019).

169 Wasserstein, R. L., Schirm, A. L. & Lazar, N. A. Moving to a World Beyond “p < 0.05”. The American Statistician 73, 1–19, doi:10.1080/00031305.2019.1583913 (2019).

170 Chen, T. et al. The Genome Sequence Archive Family: Toward Explosive Data Growth and Diverse Data Types. Genomics Proteomics Bioinformatics 19, 578–583, doi:10.1016/j.gpb.2021.08.001 (2021).

171 Members, C.-N. & Partners. Database Resources of the National Genomics Data Center, China National Center for Bioinformation in 2022. Nucleic Acids Res 50, D27–D38, doi:10.1093/nar/gkab951 (2022).

172 Ma, J. et al. iProX: an integrated proteome resource. Nucleic Acids Res 47, D1211–D1217, doi:10.1093/nar/gky869 (2019).

173 Chen, T. et al. iProX in 2021: connecting proteomics data sharing with big data. Nucleic Acids Res 50, D1522–D1527, doi:10.1093/nar/gkab1081 (2022).

## References for Supplementary Notes

174 Helle, S. C. et al. Organization and function of membrane contact sites. Biochimica et biophysica acta 1833, 2526–2541, doi:10.1016/j.bbamcr.2013.01.028 (2013).

175 Vallese, F. et al. An expanded palette of improved SPLICS reporters detects multiple organelle contacts in vitro and in vivo. Nat Commun 11, 6069, doi:10.1038/s41467-020-19892-6 (2020).

176 Owen, O. E., Kalhan, S. C. & Hanson, R. W. The key role of anaplerosis and cataplerosis for citric acid cycle function. The Journal of biological chemistry 277, 30409–30412, doi:10.1074/jbc.R200006200 (2002).

177 Strzelecki, T. & Schoolwerth, A. C. The significance of the attachment of rat kidney glutaminase to the inner mitochondrial membrane. Biochimica et biophysica acta 801, 334–341, doi:10.1016/0304-4165(84)90136-3 (1984).

178 Shapiro, R. A., Haser, W. G. & Curthoys, N. P. The orientation of phosphate-dependent glutaminase on the inner membrane of rat renal mitochondria. Archives of biochemistry and biophysics 243, 1–7, doi:10.1016/0003-9861(85)90767-2 (1985).

179 Kalra, J. & Brosnan, J. T. The subcellular localization of glutaminase isoenzymes in rat kidney cortex. The Journal of biological chemistry 249, 3255–3260 (1974).

180 Lam, S. S. et al. Directed evolution of APEX2 for electron microscopy and proximity labeling. Nature methods 12, 51–54, doi:10.1038/nmeth.3179 (2015).

181 Roberg, B., Torgner, I. A. & Kvamme, E. Inhibition of glutamine transport in rat brain mitochondria by some amino acids and tricarboxylic acid cycle intermediates. Neurochem Res 24, 809–814, doi:10.1023/a:1020941510764 (1999).

182 Curthoys, N. P. & Shapiro, R. A. Effect of metabolic acidosis and of phosphate on the presence of glutamine within the matrix space of rat renal mitochondria during glutamine transport. The Journal of biological chemistry 253, 63–68 (1978).

183 Shapiro, R. A., Morehouse, R. F. & Curthoys, N. P. Inhibition by glutamate of phosphate-dependent glutaminase of rat kidney. The Biochemical journal 207, 561–566, doi:10.1042/bj2070561 (1982).

184 Kornmann, B. et al. An ER-mitochondria tethering complex revealed by a synthetic biology screen. Science 325, 477–481, doi:10.1126/science.1175088 (2009).

185 Meisinger, C. et al. The mitochondrial morphology protein Mdm10 functions in assembly of the preprotein translocase of the outer membrane. Developmental cell 7, 61–71, doi:10.1016/j.devcel.2004.06.003 (2004).

