## Supplementary material for "AMPK targets PDZD8 to trigger carbon source shift to glutamine": Summary of lifespan and analysis in worms

**Supplementary Table 3 | Summary of lifespan and analysis in worms[a,b]**

| Genotypes/ treatments | Mean life span (days) | | | Median life span (days) | | | | | | P-value Vs saline control within each genotype (Mantel-CoX) |
|---|---|---|---|---|---|---|---|---|---|---|
| | | 95% confidence interval | | | 95% confidence interval | | | | | |
| | Estimated life span ± s.e.m. | Lower bound | Upper bound | Estimated life span ± s.e.m. | Lower bound | Upper bound | N[c] | N[d] | N[e] | |
| **Fig. 5d** | | | | | | | | | | |
| *Pdzd-8*$^{-/-}$ + PDZD8-WT | 18.194 ± 0.353 | 17.503 | 18.886 | 18.000 ± 0.430 | 17.158 | 18.842 | 185 | 15 | 200 | N/A |
| *Pdzd-8*$^{-/-}$ + PDZD8-WT + 2-DG | 23.000 ± 0.477 | 22.064 | 23.935 | 24.000 ± 0.629 | 22.768 | 25.232 | 188 | 12 | 200 | <0.001 |
| *Pdzd-8*$^{-/-}$ + PDZD8-T527A | 16.997 ± 0.399 | 16.215 | 17.779 | 18.000 ± 0.991 | 16.057 | 19.943 | 187 | 13 | 200 | 0.132 |
| *Pdzd-8*$^{-/-}$ + PDZD8-T527A + 2-DG | 15.568 ± 0.327 | 14.926 | 16.209 | 16.000 ± 0.490 | 15.039 | 16.961 | 184 | 16 | 200 | <0.001 |
| **Fig. 5e** | | | | | | | | | | |
| CA-aak2;*Pdzd-8*$^{-/-}$ + PDZD8-T527A | 19.293 ± 0.405 | 18.498 | 20.087 | 20.000 ± 0.585 | 18.853 | 21.147 | 180 | 20 | 200 | N/A |
| CA-aak2;*Pdzd-8*$^{-/-}$ + PDZD8-WT | 25.293 ± 0.512 | 24.289 | 26.296 | 28.000 ± 0.504 | 27.011 | 28.989 | 180 | 20 | 200 | <0.001 |
| **Fig. 5f** | | | | | | | | | | |
| N2 | 18.030 ± 0.376 | 17.294 | 18.766 | 18.000 ± 0.482 | 17.056 | 18.944 | 184 | 16 | 200 | N/A |
| N2 + 2-DG | 21.282 ± 0.469 | 20.363 | 22.202 | 22.000 ± 0.605 | 20.185 | 23.185 | 187 | 13 | 200 | <0.001 |
| glna-null | 17.710 ± 0.353 | 17.018 | 18.402 | 18.000 ± 0.447 | 17.124 | 18.876 | 189 | 11 | 200 | 0.383 |
| glna-null + 2DG | 18.084 ± 0.373 | 17.354 | 18.815 | 18.000 ± 0.482 | 17.056 | 18.944 | 188 | 12 | 200 | 0.920 |
| **Fig. 5g** | | | | | | | | | | |
| glna-null + KGA-WT | 18.816 ± 0.405 | 18.023 | 19.609 | 18.000 ± 0.684 | 16.659 | 19.341 | 177 | 23 | 200 | N/A |
| glna-null + KGA-WT + 2-DG | 21.750 ± 0.434 | 20.899 | 22.600 | 22.000 ± 0.454 | 21.111 | 22.889 | 188 | 20 | 200 | <0.001 |
| glna-null + KGA-33A | 18.330 ± 0.395 | 17.556 | 19.104 | 18.000 ± 0.776 | 16.480 | 19.520 | 187 | 13 | 200 | 0.309 |
| glna-null + KGA-33A + 2-DG | 18.502 ± 0.395 | 17.728 | 19.276 | 18.000 ± 0.795 | 16.442 | 19.558 | 186 | 14 | 200 | 0.487 |
| **Fig. 5j** | | | | | | | | | | |
| *Pdzd-8*$^{-/-}$ + PDZD8-WT | 18.763 ± 0.394 | 17.991 | 19.536 | 20.000 ± 0.735 | 18.559 | 21.411 | 186 | 14 | 200 | N/A |
| *Pdzd-8*$^{-/-}$ + PDZD8-WT + CR | 25.067 ± 0.526 | 24.036 | 26.099 | 26.000 ± 0.548 | 24.925 | 27.075 | 184 | 16 | 200 | <0.001 |
| *Pdzd-8*$^{-/-}$ + PDZD8-T527A | 19.114 ± 0.391 | 18.348 | 19.880 | 20.000 ± 0.559 | 18.904 | 21.096 | 179 | 21 | 200 | 0.676 |
| *Pdzd-8*$^{-/-}$ + PDZD8-T527A + CR | 16.428 ± 0.277 | 15.885 | 16.971 | 18.000 ± 0.362 | 17.290 | 18.710 | 184 | 16 | 200 | <0.001 |
| **Fig. 5k** | | | | | | | | | | |
| N2 | 18.215 ± 0.356 | 17.517 | 18.912 | 18.000 ± 0.440 | 17.137 | 18.863 | 187 | 13 | 200 | N/A |
| N2 + CR | 21.786 ± 0.437 | 20.931 | 22.642 | 22.000 ± 0.625 | 20.774 | 23.226 | 186 | 14 | 200 | <0.001 |
| glna-null | 18.229 ± 0.365 | 17.514 | 18.943 | 18.000 ± 0.448 | 17.121 | 18.879 | 185 | 15 | 200 | 0.894 |
| glna-null + CR | 18.415 ± 0.377 | 17.676 | 19.154 | 18.000 ± 0.514 | 16.993 | 19.007 | 181 | 19 | 200 | 0.895 |
| **Fig. 5l** | | | | | | | | | | |
| glna-null + KGA-WT | 19.414 ± 0.406 | 18.619 | 20.210 | 20.000 ± 0.469 | 19.081 | 20.919 | 179 | 21 | 200 | N/A |
| glna-null + KGA-WT + CR | 25.293 ± 0.5212 | 24.289 | 26.296 | 26.000 ± 0.462 | 27.011 | 26.905 | 178 | 22 | 200 | <0.001 |
| glna-null + KGA-33A | 18.971 ± 0.409 | 18.169 | 19.772 | 20.000 ± 0.525 | 18.971 | 21.029 | 182 | 18 | 200 | 0.442 |
| glna-null + KGA-33A + CR | 19.140 ± 0.380 | 18.396 | 19.885 | 20.000 ± 0.521 | 18.978 | 21.022 | 179 | 21 | 200 | 0.404 |

**Supplementary Table 3 | Summary of lifespan and analysis in worms (cont.)[a,b]**

| Genotypes/ treatments | Mean life span (hours) | | | Median life span (hours) | | | | | | |
|---|---|---|---|---|---|---|---|---|---|---|
| | | 95% confidence interval | | | 95% confidence interval | | | | | P-value Vs saline control within each genotype (Mantel-CoX) |
| | Estimated life span ± s.e.m. | Lower bound | Upper bound | Estimated life span ± s.e.m. | Lower bound | Upper bound | N[c] | N[d] | N[e] | |
| **Fig. 5p** | | | | | | | | | | |
| *Pdzd-8*$^{-/-}$ + PDZD8-WT | 10.231 ± 0.558 | 9.138 | 11.324 | 11.000 ± 0.929 | 9.179 | 12.821 | 51 | 9 | 60 | N/A |
| *Pdzd-8*$^{-/-}$ + PDZD8-WT + CR | 14.878 ± 0.663 | 13.578 | 16.177 | 16.000 ± 0.779 | 14.474 | 17.526 | 47 | 13 | 60 | <0.001 |
| *Pdzd-8*$^{-/-}$ + PDZD8-T527A | 10.268 ± 0.493 | 9.302 | 11.234 | 11.000 ± 0.782 | 9.467 | 12.533 | 54 | 6 | 60 | 0.697 |
| *Pdzd-8*$^{-/-}$ + PDZD8-T527A + CR | 10.947 ± 0.487 | 9.992 | 11.901 | 12.000 ± 0.551 | 10.920 | 13.080 | 47 | 13 | 60 | 0.829 |
| **Fig. 5r** | | | | | | | | | | |
| glna-null + KGA-WT | 10.485 ± 0.513 | 9.480 | 11.490 | 11.000 ± 0.924 | 9.189 | 12.811 | 47 | 13 | 60 | N/A |
| glna-null + KGA-WT + CR | 14.233 ± 0.722 | 12.818 | 15.647 | 15.000 ± 0.867 | 13.301 | 16.699 | 49 | 11 | 60 | <0.001 |
| glna-null + KGA-33A | 10.804 ± 0.429 | 9.964 | 11.644 | 11.000 ± 0.575 | 9.874 | 12.126 | 48 | 12 | 60 | 0.811 |
| glna-null + KGA-33A + CR | 10.631 ± 0.491 | 9.668 | 11.595 | 11.000 ± 0.564 | 9.894 | 12.106 | 48 | 12 | 60 | 0.969 |
| **Extended Data Fig. 9h** | | | | | | | | | | |
| *Pdzd-8*$^{-/-}$ + PDZD8-WT | 9.584 ± 0.561 | 8.485 | 10.683 | 10.000 ± 0.859 | 8.316 | 11.684 | 49 | 11 | 60 | N/A |
| *Pdzd-8*$^{-/-}$ + PDZD8-WT + 2-DG | 13.767 ± 0.762 | 12.274 | 15.260 | 15.000 ± 0.758 | 13.514 | 16.486 | 50 | 10 | 60 | <0.001 |
| *Pdzd-8*$^{-/-}$ + PDZD8-T527A | 10.565 ± 0.464 | 9.654 | 11.475 | 11.000 ± 0.627 | 9.771 | 12.229 | 49 | 11 | 60 | 0.576 |
| *Pdzd-8*$^{-/-}$ + PDZD8-T527A + 2-DG | 10.407 ± 0.523 | 9.381 | 11.432 | 11.000 ± 0.732 | 9.565 | 12.435 | 48 | 12 | 60 | 0.430 |
| **Extended Data Fig. 9i** | | | | | | | | | | |
| glna-null + KGA-WT | 10.532 ± 0.510 | 9.533 | 11.531 | 11.000 ± 0.972 | 9.095 | 12.905 | 48 | 12 | 60 | N/A |
| glna-null + KGA-WT + 2-DG | 10.867 ± 0.460 | 13.356 | 16.067 | 16.000 ± 0.842 | 20.774 | 17.651 | 47 | 13 | 60 | <0.001 |
| glna-null + KGA-33A | 11.494 ± 0.453 | 10.605 | 12.383 | 12.000 ± 0.551 | 10.919 | 13.081 | 45 | 15 | 60 | 0.651 |
| glna-null + KGA-33A + 2-DG | 10.867 ± 0.460 | 9.965 | 11.770 | 11.000 ± 0.518 | 9.984 | 12.016 | 45 | 15 | 60 | 0.024 |

[a]Independent repeats of each health span experiment were performed. Data from representative experiments are shown.
[b]Health span data sets within each panel of this table were done in parallel and statistical analyses was done within the data set.
[c]Number of worms scored (death events).
[d]Number of worms censored.
[e]Total number of worms.
