## Supplementary material for "AMPK targets PDZD8 to trigger carbon source shift to glutamine": Uncropped gel images

**Supplementary Fig. 1 | Uncropped gels.**

Blots, as shown in this figure, were cut into slices before incubation with primary antibodies. The Pierce™ Prestained Protein MW Marker, Cat. 26612, from ThermoFisher Scientific, was used as the protein marker.

**Fig. 1a**

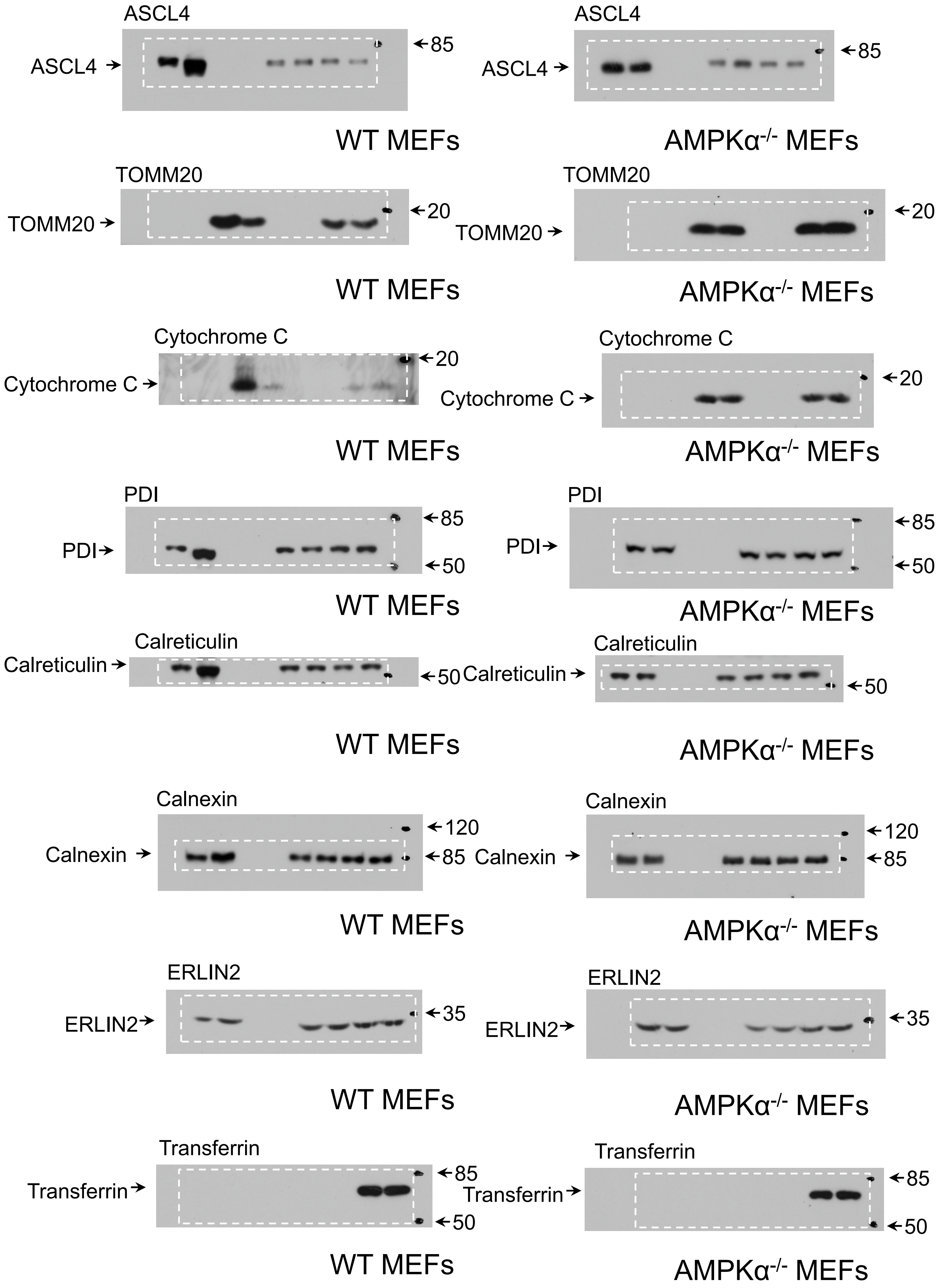

ASCL4
ASCL4
85
WT MEFs
ASCL4
ASCL4
85
AMPKα-/- MEFs
TOMM20
TOMM20
20
WT MEFs
TOMM20
TOMM20
20
AMPKα-/- MEFs
Cytochrome C
Cytochrome C
20
WT MEFs
Cytochrome C
Cytochrome C
20
AMPKα-/- MEFs
PDI
PDI
85
50
WT MEFs
PDI
PDI
85
50
AMPKα-/- MEFs
Calreticulin
Calreticulin
50
WT MEFs
Calreticulin
Calreticulin
50
AMPKα-/- MEFs
Calnexin
Calnexin
120
85
WT MEFs
Calnexin
Calnexin
120
85
AMPKα-/- MEFs
ERLIN2
ERLIN2
35
WT MEFs
ERLIN2
ERLIN2
35
AMPKα-/- MEFs
Transferrin
Transferrin
85
50
WT MEFs
Transferrin
Transferrin
85
50
AMPKα-/- MEFs

#### Fig. 1h

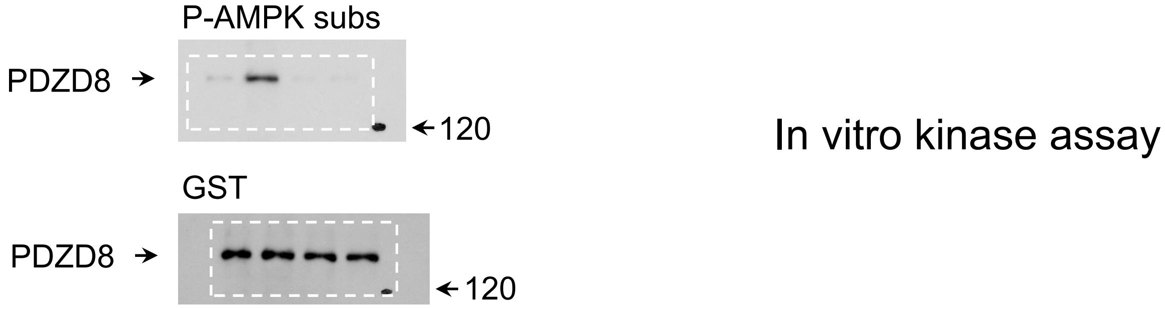


#### Fig. 1i

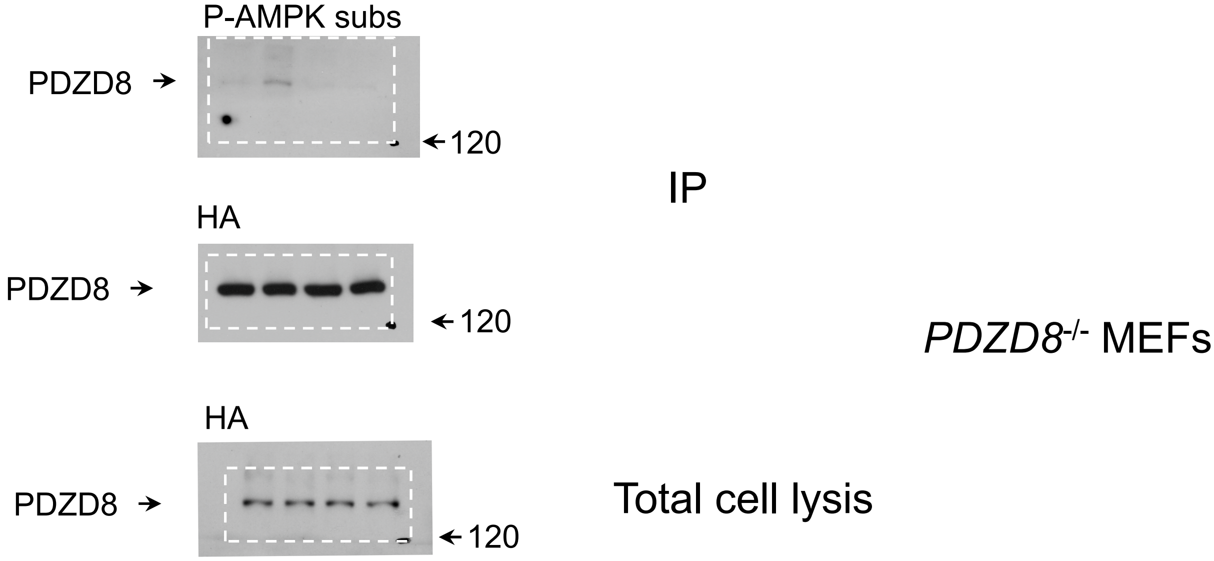


#### Fig. 1j

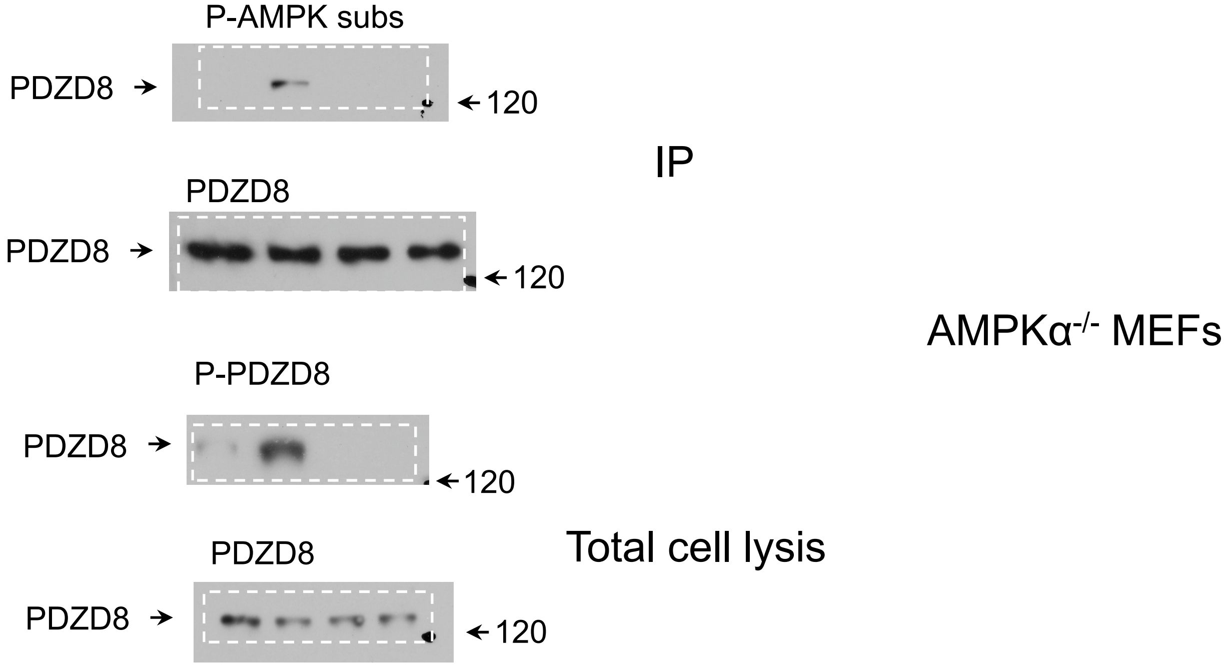

#### Fig. 1k

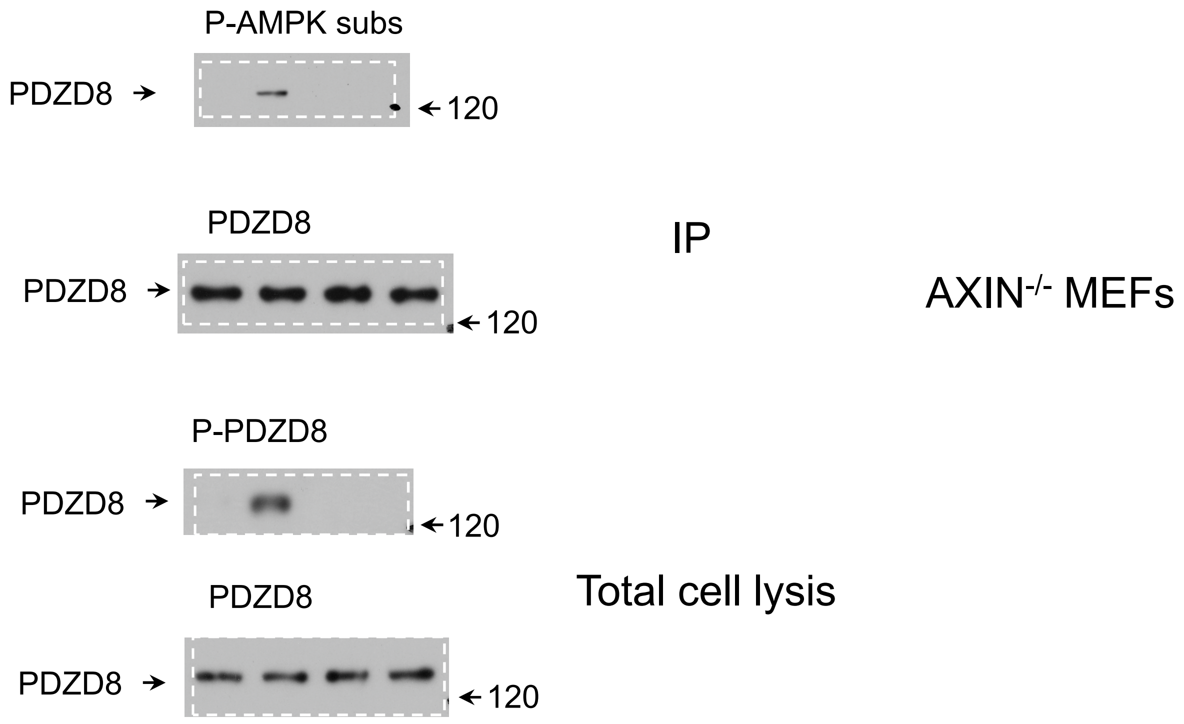
P-AMPK subs
PDZD8
120
PDZD8
IP
AXIN-/- MEFs
PDZD8
120
P-PDZD8
PDZD8
120
PDZD8
Total cell lysis
PDZD8
120

#### Fig. 1l

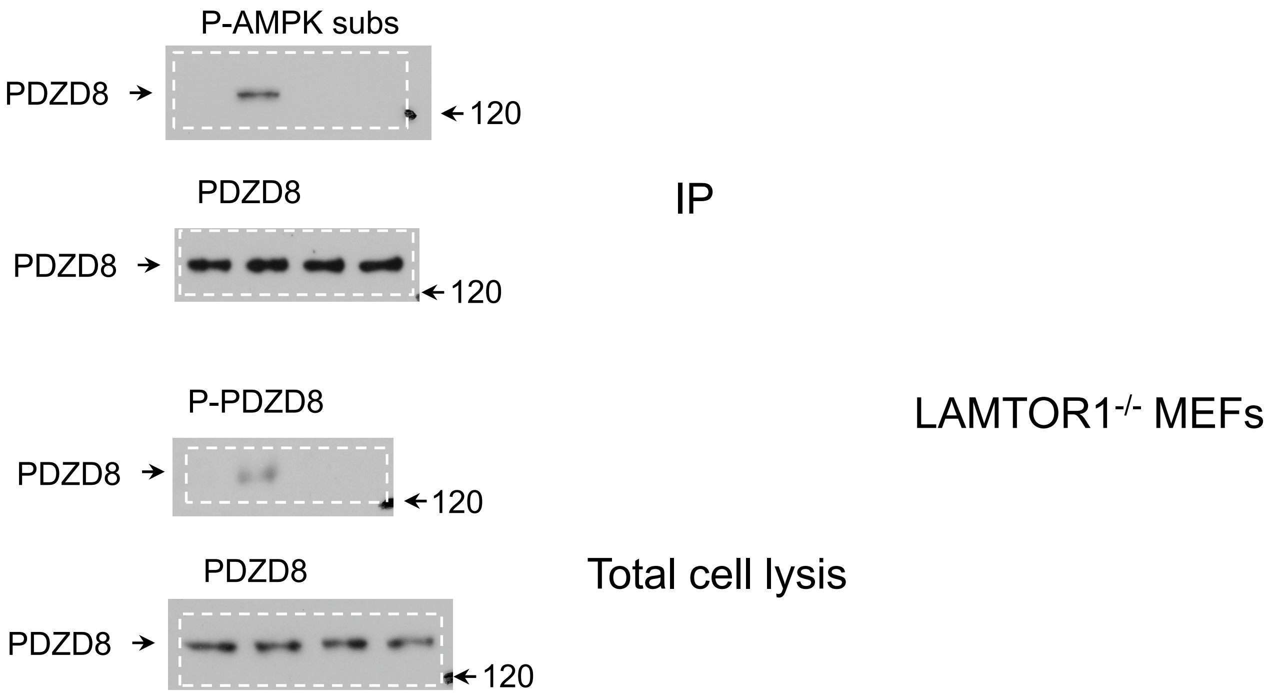
P-AMPK subs
PDZD8
120
PDZD8
IP
PDZD8
120
P-PDZD8
LAMTOR1-/- MEFs
PDZD8
120
PDZD8
Total cell lysis
PDZD8
120

### Fig. 2r

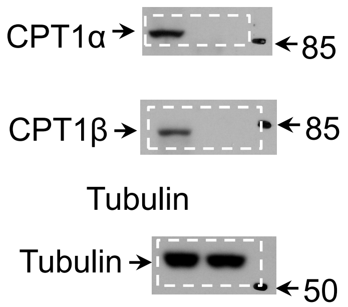

CPT1α
85
CPT1β
85
Tubulin
Tubulin
50


### Fig. 3h

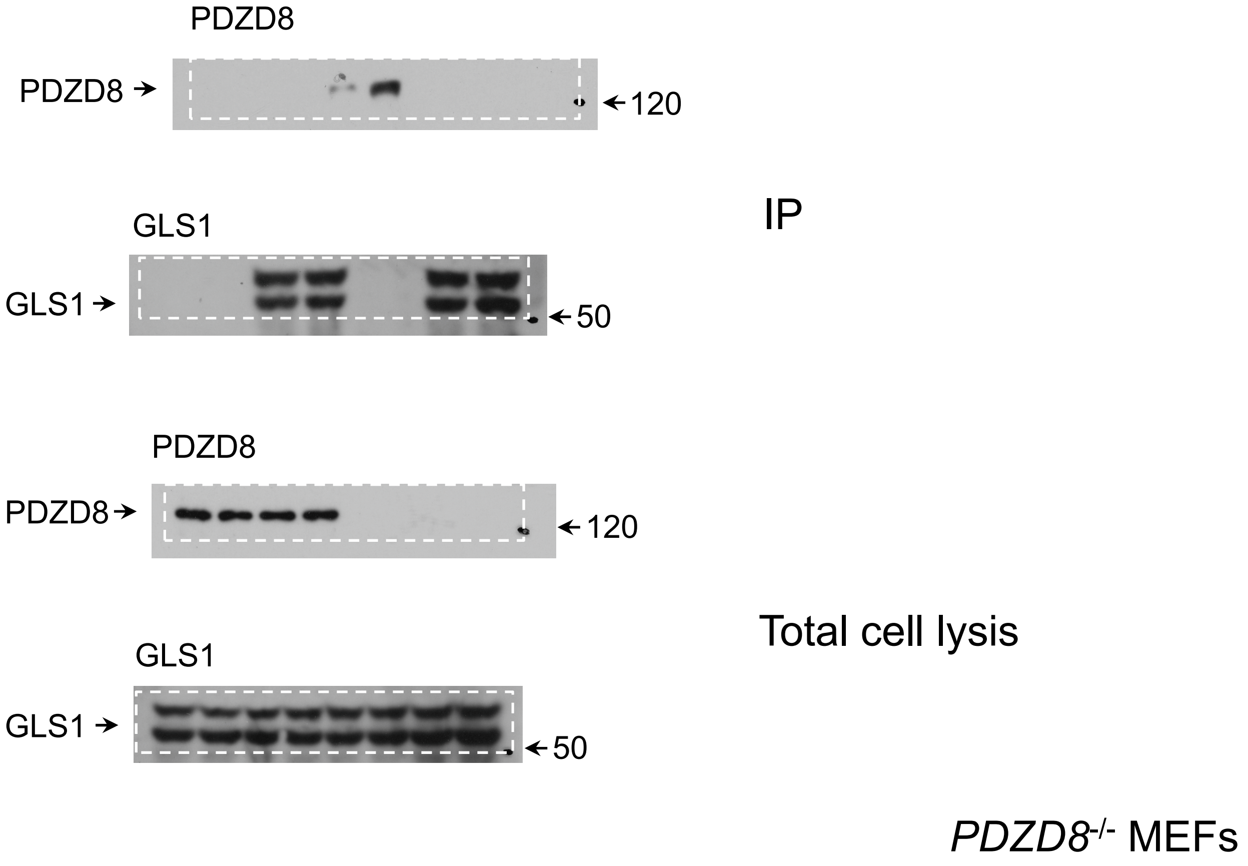

PDZD8
PDZD8
120
IP
GLS1
GLS1
50
PDZD8
PDZD8
120
Total cell lysis
GLS1
GLS1
50
*PDZD8*-/- MEFs

#### Fig. 3l

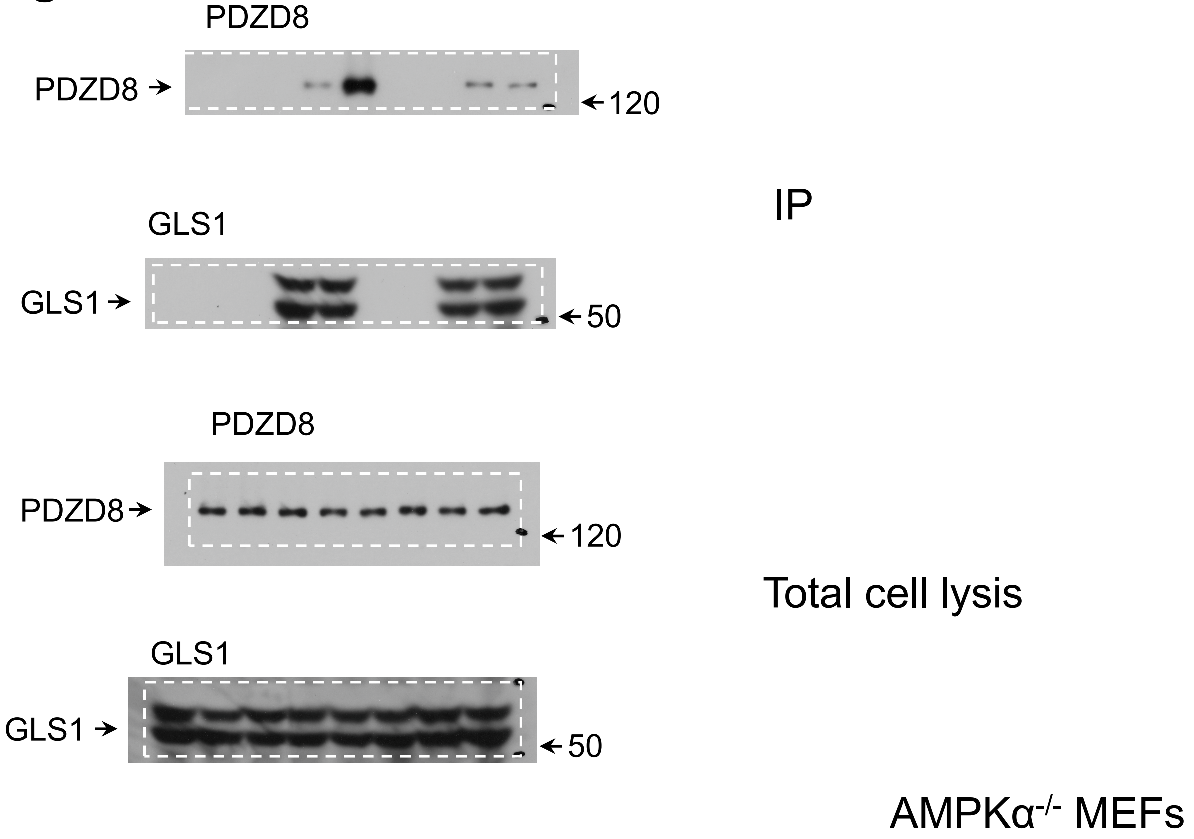

PDZD8
PDZD8
120
IP
GLS1
GLS1
50
PDZD8
PDZD8
120
Total cell lysis
GLS1
GLS1
50
AMPKα-/- MEFs


#### Fig. 3m

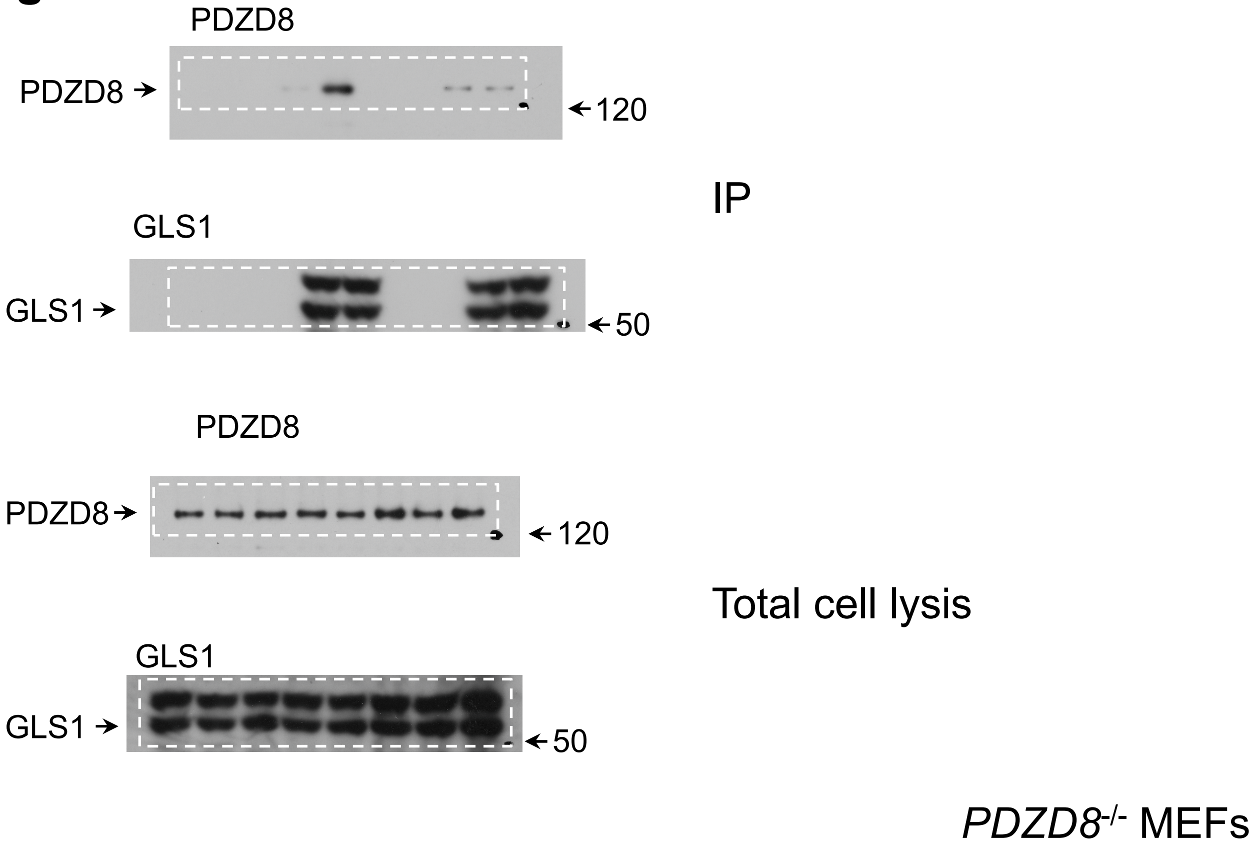

PDZD8
PDZD8
120
IP
GLS1
GLS1
50
PDZD8
PDZD8
120
Total cell lysis
GLS1
GLS1
50
PDZD8-/- MEFs

### Fig. 3p

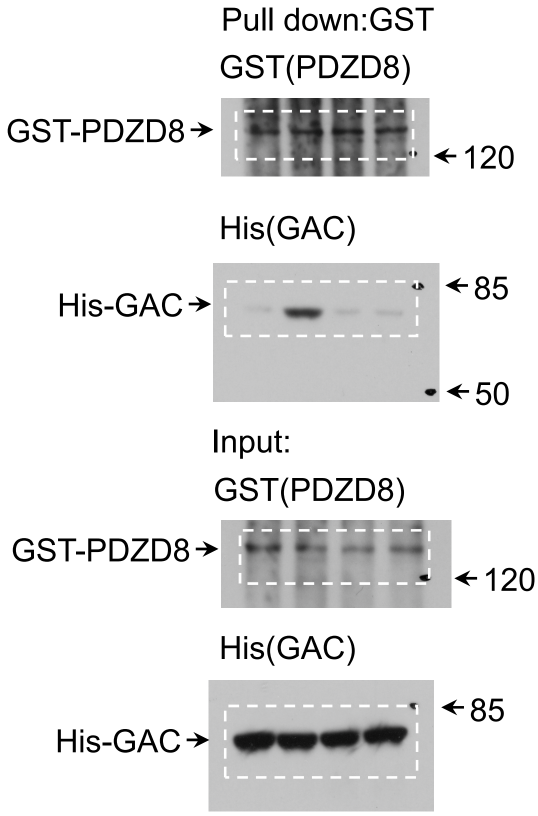


### Fig. 3p

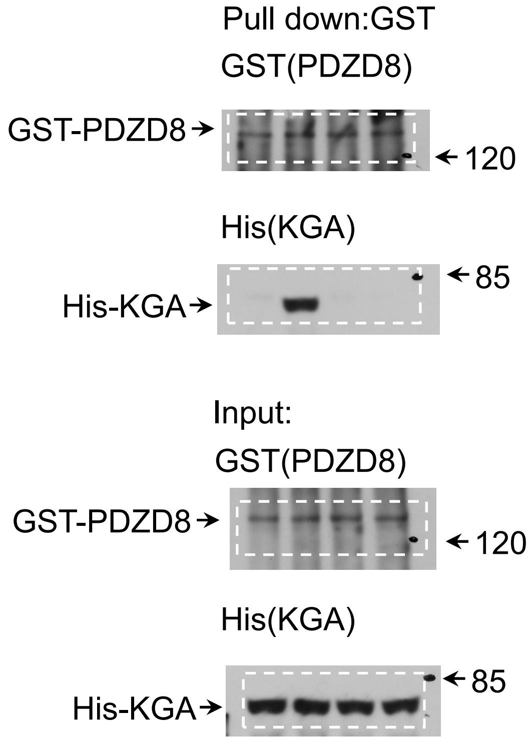


### Fig. 4i

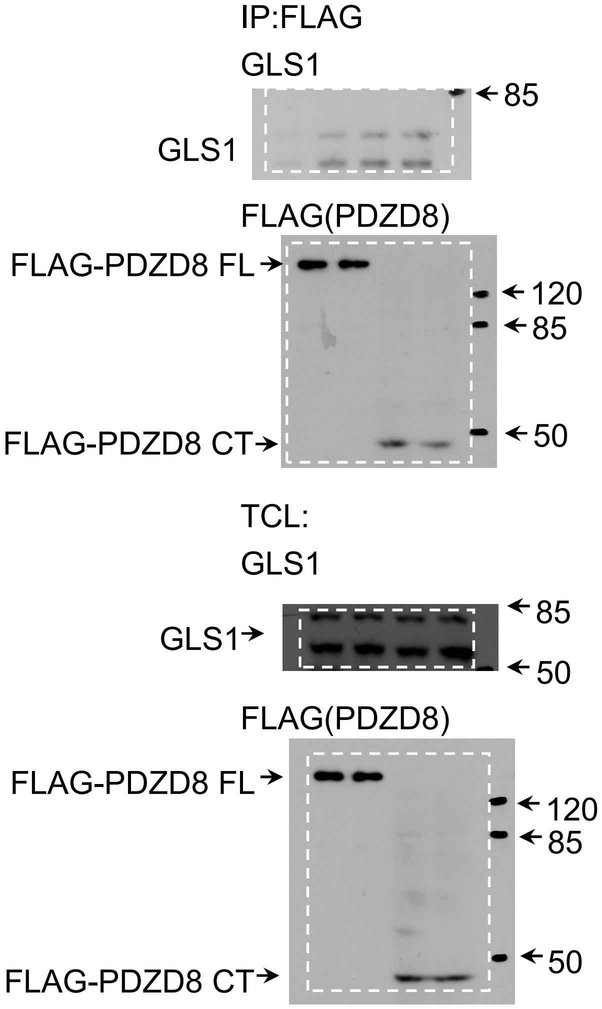

### Extended Data fig. 2a-1

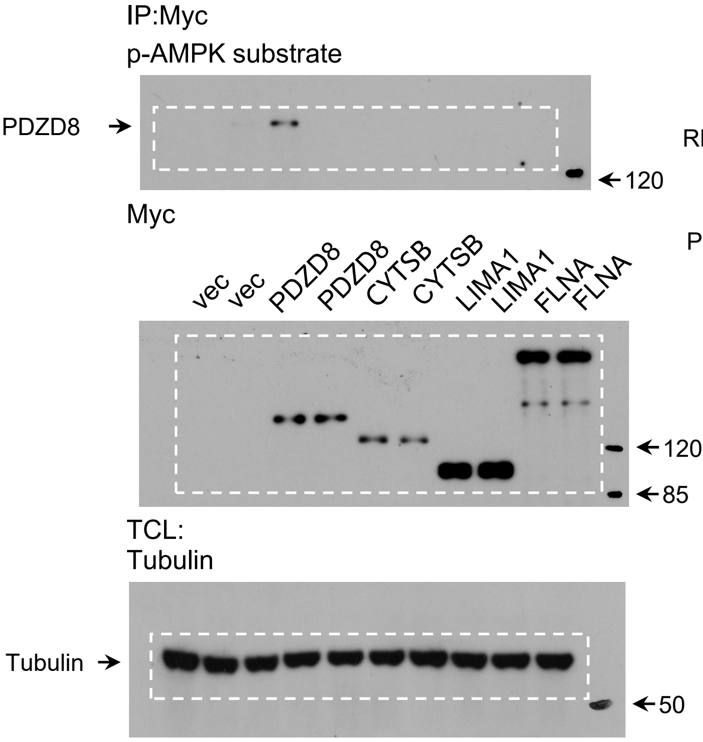


### Extended Data fig. 2a-2

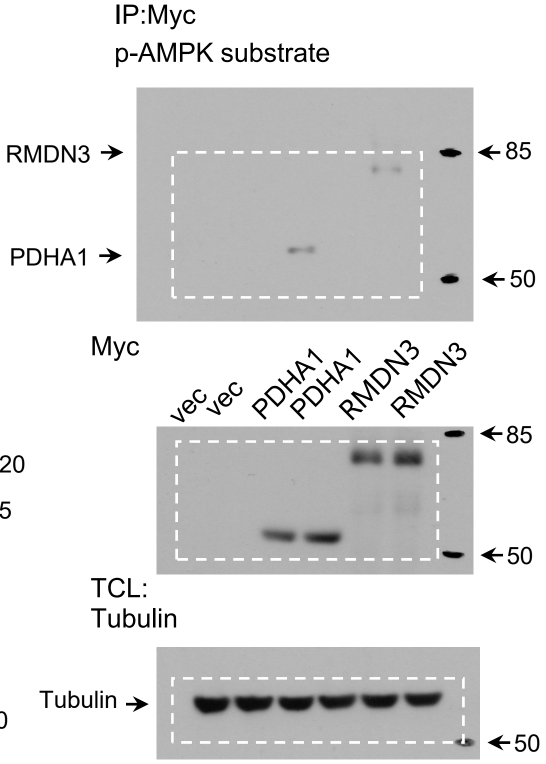


### Extended Data fig. 2a-3

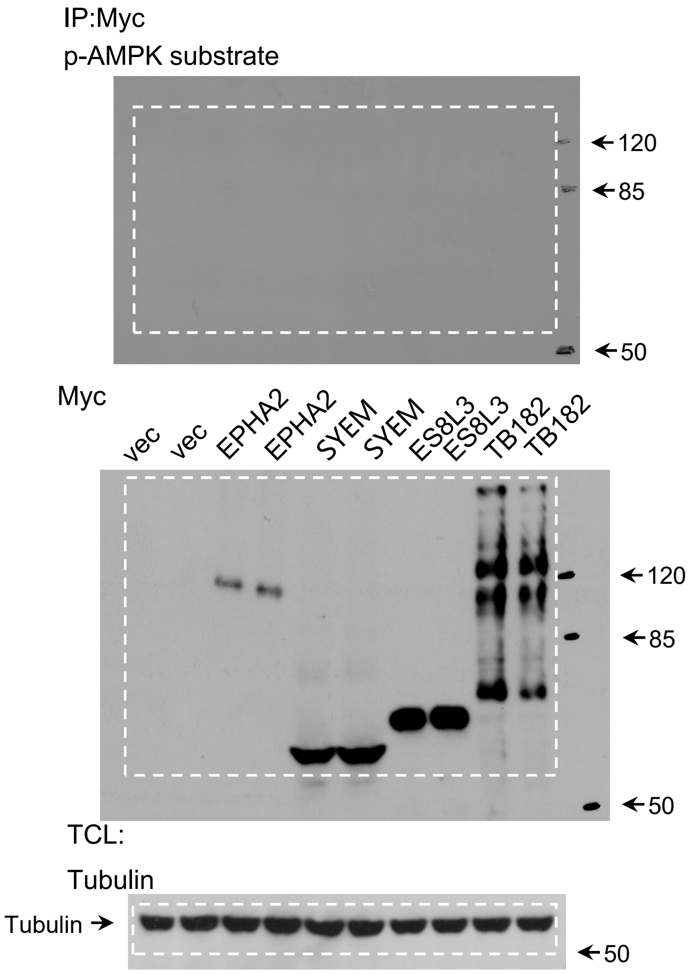


### Extended Data fig. 2a-4

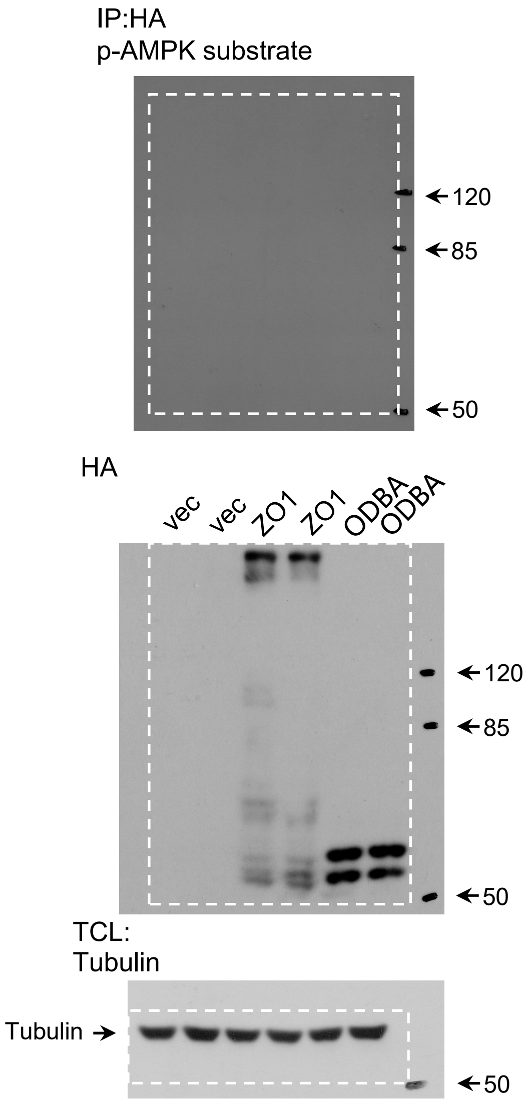

#### Extended Data fig. 2a-5

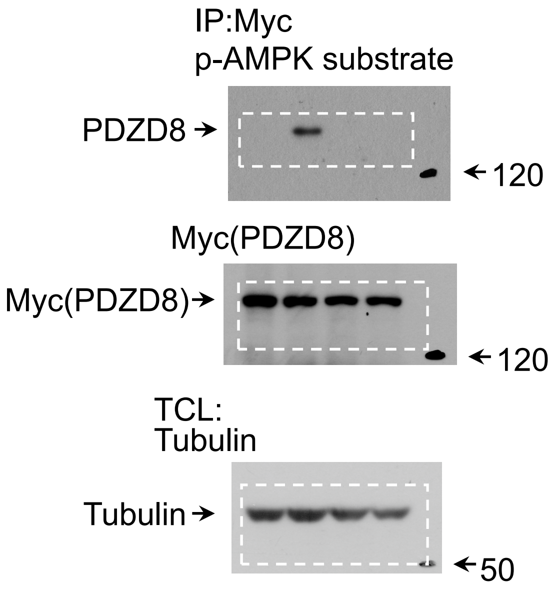


#### Extended Data fig. 2a-6

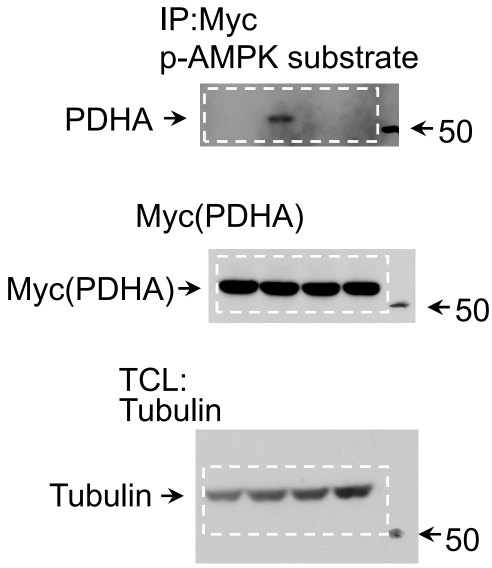


#### Extended Data fig. 2a-7

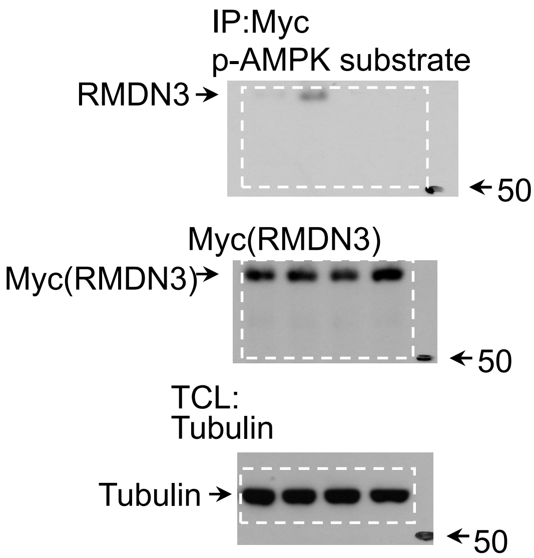


#### Extended Data fig. 2b

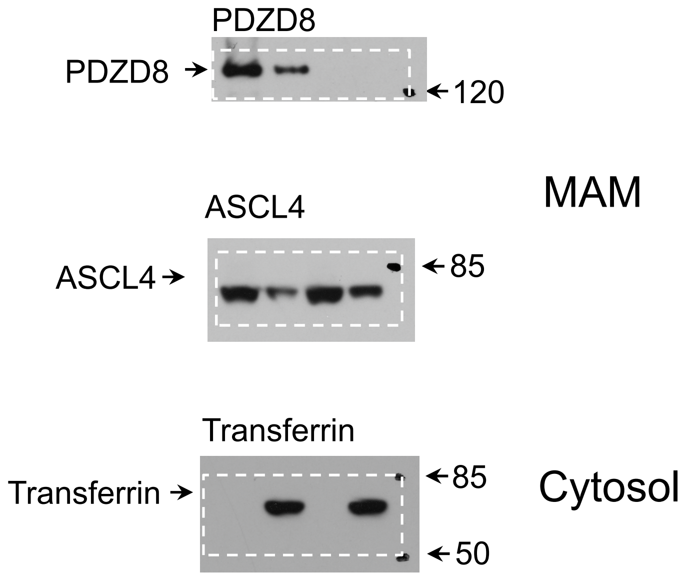

#### Extended Data fig. 2g Left

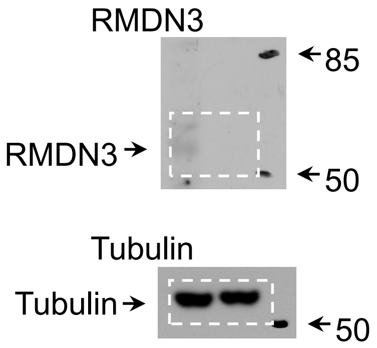


#### Extended Data fig. 2g Right

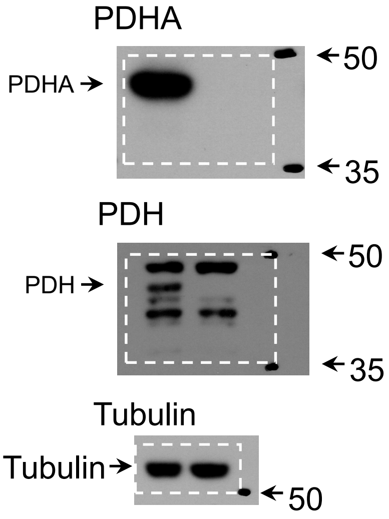


#### Extended Data Fig. 3b

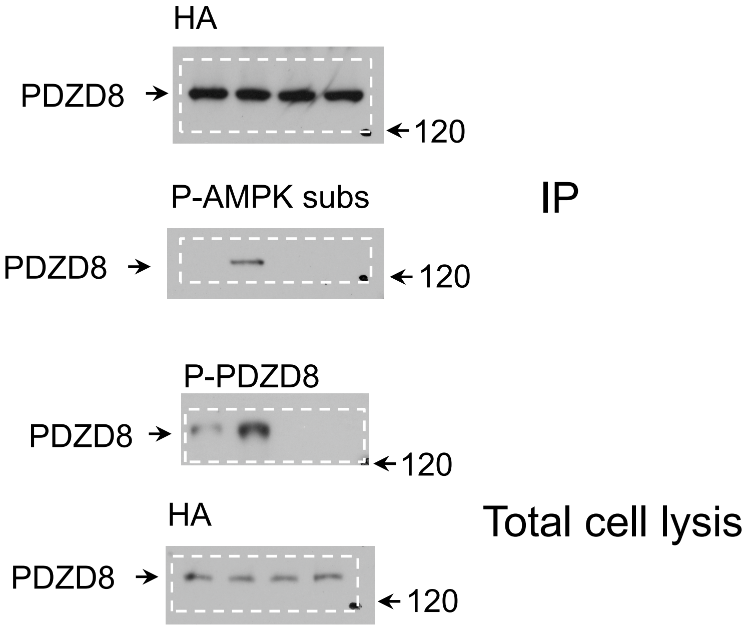


#### Extended Data fig. 4a

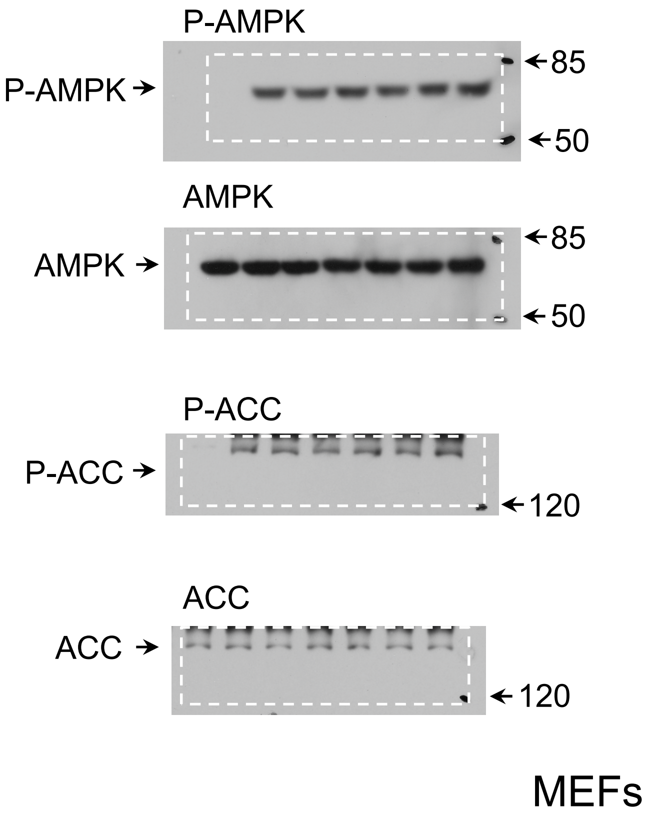

### Extended Data fig. 5c

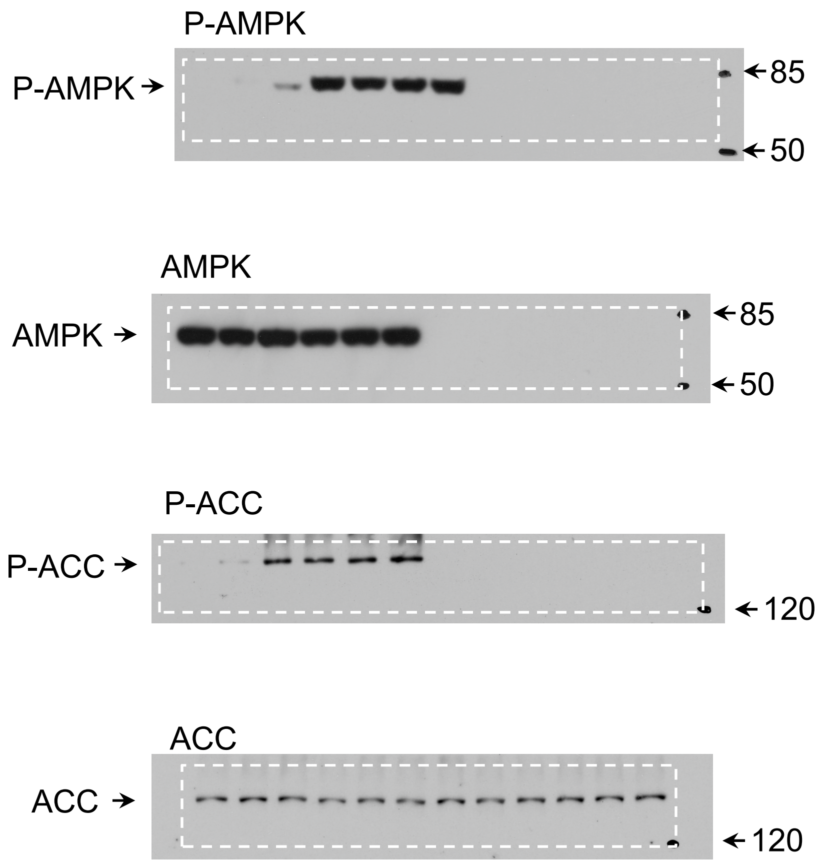


Muscle

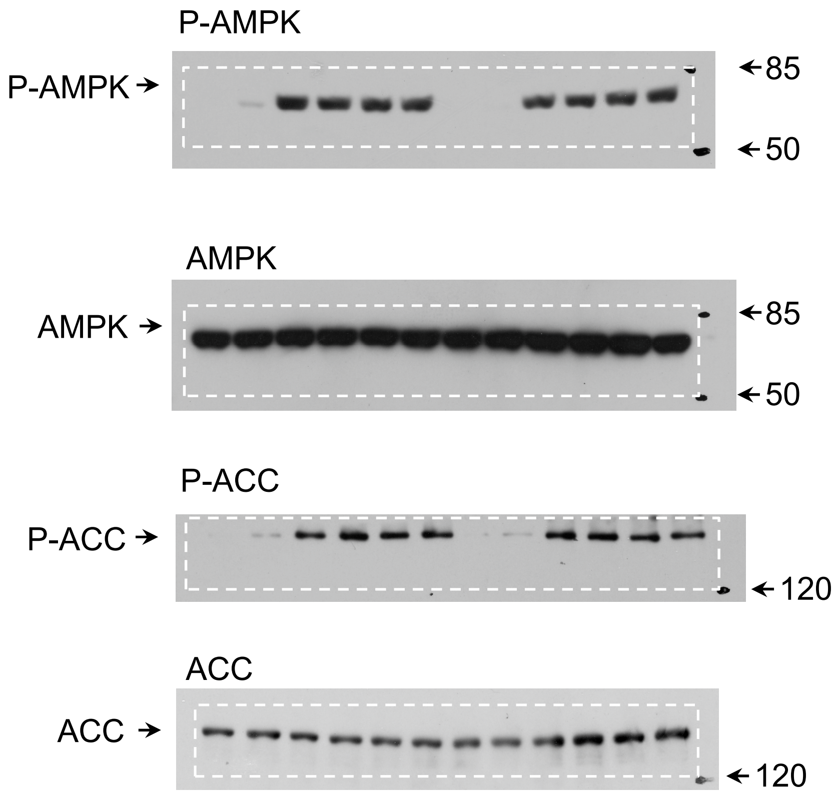


Liver

#### Extended Data Fig. 5i

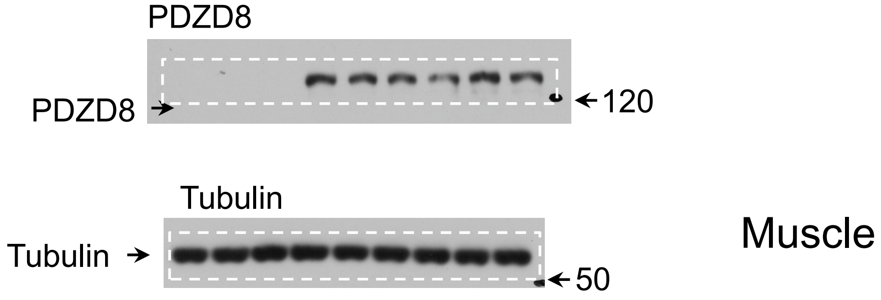


#### Extended Data Fig. 5n

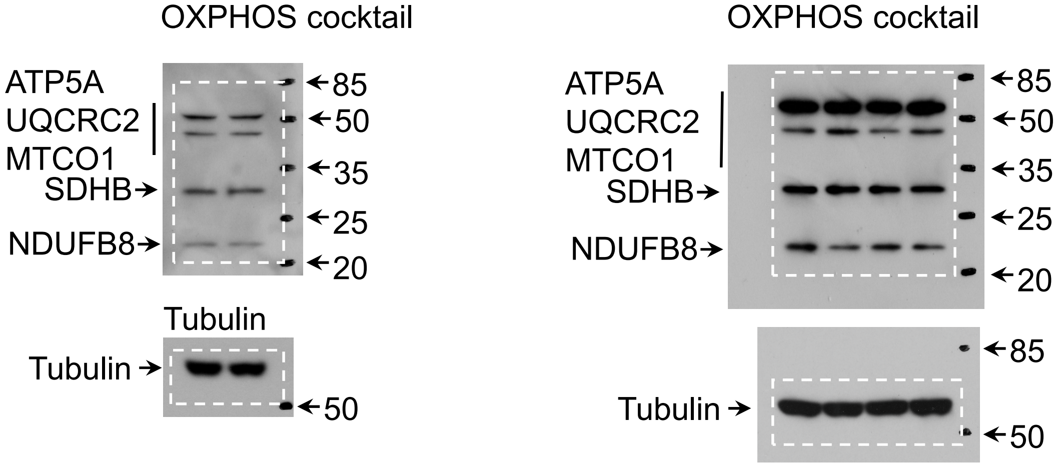

### Extended Data Fig. 6b

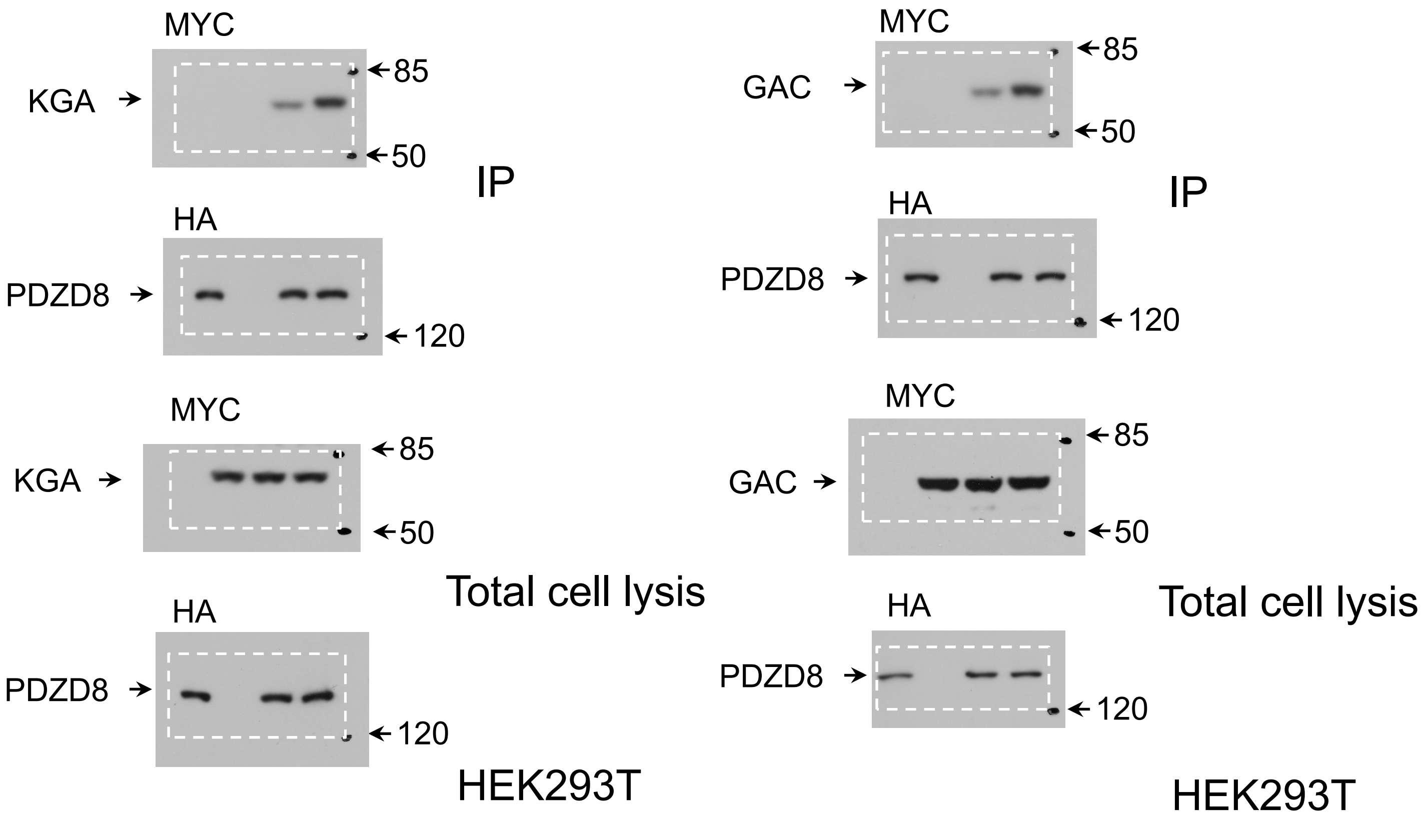


### Extended Data Fig. 6c

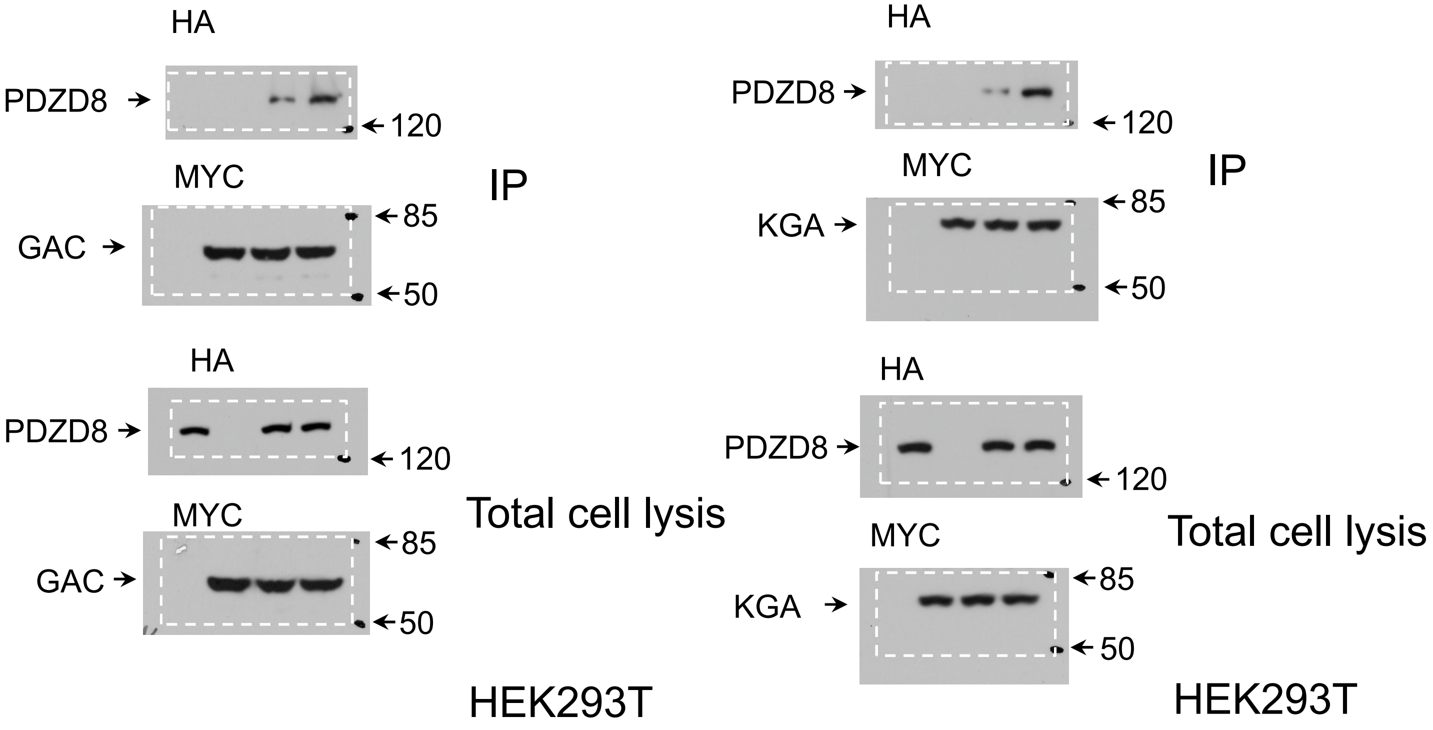

#### Extended Data Fig. 7a left

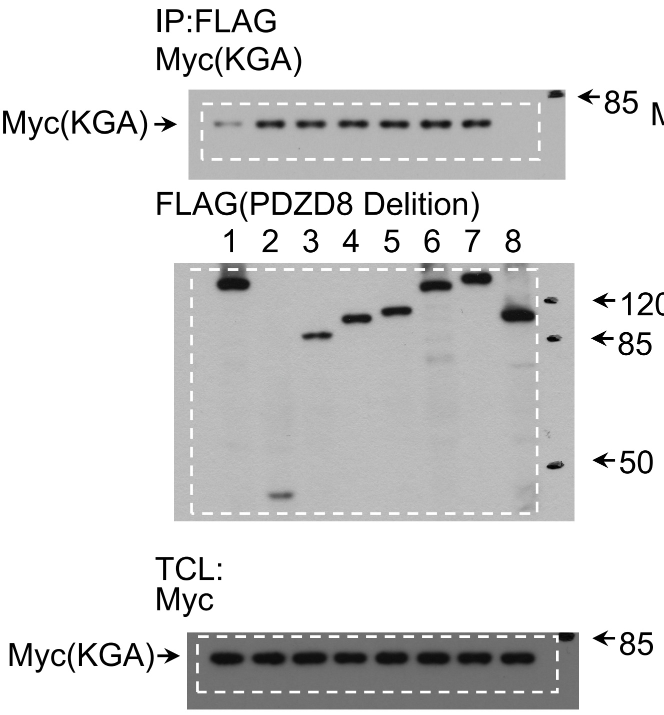


#### Extended Data Fig. 7a right

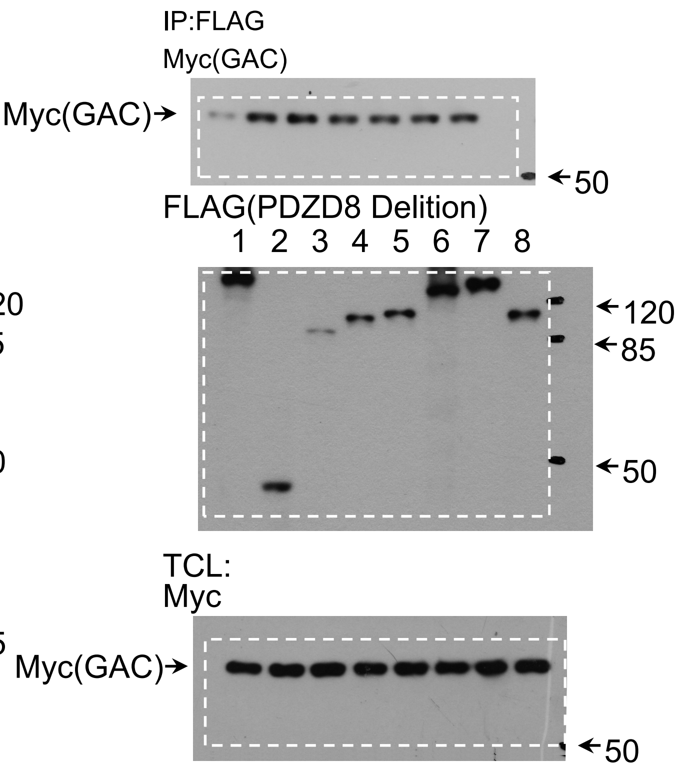


#### Extended Data Fig. 8a left

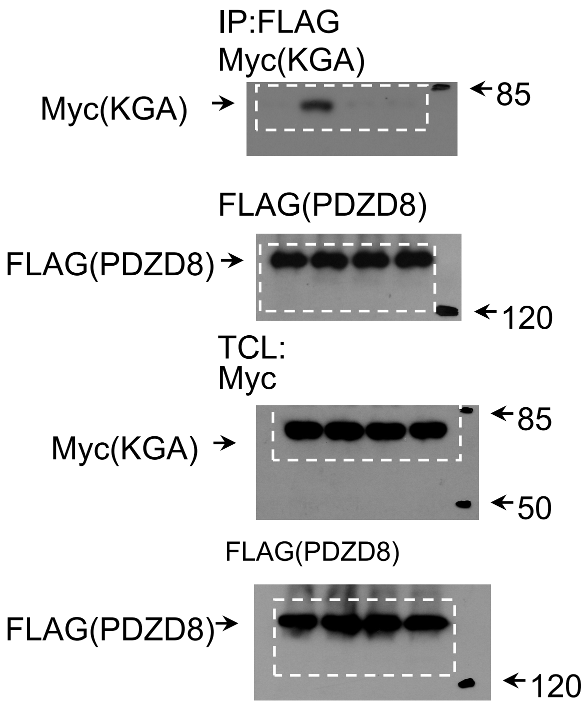


#### Extended Data Fig. 8a right

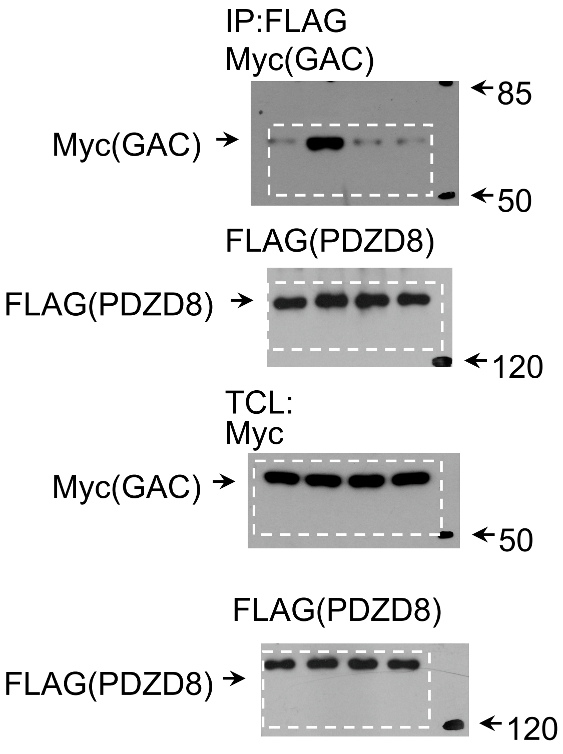

#### Extended Data Fig. 8c top

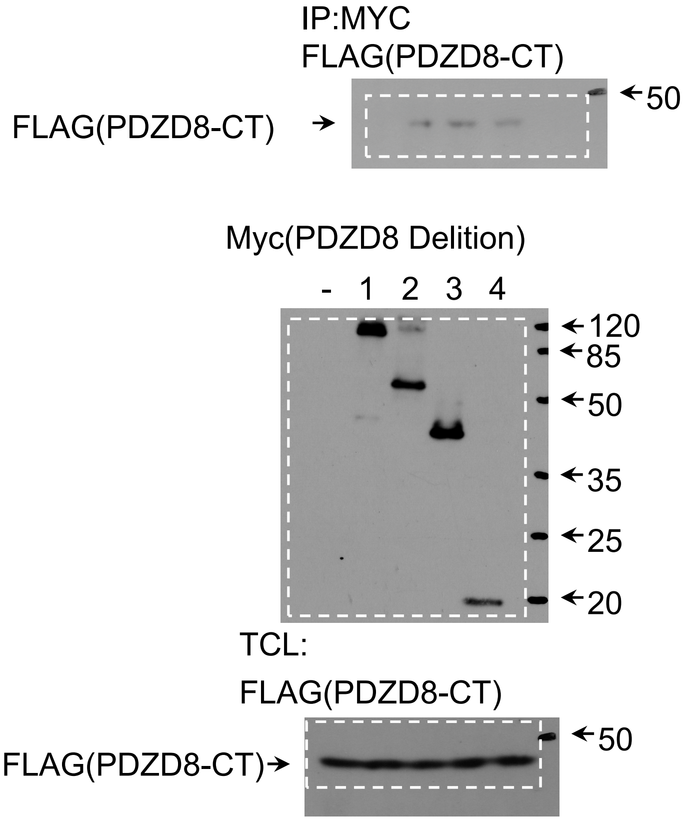


#### Extended Data Fig. 8c middle

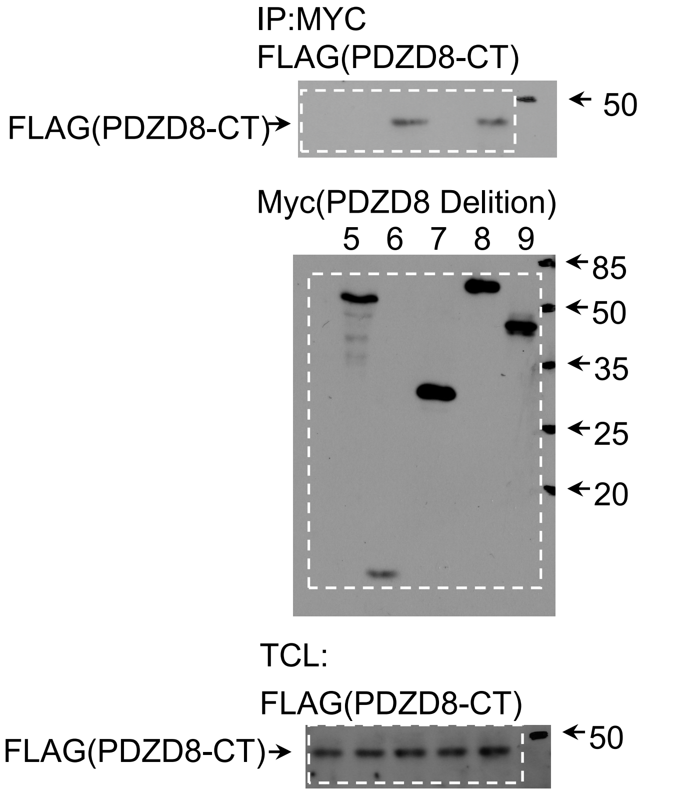


#### Extended Data Fig. 8c bottom

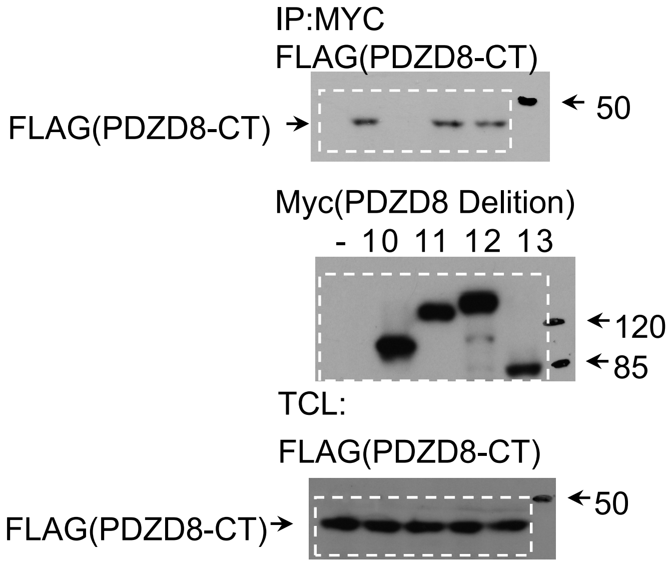

#### Extended Data Fig. 9a

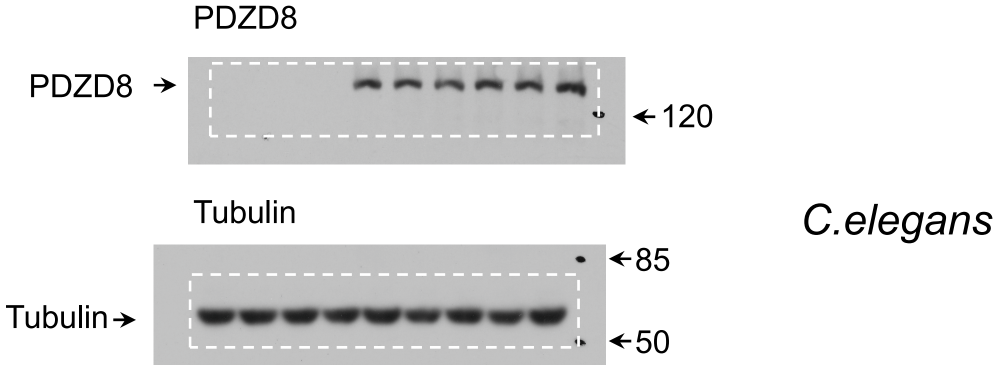


#### Extended Data Fig. 10a

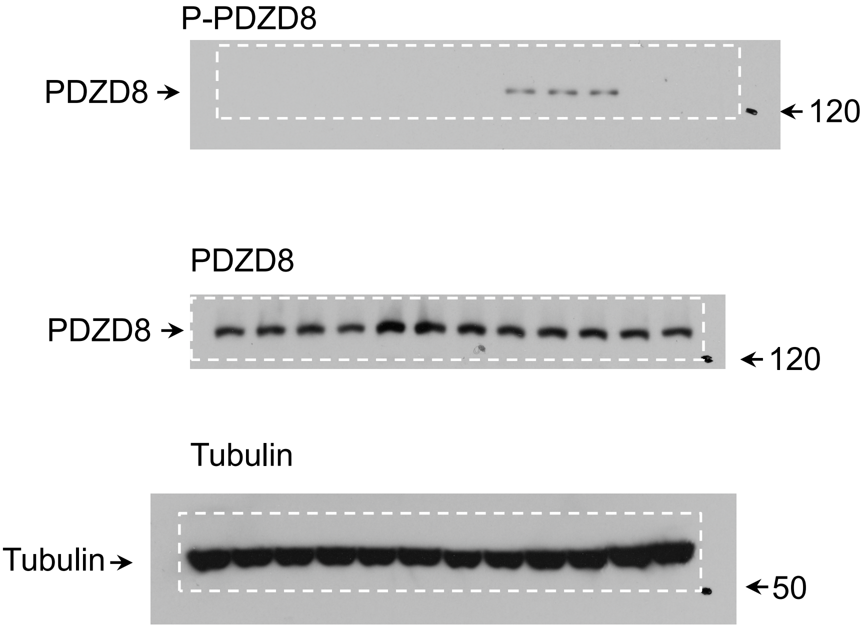

### Extended Data Fig. 10e

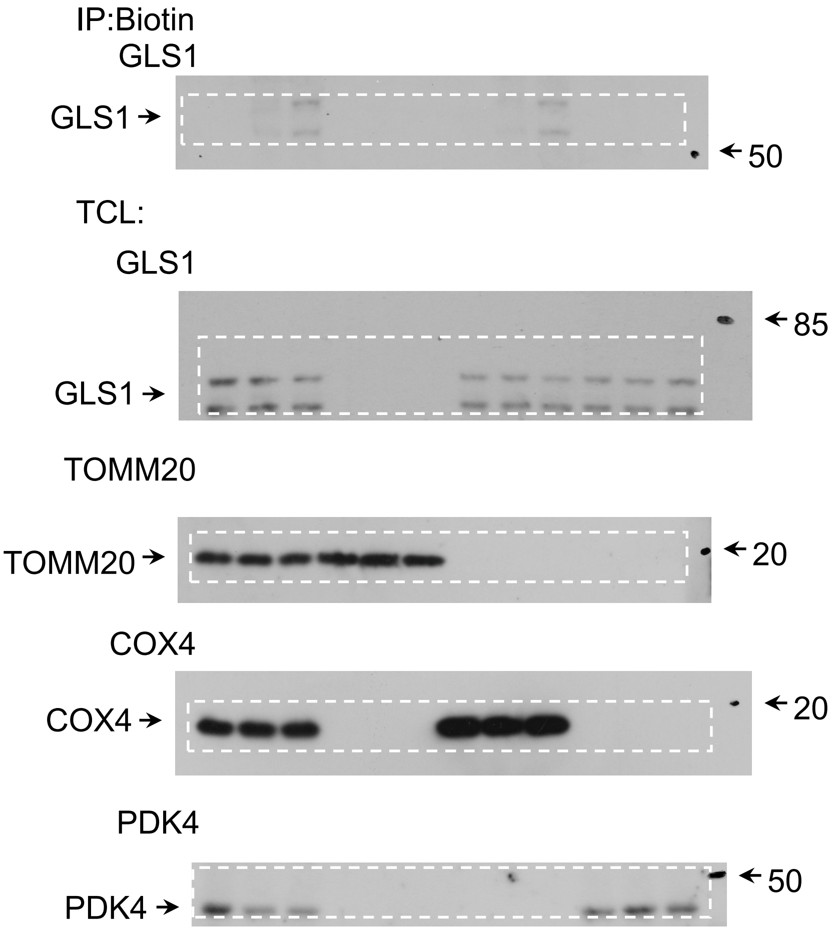


### Extended Data Fig. 10e

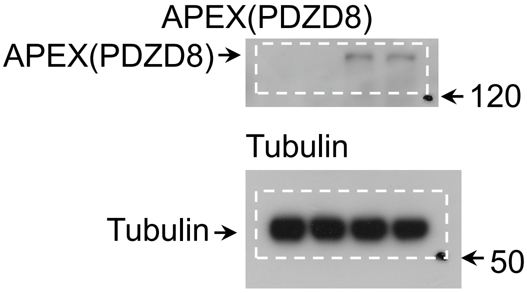
